# Trans-Omics analysis of post injury thrombo-inflammation identifies endotypes and trajectories in trauma patients

**DOI:** 10.1101/2023.08.16.553446

**Authors:** Mitchell J. Cohen, Christopher B. Erickson, Ian S. Lacroix, Margot Debot, Monika Dzieciatkowska, Sanchayita Mitra, Terry R. Schaid, William M. Hallas, Otto N. Thielen, Alexis L. Cralley, Anirban Banerjee, Ernest E Moore, Christopher C. Silliman, Angelo D’Alessandro, Kirk C. Hansen

**Affiliations:** Department of Surgery, University of Colorado Anschutz Medical Campus; Aurora, CO; Department of Biochemistry and Molecular Genetics, University of Colorado Anschutz Medical Campus; Aurora, CO; Denver Health Medical Center; Denver, CO; Department of Pediatrics, University of Colorado Anschutz Medical Campus; Aurora, CO; Vitalant Research Institute; Denver, CO; Center for Cancer and Blood Disorders, Children’s Hospital of Colorado; Aurora, CO

## Abstract

Understanding and managing the complexity of trauma-induced thrombo-inflammation necessitates an innovative, data-driven approach. This study leveraged a trans-omics analysis of longitudinal samples from trauma patients to illuminate molecular endotypes and trajectories that underpin patient outcomes. We hypothesized that trans-omics profiling reveals underlying clinical differences in severely injured patients that may present with similar clinical characteristics but ultimately have different responses to treatment and outcomes. Here we used proteomics and metabolomics to profile 759 of longitudinal plasma samples from 118 patients at 11 time points and 97 control subjects. Patients were stratified by shock and injury severity, revealing a spectrum of responses to trauma and treatment that are fundamentally tied to their unique underlying biology. Ensemble models were employed, demonstrating the predictive power of these molecular signatures with area under the receiver operating curves of 80 to 94% for key outcomes such as death. Then, transomics-based patient states were defined to create a map of unique pathophysiologic states encountered by trauma patients across time. Last, distinct longitudinal patient trajectories were identified that group patients according to their path through trauma transomics state maps. Unsupervised clustering of longitudinal trans-omics data identified specific clinical phenotypes while omics-based trajectories increased resolution on outcome prediction. The molecularly defined endotypes and trajectories provide an unprecedented lens to understand and potentially guide trauma patient management, opening a path towards precision medicine. This strategy presents a transformative framework that aligns with our understanding that trauma patients, despite similar clinical presentations, might harbor vastly different biological responses and outcomes.

**One-sentence summary:** Transomic analyses of longitudinal plasma samples from severely injured patients identifies endotypes and trajectories that predict clinical outcomes.

## INTRODUCTION

Trauma and hemorrhage remain a leading cause of death worldwide (*1, 2*). Underlying this mortality and the prime cause of morbidity in survivors is a poorly characterized inflammatory perturbation (*3, 4*). Care for trauma patients is complicated by multiple factors that affect clinical outcomes. First, while improvements in major hemorrhage protocols have saved lives, infusion of blood-derived products can lead to subsequent complications including coagulopathies(*5–9*). Second, tissue components released into circulation following trauma activate a non-specific inflammatory response(*10–13*). This inflammatory response is in part triggered by metabolic derangements initiated during initial tissue hypoxia and reperfusion(*14–19*). Finally, resuscitated and ‘stabilized’ patients are at risk of infection, in part due to a dysregulated inflammatory state that exhibits an attenuated response to invading pathogens(*10, 11*). Confounding this lack of knowledge of the endotypic milieu after trauma is the unknown prior health status of each individual patient and their specific responses to injury. Changes in hemostasis and inflammation, termed thromboinflammation, are central to this biology and have been characterized by our group and others(*20–25*). While significant advances over the preceding decades have saved lives, a broader understanding of how thromboinflammation drives patient-specific responses to shock and trauma is essential to provide personalized care for injured warfighters and civilians(*26, 27*).

While omics characterization has revolutionized cancer therapy and underlies personalized medicine, the care of the severely injured patient is based on overly simplified scoring systems, reductionist measures of a few mediators, or clinical “gestalt” which miss much of the existing post-injury biological dimensionality(*28–34*). Traditional clinical trials in trauma or associated sequalae have failed to provide sufficient insight and guidance to improve care(*35–40*).

Our group completed a randomized trial of plasma-based trauma resuscitation, which afforded a unique opportunity for controlled sampling, clinical care, and outcome data(*41*). We hypothesized that trans-omics profiling reveals underlying pathological differences in severely injured patients that may otherwise present with similar clinical profiles but ultimately have very different responses to treatment and clinical outcomes. Here, unsupervised clustering of molecular data obtained from severely injured patient plasma was performed to characterize metabolomics and proteomics signatures after trauma, and determine molecularly-defined clinical trajectories and endotypes.

## RESULTS

### Trauma patient characteristics and data analysis workflow

A total of 97 healthy control subjects and 118 patients with both plasma metabolomics and proteomics were enrolled in this study (**Figure 1A**). Blood was collected at specified timepoints (Field, Emergency Department arrival (ED), post-injury hours 2, 4, 6, 12, 24, 48, 72, 120, 168) until the patient’s discharge (**Figure 1B**), for a total of 856 samples (759 patient samples and 97 healthy controls). From patient plasma, 1012 proteins and 472 metabolites were quantified using untargeted DIA proteomics and metabolomics by liquid-chromatography coupled with tandem mass spectrometry (LC-MS/MS) (**Figure 1C**). Metabolomics and proteomics analyses were performed on these longitudinal samples, while clinical records were available for all patients throughout the study. Patients were grouped according to their shock severity (base excess (BE) at ED < -10 High Shock HS) and New Injury Severity Score (NISS) > 25 High Trauma HT) into High-Shock-High-Trauma (HSHT, N=35), High-Shock-Low-Trauma (HSLT, N=18), Low-Shock-High-Trauma (LSHT, N=28), and Low-Shock-Low-Trauma (LSLT, N=36) groups (**Figure 1D**). For 20 patients with missing BE at ED, omics data from an independent cohort of 333 trauma patients were employed to train and test an imputation model for value assignment (**Figure S1**). Patient characteristics have been previously detailed and are outlined here by S/T group (**Figure 1E**)(*41*). In this study, longitudinal omics trends were first analyzed in the ‘average’ trauma patient, and according to commonly used S/T grouping. Then, unsupervised hierarchical clustering of metabolomic and proteomic UMAP embeddings was performed to identify omics-based patient states and clinical patterns (**Figure 1F**). Finally, omics-based patient trajectories were identified according to their path through omics patient state space, yielding novel clinical and biological insight into trauma patient trajectories through injury and/or recovery (**Figure 1G**).

**Figure 1.**
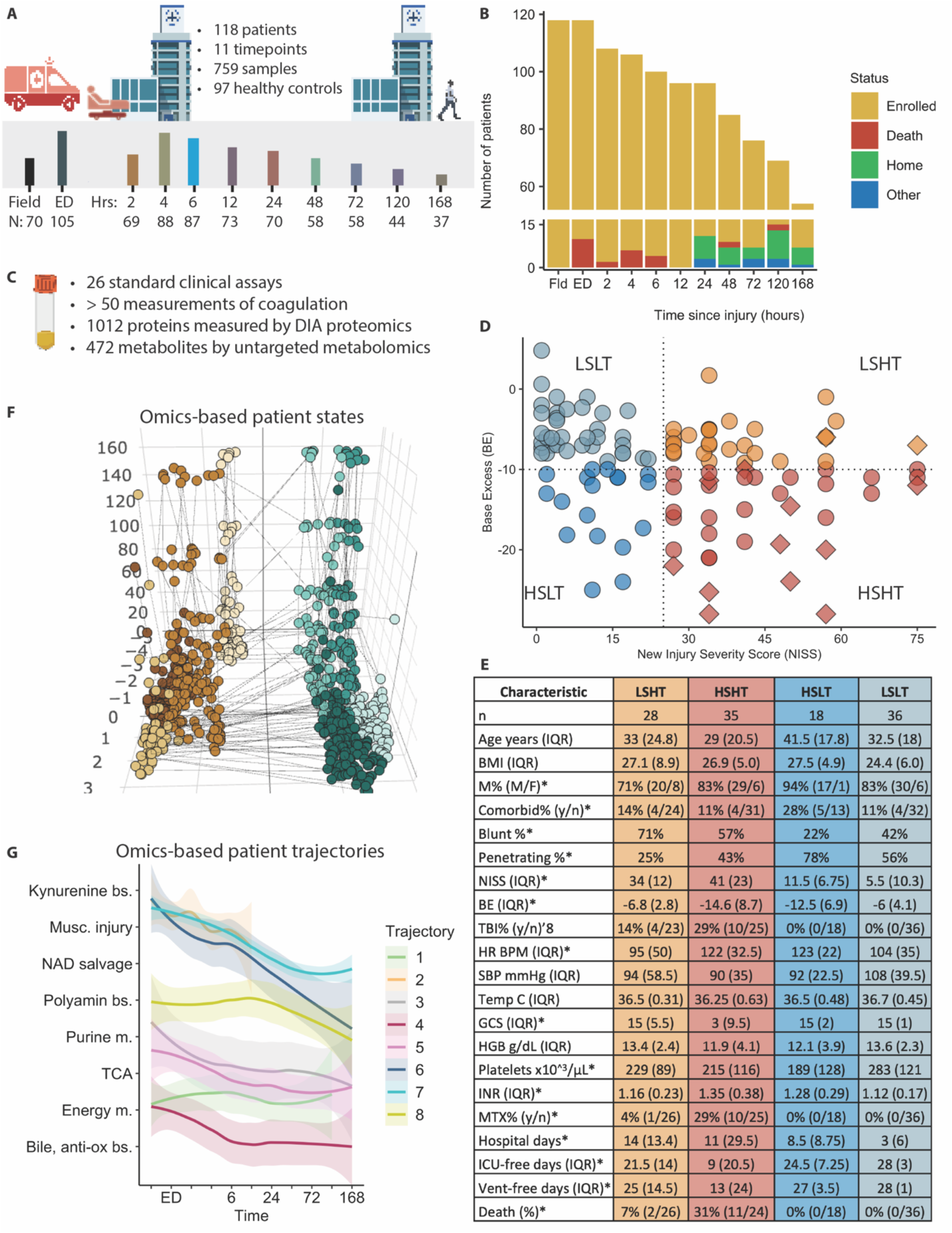
caption: Overview of data collection, study design, and patient characteristics. (**A**) Patients who met the COMBAT criteria for shock and trauma were enrolled at the Field timepoint. Blood was collected in the Field, and at 10 subsequent timepoints. (**B**) The number of patients currently enrolled, deceased, or discharged varied by timepoint. (**C**) Blood collected at each timepoint was processed for LC-MS/MS proteomics and metabolomics, and also for routine clinical blood chemistry assays. (**D**) Patients were categorized into Shock/Trauma (S/T) groups Low-Shock-Low-Trauma (LSLT); Low-Shock-High-Trauma (LSHT); High-Shock-Low-Trauma (HSLT); or High-Shock-High-Trauma (HSHT) based on NISS and BE at ED arrival (NISS >= 25 is HT; BE <= -10 is HS); diamond is deceased. (**E**) Clinical characteristics, blood chemistry measurements, injury mechanism (blunt, i.e. car crash; penetrating, i.e. gunshot), need for massive transfusion (MTX) are outlined with median values +/- IQR (*p<0.05, ANOVA, TukeyHSD post-hoc), or percent (% *p<0.05, χ^2^). (**F-G**) S/T groups were then compared to outcomes based on unsupervised clustering of omic data followed by prediction learning to identify omic-based patient trends and outcome trajectories.

### Omics patterns of the ‘average’ trauma patient

To analyze molecular patterns in the ‘average’ trauma patient, C-means clustering identified 10 unique, longitudinal kinetic patterns of omics data (**Figure S2**). For metabolomics we observed an early enrichment of the tricarboxylic acid (TCA) cycle and arginine biosynthesis, and prolonged amino acid and thiamine metabolism (**Figure 2A**, **Figure S3**). Hypotaurine and taurine metabolism peaked from 2-6 hours, and sphingolipid and histidine metabolism remained elevated until 24 hours. At 24-48 hours, we observed increases in plasma levels of nucleic acid, glutathione, and cyclic-ring amino acid metabolism; as well as hormone and fatty acid biosynthesis. Finally, 5-7 days post-injury had high levels of metabolites involved in biosynthesis of bile acids, aminoacyl-tRNA, proteolysis and amino acid metabolism.

**Figure 2.**
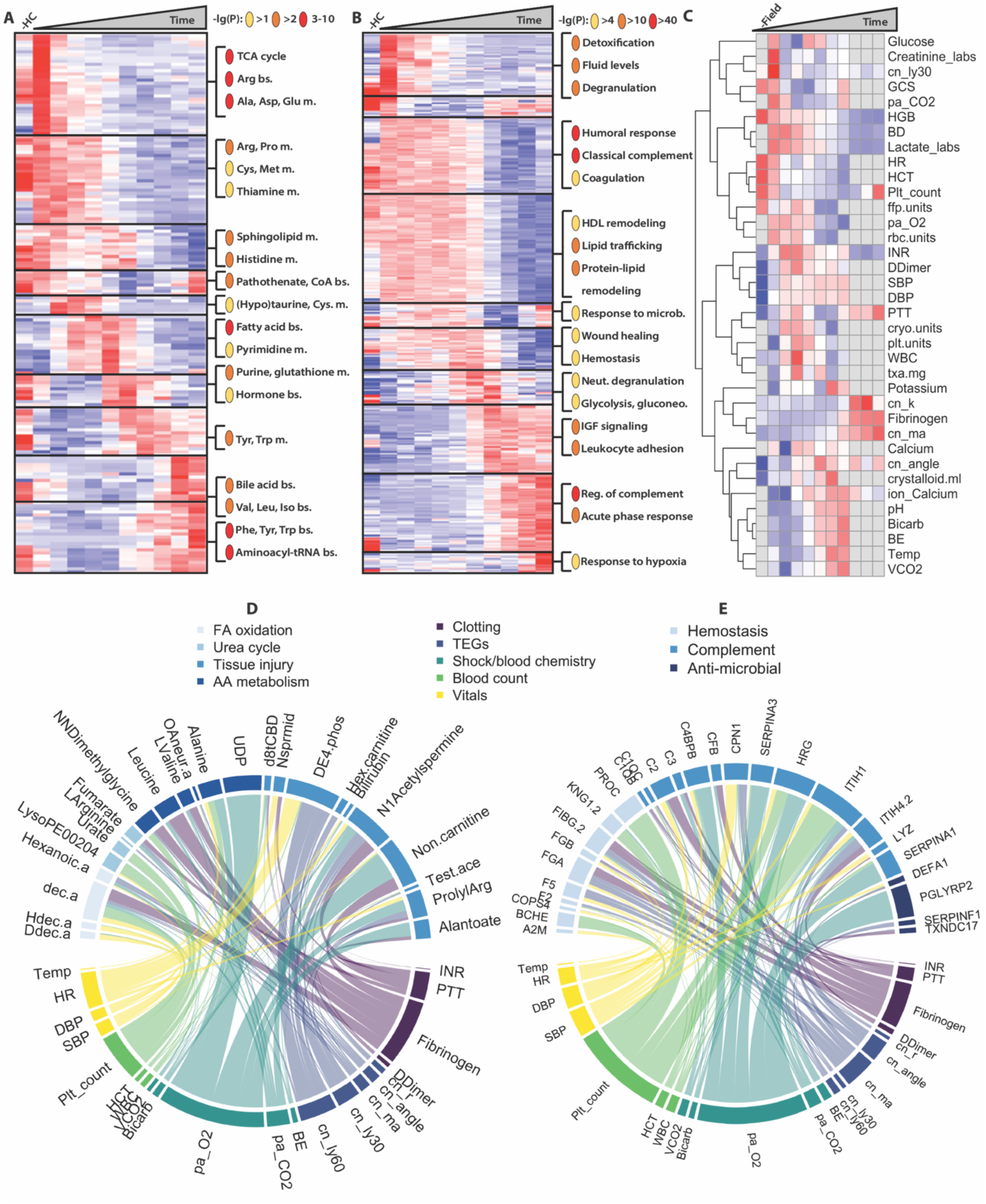
caption: Metabolomic, proteomic, and clinical trends of the “average” trauma patient. (**A**) Heatmap of median values of metabolites sharing similar temporal, kinetic patterns from C-means clustering (left side); and enrichment terms of each cluster (right side, enrichment scores from MetaboAnalyst). (**B**) Heatmap of median values of proteins sharing similar temporal, kinetic patterns from C-means clustering (left side); and GO enrichment terms of each cluster (right side, enrichment scores from Metascape). (**C**) Clinical chemistry labs showing the relative, median value at each timepoint. (**D-E**) Strength of association (partial slope from mixed modeling) between clinical measurements and associated metabolites D) or proteins E), with line thickness showing relative partial slope. dec.a = Decanoic acid. Ddec.a = Dodecanoic acid. Hdec.a = Hexadecenoic acid. Hexanoic.a = Hexanoic acid. OAneur.a = OAcetylneuraminic.acid. d8tCBD = delta-8-tetrahydrocannabinol. Nsprmid = N1-Acetylspermidine. Hex.carnitine = Hexenoylcarnitine. Non.carnitine = Nonanoylcarnitine. DE4.phos = DErythrose.4phosphate. Test.ace = Testosterone.acetate. ProlylArg = Prolyl-Arginine. TEG = Thromboelastography.

For proteomics enrichment, there were initially high levels of proteins involved in detoxification, management of body fluid levels, and neutrophil and platelet degranulation (**Figure 2B, Figure S4**). This detoxification phase was followed by the initiation of a broad humoral immune response, complement activation, and (regulation of) coagulation including clot formation and fibrinolysis. These proteins remained elevated out to ∼24hrs, and were accompanied by a concomitant rise in protein-lipid (i.e. HDL) remodeling and lipid trafficking; an anti-microbial response, and a general response to wounding and hemostasis. From 24-48hrs there was a second spike in neutrophil degranulation proteins and an elevation in proteins involved in glycolysis and gluconeogenesis. Following this second spike was an acute phase response (APR), leukocyte adhesion and activity, negative regulation of complement, and a response to hypoxia.

For clinical measurements, consistent with the literature, we observed expected trends in clinical markers of shock, including early elevations in heart rate, lactate, and BE alongside reduced blood pressure that normalized quickly after resuscitation and hemorrhage control (**Figure 2C**). Additional measures of metabolic acidosis (pH, bicarbonate) were consistent with these trends as well. The average patient arrived coagulopathic with reduced MA and fibrinogen alongside elevated LY30, which were also corrected over time, though not as abruptly as normalization in physiologic parameters of hemorrhagic shock. Patients arrived, on average, hypocalcemic with exacerbation of this hypocalcemia within the first 2 hours, secondary to the reported influx of calcium into the endothelium. Subsequent administration was likely responsible for the correction in ionized calcium.

Next, linear mixed modeling (adjusted for age, sex, shock/BE, injury/NISS, and time) was performed to identify proteins and metabolites highly associated with common clinical measurements (**Figure S5**). For metabolomics, strongest relationships were between carnitines and phosphates with platelet count, P_a_O_2_, and fibrinogen (**Figure 2D**); measures of hypoxia and fatty acid oxidation and vital signs; and between tissue injury metabolites (spermine, spermidine, bilirubin, carnitines) and clotting. For proteomics, the strongest relationships were between clotting assays and coagulation and complement proteins; blood gas and APR proteins; and platelet count with complement and coagulation proteins (**Figure 2E**).

### Omics differences by shock severity and tissue injury

Outcome for trauma patients is thought to be governed by a ‘golden hour’ where interventions within the first 60 minutes are deemed to be crucial to establishing a trajectory towards survival and good outcome. To identify omics expression patterns immediately following injury within this ‘golden hour’, patients were categorized by S/T scores (Figure 1D), and analyte levels were compared among these groups. A proteomic signature of tissue injury from significantly higher levels of 186 proteins was identified in the HT groups at ED (**Figure 3A**, **Figure S6**). Many of these were histones, extracellular (COL18A1, PRG4) or cytosolic (ACTAs, TUBBs, RHOA) proteins and are involved in cell metabolic processes (PSAT1, ARG1, ACAT2) or detoxification (ENO1, ADH, ALDH, SOD1) (**Figure 3B**). Significantly elevated proteins within the HS groups were primarily involved in antioxidant activity (FABP1, TXN, GSTA1) or glycolysis and gluconeogenesis (ALDOB, TPI1, LDHA) (**Figure 3C**). Beyond ED, HT patients had significantly elevated levels of glycolytic (ADH1B, GOT1, PGAM1), detoxification (FABP1, GSTA1, PRDX6, S100A9, CAT, CPN1), and tissue injury and hemolysis proteins (MB, HBA, HBB) as well as decreased gelsolin (GSN) to bind circulating actin (**Figure 3D-G**). From 12-24 hours, within the HT groups were elevated levels of inflammatory proteins (SAA1/2, CRP, IL1RL1), peptidases and anti-microbials (CTSB, LBP, CHI3L1), as well as protease inhibitors (SERPINA1, TIMP1) and the tPA/fibrinolysis inhibitor SERPINE1. From 24-48 hours, HT groups also showed significantly elevated fibrinogen chains (FGA, FGB, FGG), and decreased complement subunits (C1QB, C1QC).

**Figure 3:**
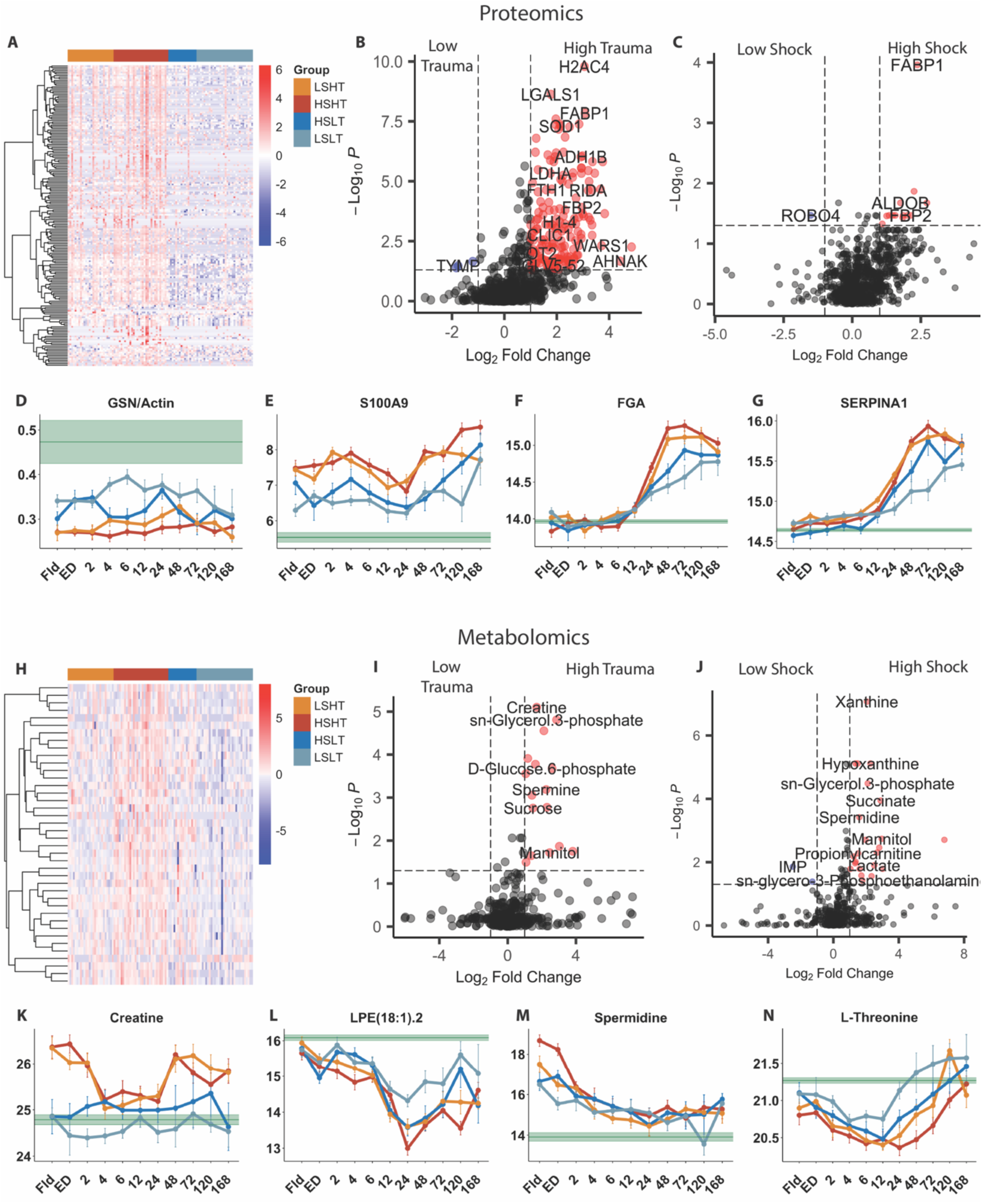
Golden hour trauma patient plasma omics. (**A**) Proteomic heatmap of significant analytes at ED by S/T group (FDR-corrected p<0.05 following ANOVA). (**B**) Proteomic volcano plot between HT (right. N=63) vs. LT (left N = 54) colored by significance (FDR-corrected p<0.05 following t-test) and relative fold change (red for HT/LT > 2x; blue for LT/HT > 2x). (**C**) Proteomic volcano plot between HS (right. N=53) vs. LS (left. N=64) colored by significance (FDR-corrected p<0.05 following t-test) and relative fold change (red for HS/LS > 2x; blue for LS/HS > 2x). (**D-G**) Line plots of median +/-SEM of tissue injury (GSN/Actin), detox (S100A9), clotting (FGA), and protease inhibitor (SERPINA1) proteins over time (*p<0.05 Sidak correction on estimated marginal means of mixed model). Green line = HC values. (**H-J**) Similar analysis for metabolomics showing S/T heatmap (H), and HT/LT (I) and HS/LS (J) volcano plots at ED. (**K-N**) Line plots of metabolites involved in tissue injury (creatine), lipid synthesis (LPE(18:1)), hypoxia (spermidine), protein synthesis (L-Threonine).

For metabolomics, 37 metabolites were significantly elevated in the HSHT group (**Figure 3H**, **Figure S7**). Within HT, a wide range of metabolites including amino acids, nucleotides, galactose, and propanoate, as well as elevated levels of hypoxia markers succinate and creatine (**Figure 3I**). In HS patients, there were significantly elevated levels of polyamines, TCA cycle intermediates (succinate, fumarate, malate), lactate, beta-alanine and markers of glycero(phospho)lipid metabolism (**Figure 3J**). Beyond ED, high levels of these metabolites persisted in the HSHT group until 6 hours, at which time the LSLT group showed elevated metabolites involved in biosynthesis of aminoacyl-tRNA, amino acids, and fatty acids which indicated metabolic recovery (**Figure 3 K-N**).

HT patients frequently sustain traumatic brain injury (TBI) which complicates treatment and outcomes. For patients with TBI, proteomic differences in HSHT (elevated EXT2), LSHT (elevated SIGLEC14, PAICS, HPRT1, COMT; decreased ASGR2), or HT combined (CCL18 lymphocyte attraction) indicated differences in serum glycoprotein homeostasis, purine synthesis and metabolism, cell adhesion, and catecholamine degradation (**Figure S8**). Further, TBI patients within HT groups had consistently elevated levels of dopamine, and metabolites involved in (hypo)taurine metabolism and catecholamine biosynthesis.

### Classification prediction and VIP analysis

Predicting the need for massive transfusion (MTX), and adverse outcomes including coagulopathy (INR>1.4), acute lung injury (ALI), extended ICU- (ICU-free days < median) and ventilator time (Vent-free days < median), and death can be difficult. Ensemble machine learning which creates an optimized, complied model was employed to predict outcomes and the need for MTX following trauma (**Figure S9**)(*42*). ED trans-omic data were split into 75%-25% training and test sets, respectively. With AUC ∼90% for death, coagulopathy, MTX, and ventilator time, and ∼80% for ALI and ICU time, each outcome was associated with a unique panel of analytes contributing to Variable Importance in Projection (VIP) scores (**Figure S10**). Top VIP features were detected at significantly higher levels in patients who experienced worse outcomes. Of note, both death and MTX VIPs indicated hypoxia, short-chain fatty acid metabolism, and elevated polyamines; INR VIPs indicated tissue and RBC lysis, and were involved in oxidative response; extended ICU stay, and extended ventilator VIPs enriched for glycolysis and high tissue oxidation, while ALI VIPs enriched for catabolic processes, systemic inflammation, and RBC lysis. Further, models trained on an independent omics dataset of 333 trauma patients from a separate cohort (TACTIC study) predicted death (AUC 91%), ICU-free days (84%), and ventilator-free days (87%) in the current dataset (**Figure S11**).

### Mapping metabolic trauma patient states

Next, a map of omics-based trauma patient states was created. Hierarchical clustering on 2D UMAP embeddings of metabolomics data yielded 8 distinct metabolic patient states (**Figure 4A**, **Figure S12**). These trauma patient states that differentiated datapoints according to underlying metabolic differences also showed clinical differences, and served as the basis for the omics-based patient trajectories presented next. Datapoints associated with deaths, low ICU-free days, and severe injury (HSHT) were scattered across several patient states (PS 3, 5, 8. **Figure S12**). Two main metabolic meta states were observed and generally characterized by catabolism (Right), and energy production and biosynthesis (Left. **Figure 4A**. **Figure S13**). Metabolic signatures (median expression within the upper quartile) revealed unique sets of metabolic pathways enriched in each patient state (**Figure 4B**, **Figure S14**). The density of datapoints per timepoint indicated the temporal nature of each patient state, with patient states classified as early injury (1, 5, 8), mid- injury (2, 3, 4), and late injury (6, 7) (**Figure 4C**). States 1 (lysophospholipids), 3 (pentose phosphate pathway), 5 (aromatic amino acids tryptophan and its indole derivative; tyrosine; branched chain and other amino acids – valine, citrulline; medium to long-chain acyl carnitines – AcCa 6:1, 18:0, 18:1; purine metabolites: hypoxanthine, urate, caffeine, theophylline; lysophosphatidyl-ethanolamines, -cholines and -serines of the 16:0 and 18:1; 20:4 C series) and 8 (markers of blood transfusion – mannitol and S-adenosylhomocysteine; antibiotics – amoxicilline) showed an enrichment at earlier time points, followed by progressive decline. Opposite trends were observed for states 2 (exogenous metabolites from iatrogenic intervention), 6 (carnitine metabolism and hemolysis markers) and 7 (pharmacological intervention with acetaminophen and pain management with opioids). Amino acid metabolism, especially sulfur amino acids, were over- represented in state 4 (cysteine, homocysteine, kynurenine, leucine, methionine sulfoxide, phenylalanine, taurine), which showed no major temporal trends.

**Figure 4:**
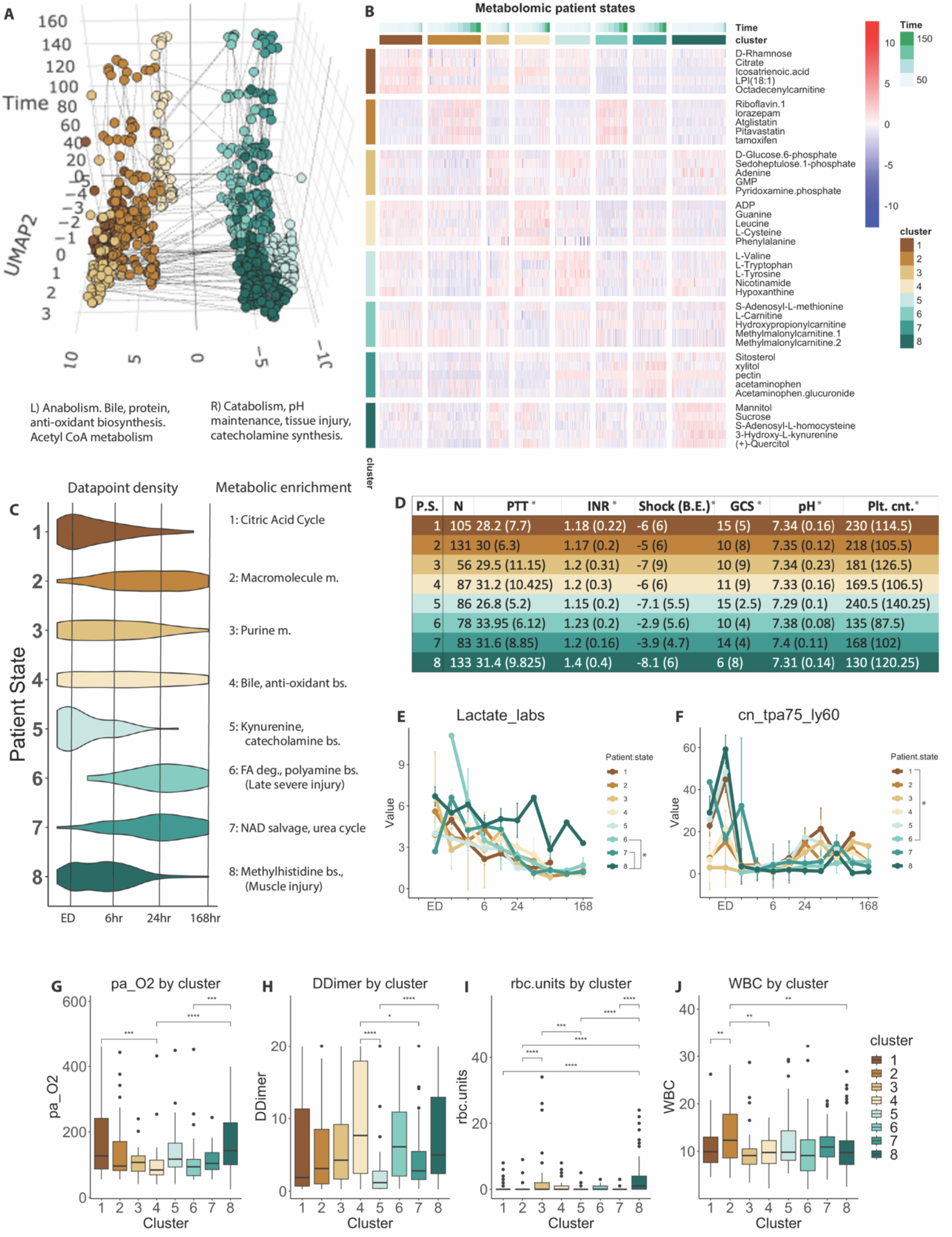
Metabolomic patient states. (**A**) 3D UMAP of clustered metabolomic data stratified by timepoint; two meta groups generally defined by biosynthesis of macromolecules (L) versus elevated catabolism (R). Each colored circle is a patient-timepoint, while each line connects a patient’s datapoints across time. (**B**) Each state had a significantly elevated metabolic signature that enriched for different cellular metabolic processes. Top 5 shown, full list in Figure S14. (**C**) The density of datapoints within patient states varied temporally, with early injury (states 1, 5, 8), mid-injury (2, 3, 4), and late-injury (6, 7) enriched patient states. (**D**) Clinical measurements for each metabolic state is outlined as median +/-IQR (*p<0.05 via ANOVA, TukeyHSD post-hoc). (**E-J**) Line graphs (*p<0.05 Sidak correction on estimated marginal means of mixed models) showing temporal trends and (**G-J**) box plots showing static trends (*p<0.05 by ANOVA, TukeyHSD post-hoc) of clinically relevant measurements. Full list of clinical results in SF15.

Clinically, each state expressed unique static and dynamic clinical measurements (**Figure 4D-J**, **Figure S15**). State 8 exhibited the most perturbed vital signs (lowest body temperature and blood pressure), reflected the most shock (lowest BE and bicarbonate, highest lactate), and was the most coagulopathic (highest INR, lowest clotting factors). This state was further characterized by persistent brain injury (low GCS) and a longitudinal fibrinogenemic and thrombocytopenic coagulopathy. Other notable differences included those in blood chemistry (low P_a_O_2_ in state 4; low pH states 3 & 8), blood count (high WBCs, platelets, and hemoglobin and hematocrit in states 1, 2, 5), clotting (lowest PTT, INR, D-dimer in state 5; lowest Von Klauss fibrinogen in state 5), and transfusion of blood products (high tranexamic acid (TXA) in state 4; high RBCs, FFP, and crystalloids in states 4 & 8) (**Figure 4G-J**). Several states were clinically similar reflecting our hypothesis that metabolomic differences would distinguish otherwise clinically indistinguishable states.

### Mapping proteomic trauma patient states

Hierarchical clustering of trauma patient proteomics UMAP embedding also mapped 8 distinct patient states, demarcated by proteomics differences across datapoints (**Figure 5A**. **Figure S16**). Datapoints associated with death, low ICU-free days, and severe injury (HSHT) were more localized, demarcating more severely (Right) and less severely-injured (Left) patient states (**Figure S16**). As with metabolomics, the density of datapoints within each patient state identified early- (states 1, 2, 7), mid- (3, 8), or late-injury (4, 5, 6) patient states (**Figure 5B**). Further, the differences in proteomics signatures identified unique biological processed enriched within each patient state (**Figure 5B-C**). States 1 (early severe injury) and 3 (mid- severe injury) showed high levels of tissue injury proteins that enriched for catabolism, response to ROS, and neutrophil and platelet degranulation (state 1), and proteasome expression, FA degradation, and response to stimuli in state 3 (**Figure 5B**, **C**. **Figure S17**). For later-injury states, state 4 had high levels of acute phase response (APR) proteins and early complement activation; while state 5 enriched for complement, cell adhesion, and acute inflammation. Other patient states contained less injured patient-timepoints and had higher levels of coagulation proteins such as F2 and F13B (states 2, 6, 8). State 7, an early injury cluster in between high injury state 1 and low injury state 2 had high levels of proteins involved in hemolysis and oxygen transport.

**Figure 5:**
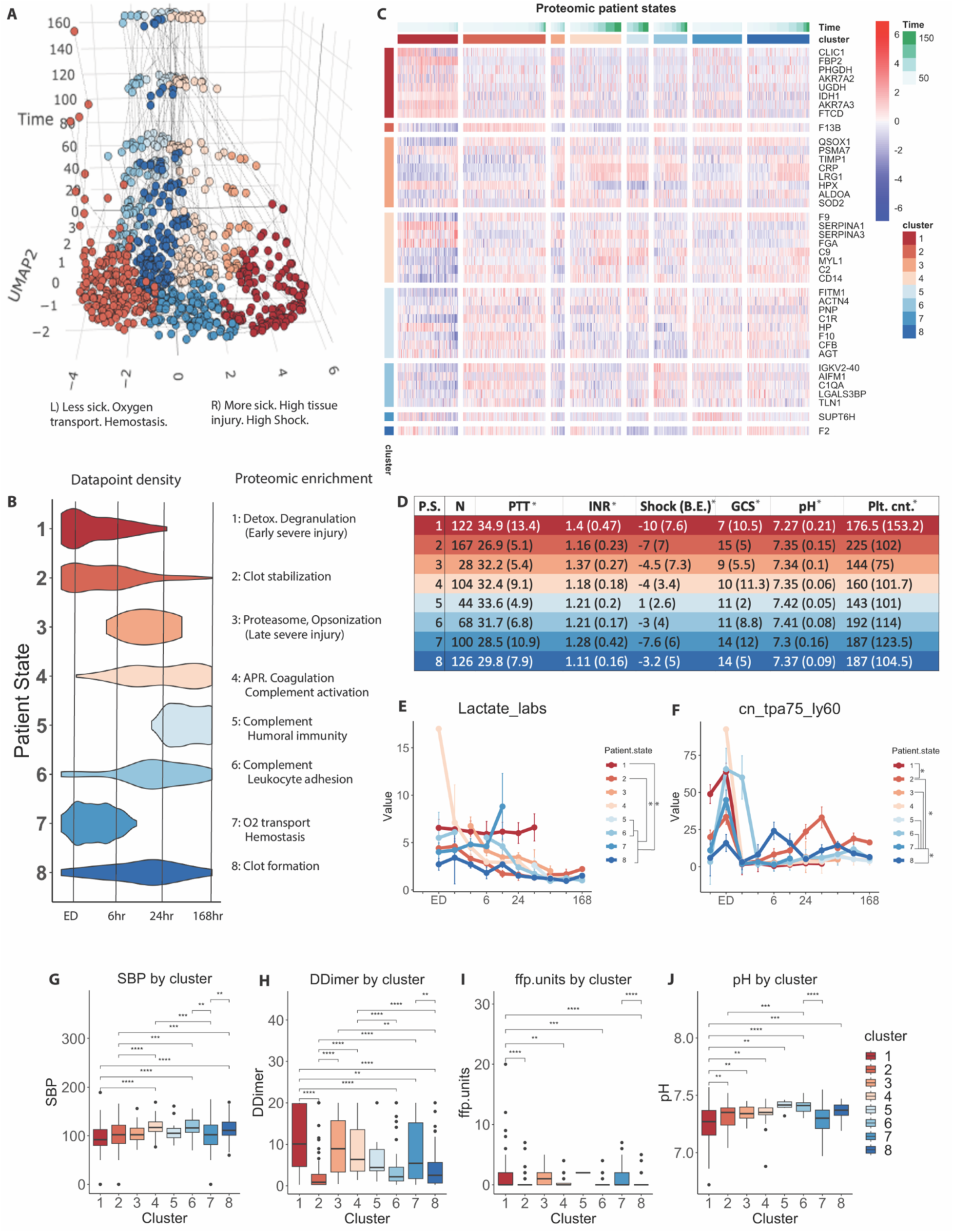
Proteomic patient states. (**A**) 3D UMAP of clustered proteomic data stratified by timepoint with sicker (R), and less sick (L) sides. (**B**) Patient states varied temporally with early injury (states 1, 2, 7), mid-injury (3, 8), and late-injury (4, 5, 6) enriched clusters. (**C**) Each state had a significantly elevated proteomic signature that enriched for different biological processes. (**D**) Clinical measurements for each proteomic state is outlined as median +/-IQR (*p<0.05 via ANOVA, TukeyHSD post-hoc). (**E-J**) Line graphs (*p<0.05 Sidak correction on estimated marginal means of mixed models) showing temporal trends and (**G-J**) box plots showing static trends (*p<0.05 by ANOVA, TukeyHSD post-hoc) of clinically relevant measurements. Full list in SF17.

Clinically, states 1 and 3 had most perturbed vitals (lowest temp, blood pressure, GCS, and highest heart rate) and blood chemistry (lowest VCO_2_, BE, bicarbonate; high lactate) (**Figure 5D-J**, **Figure S18**). These states also indicated coagulopathy and evidence of fibrinolysis (high PTT, INR, D-Dimer, TEGs; low clotting factors 7 and 9), and received high amounts of transfused blood products (RBCs, platelets, cryoprecipitate). States 2 (high P_a_O_2_, platelets, HGB, HCT, clotting factors; low body temp, D-Dimer), 7 (high P_a_O_2_, lactate; low body temp, clotting factors), and 8 (low heart rate, D-Dimer; high VCO_2_) were mixed. The remaining states 4, 5, and 6 were clinically similar but had intermediate levels of clinical measurements operating at later timescales. Notable differences include SBP (high in states 4, 8; low in states 1, 5), D-dimer levels (low in states 2, 6, 8; high in states 1, 3, 4, 7), transfusion of FFP (high in states 1, 7), and pH (low in state 1, high in states 5, 6).

The combined metabolomics and proteomics model contained aspects of both models (**Figure S19**). Early severe injury state 4 showed high levels of proteins related to tissue injury, metabolism and biosynthesis of organic molecules, response to stress, and neutrophil degranulation. Further, high levels of succinate, lactate (hypoxia), creatine, and carnitines were detected. Late severe injury state 8 showed high levels of analytes involved in glycolysis, gluconeogenesis, as well as methyl-histidine and pyruvate metabolism. Adjacent states 6 and 7 enriched for proteins involved in the complement and coagulation cascades, acute phase response, bile acid synthesis, and fatty acid metabolism. Less sick patient states (1, 2, 3, 5) showed high levels of taurine metabolism, complement proteins, glycolysis, fatty acid synthesis, and immunoglobulins.

### Identifying endotypes through metabolic-based trauma patient trajectories

Previously, disease endotyping with molecular data has identified unique sub-phenotypes associated with differing outcomes and treatments within a broader patient population(*43–47*). Utilizing trauma patient state omics maps created above to identify trauma endotypes, distinct omics-based trauma patient trajectories were created by grouping patients that shared the same initial and 12/24 hour metabolic states (**Figure 6A**). Thus patients in each trajectory (T) had a unique metabolic pathway from initial injury through discharge; each T was defined strictly by changes in metabolism across time (**Figure 6B, Figure S20**). Clinical sub-phenotypes were identified which had different outcomes that coincided with metabolic trajectories despite having similar presenting clinical characteristics. For major clinical measurements upon arrival, there were no significant differences in injury severity among trajectories; slight elevations in BD and lactate in T1 and T8; elevations in INR in T6 and T7; and lower GCS in T7 (all n.s. **Figure 6D-H**). T1 and T6 which began in C.A.C enrichment and methylhistidine metabolism (muscle injury), respectively received the most massive transfusions (MTX) and had the highest mortality rates (**Figure 6I,K**). T3 and T8 both had high incidences of TBI and blunt injury mechanism and developed liver organ failure (OF), while patients in T3 developed more lung OF (56% vs. 18%) and patients in T8 had more comorbidities (14% vs. 0%) (**Figure 6L-P, Figure S21**). Metabolically, T3 remained in a state of bile/cholesterol and anti-oxidant synthesis while T8 exhibited high rates of transition between polyamine synthesis and purine metabolism. T7 had the highest rate of penetrating injuries, developed liver and heart OF at the highest rates, and also lung OF at a high rate (50%), but 0% mortality and fewer ICU-free days on average. Metabolically, this trajectory began in high methylhistidine metabolism (tissue injury) and ended in NAD salvage.

**Figure 6:**
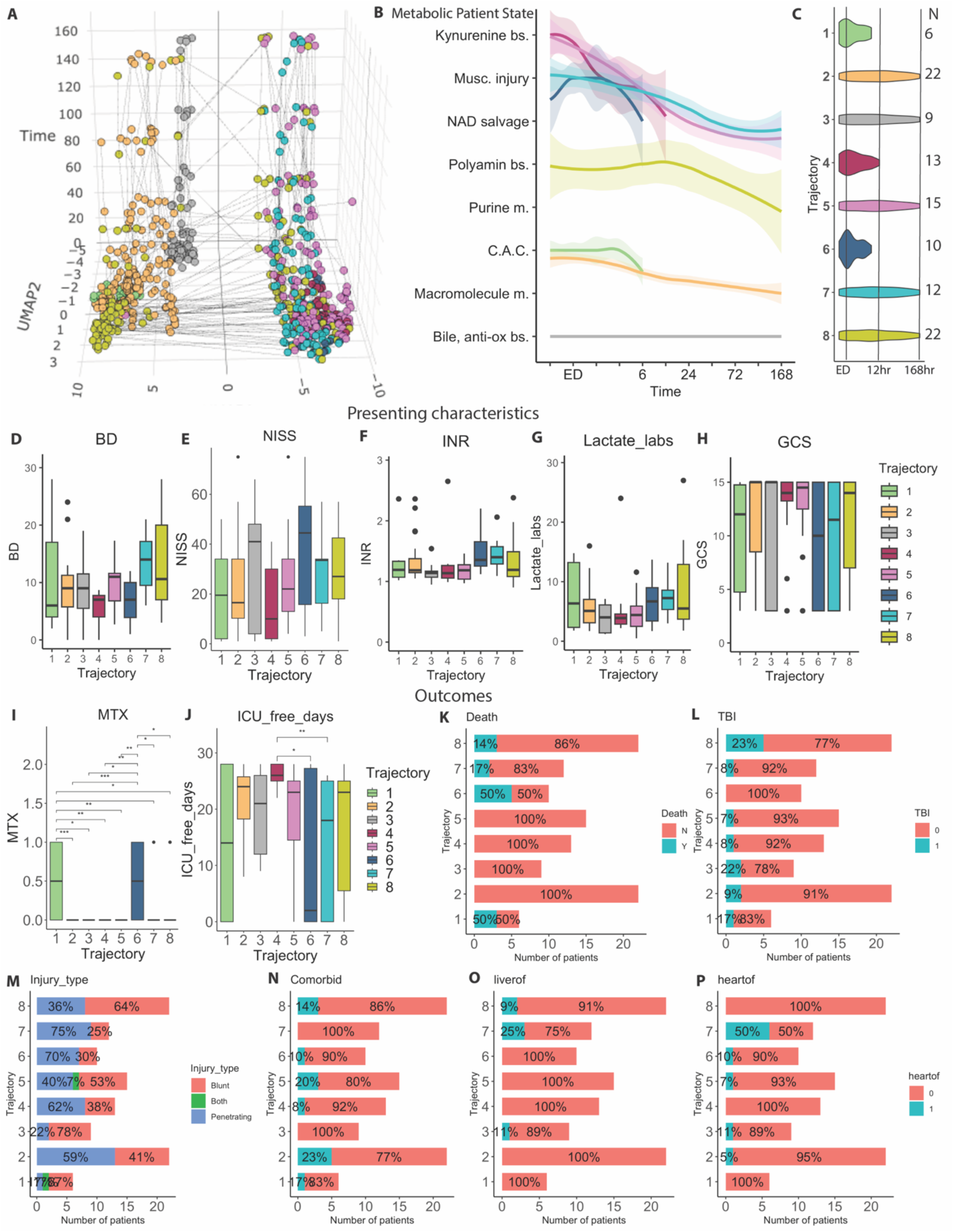
Metabolomic patient trajectories. (**A**) Trauma patient metabolomic UMAP space colored by metabolomic patient trajectories. (**B**) Line plot of each patient trajectory through metabolic patient state space +/- 95% CI; x-axis is time, y-axis is patient state. (**C**) Temporal nature of trajectories, with most patients in trajectories 2, 4, and 6 leaving relatively early. (**D-P**) Box plots (*p<0.05 ANOVA, TukeyHSD post-hoc) and proportions (**K-P**) of clinical measurements for each trajectory.

Comparing the temporal nature of trajectories, both T1 and T2 began in C.A.C. enrichment. T1 was short-lived with high mortality (50%), while patients in T2 switched to metabolism of most biomolecules (amino acids, carbohydrates, fatty acids) and had 0% mortality. While there were no clinical differences between these trajectories, T1 presented with continuously elevated hypoxia-related metabolites and carnitines and T2 had low levels of these metabolites and elevated icosatrienoic acid. Similarly, T4 and T5 both began in the same state of high kynurenine biosynthesis. Patients in T4 had the highest ICU free days were and discharged to home, rehabilitation, or another hospital; while patients in T5 switched to an NAD salvage metabolic state and were sicker with fewer ICU and ventilator-free days, longer hospital stays, and developed more lung OF (40% vs. 8%).

### Identifying endotypes through proteomic-based trauma patient trajectories

For proteomics, 12 trajectories were identified, each with a unique path through the proteomic trauma patient state map and that represented more severely- vs. less severely-injured, short- vs. long-lived, and transitional trajectories (**Figure 7A-C**, **Figure S22**). As with metabolic trajectories each proteomic trajectory was defined by its proteomic signature, and was associated with a clinical sub-phenotype despite similar presenting clinical characteristics for some trajectories (**Figure S23**). T1-3 had the highest tissue injury, while T1 was the most severely injured with and highest INR and BD (severe shock) and lowest GCS (**Figure 7D-H**). These trajectories all begin in the same proteomic state of detoxification and degranulation. T1 then oscillates across opsonization and complement activation before ending around 12 hours with an 82% mortality rate (**Figure 7I**). T2 remains in detoxification and travels through opsonization, complement activation, and APR before landing in hemostasis and clot stabilization with a lower (8%) mortality rate; while T3 initially transverses these states before ending in the same state of clot formation and stabilization also with an 8% mortality rate. Survivors in T2 vs. T3 are among the sickest with fewer ICU-free days and longer hospital stay, yet develop differences in other outcomes including lung OF (92% vs. 62%), heart OF (50% vs. 23%), and TBI (17% vs. 54%) reflective of blunt injury type (67% vs. 92%) (**Figure 7J-L, Figure S23)**. Starting in clot stabilization, T4-7 followed different proteomic trajectories across patient states with main differences in toxicology screens (T4 highest cocaine, amphetamine, and opioids) and lung OF (**Figure 7O, Figure S23)**. Short-lived T9 had the second highest mortality rate (29%). Its counter-trajectory T10 began in the same proteomic state of hemostasis, yet had 0% mortality along with a higher incidence of pre-existing obesity (47% vs. 0%) and BMI (27.1 vs. 23.4 kg/m^2^ median) (**Figure 7M)**. T8 which began and remained in complement activation and clot formation was unique in having significantly higher age than other trajectories (**Figure 7P)**.

**Figure 7:**
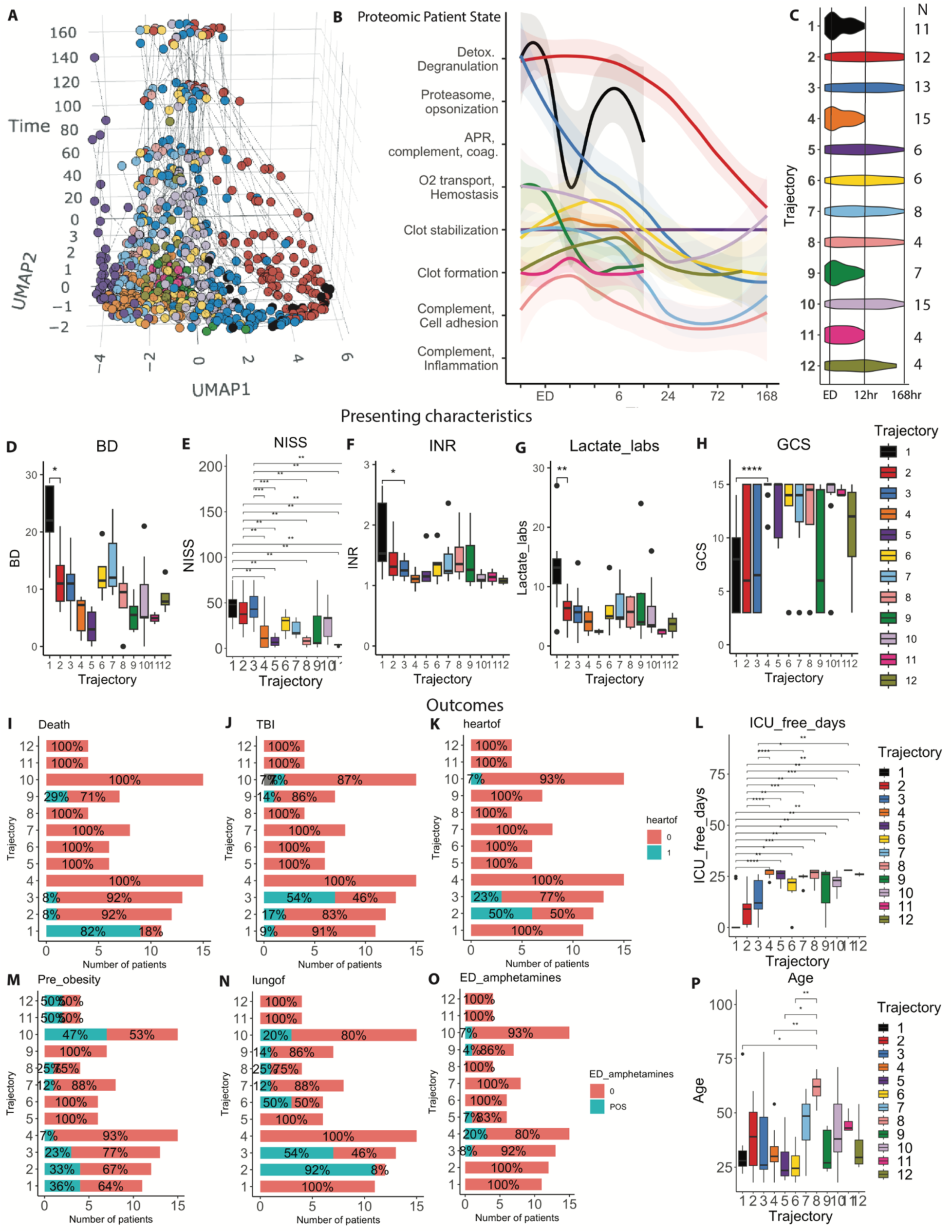
Proteomic patient trajectories. (**A**) Trauma patient proteomic UMAP space colored by proteomic patient trajectories. (**B**) Line plot of each trajectory through proteomic patient state space +/- 95% CI; x-axis is time, y-axis is patient state. (**C**) Temporal nature of trajectories. (**D-P**) Box plots (*p<0.05 ANOVA, TukeyHSD post-hoc) and proportions of clinical measurements for each trajectory.

### Integrating omics patient states and trajectories for outcome prediction

Last, the clinical utility of integrating combined proteomics with metabolomics-based patient states and trajectory maps was determined as personalized omics data could improve outcome prediction for some patients. Generally speaking, patients with HS and/or HT were scattered across all combined omics start states and trajectories (**Figure 8A**). When categorized by combined omics patient states, patients were grouped into distinct trajectories defined by longitudinal changes in their metabolism and proteome (**Figure 8B,C. Figure S19**). Integrating these data, patients with similar start states were further categorized into divergent outcomes and phenotypes (**Figure S24**). For example considering death, patients with high trauma, high shock, and who are coagulopathic (INR>1.4) are considered critically injured with poor outcomes. In our dataset with 13 deaths, 55% (6/11) HSHT coagulopathic (INR>=1.4) patients died, with the remaining 5 deaths lacking coagulopathy or shock. When considering the omics start states, 92% (12/13) deaths occurred in patients who had high tissue injury and also began in omics states 4 and 8 (detoxification, high polyamines, methylhistidine and pyruvate metabolism). Simply starting in these states gave patients a 41% (12/29) chance of mortality regardless of presenting clinical characteristics. Integrating trajectory information, patients who remained in omics states 4 and 8 beyond ED had a 100% (12/12) chance of death (T5; 2 patients in T6), while patients beginning in the same states whose omics transitioned to either taurine metabolism, APR, and glycolysis (T7); or bile synthesis, and fatty acid and riboflavin metabolism (T6) had 0% mortality (**Figure 8D-G**). This is despite no significant differences in NISS and shock, although non-significant differences in INR among T5-7. Considering other outcomes, all patients in T1-3 began in omics states 1 (taurine metabolism) or 2 (glycolysis and complement activation), with no significant differences in NISS, BD, or INR (**Figure 8E-G**). Patients that remained in state 1 (T1) had, on average fewer ventilator-free and ICU-free days although not significant (p<0.06 for all) (**Figure 8H,I**).

**Figure 8:**
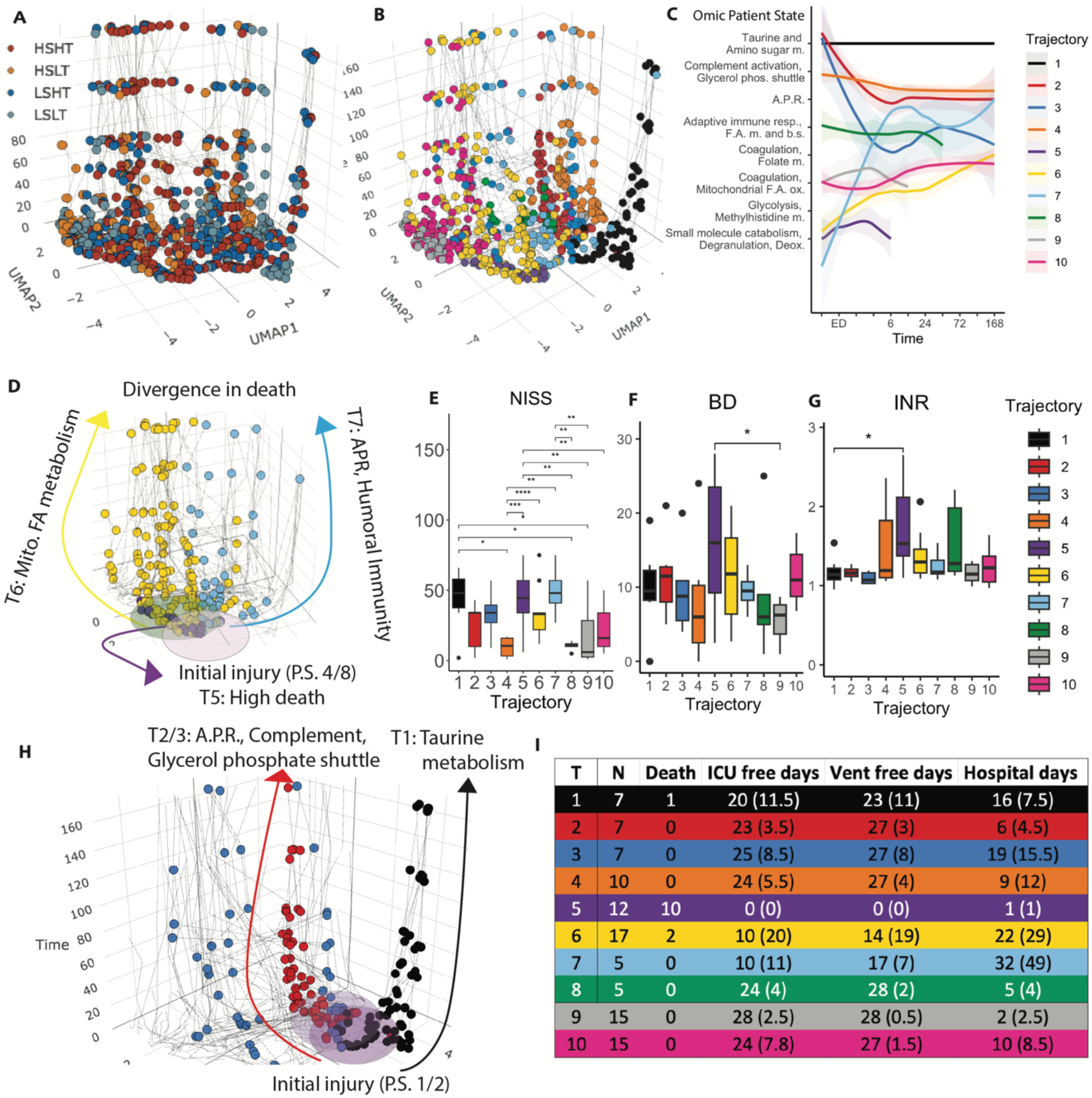
Trans-omics patient trajectories and outcomes. (**A**) Trauma patient combined omics UMAP space colored by tissue injury (NISS) and shock (BD). (**B**) UMAP colored by combined omics patient trajectories. (**C**) Line graph of combine omics patient trajectories through omics patient states. (**D**) Trajectories of all patients which began in the same omics states 4/8; T5 (purple) has high mortality, T6 then exits into mitochondrial fatty acid metabolism (yellow), T7 then exits into Acute Phase Response (light blue). (**E-G**) Comparison of clinical measurements across trajectories. (**H**) Trajectories of all patients who began in the same omics states 1/2; T1 remains in taurine metabolism (black), T2/3 exit into A.P.R. and complement activation (red, blue). (**I**) Table of clinical outcomes across trajectories.

## DISCUSSION

Clinicians rely on years of experience to gain the clinical acumen necessary to characterize the physiologic state of a patient and predict the patients’ likely clinical transition and outcome. As more data becomes available, better clinical measurements and perturbations continue to narrow the confidence interval of a patient’s given physiologic state. Unfortunately, the trauma literature is filled with single and multivariate regression models and scoring systems, an approach that leads to overfit and mis-specified models for outcome prediction. Most importantly clinicians often lack an ability to understand why two seemingly ‘identical’ patients have divergent outcomes. To remedy this gap, here we utilized longitudinal trans-omics data to provide a more comprehensive characterization of the post-trauma response to injury and subsequent treatment. These findings provide a more holistic view of the post-trauma milieu, and most importantly, reveal that patients separate into biologic phenotypes which cannot be discerned by demographic, injury, or clinical data. Recently, several groups have characterized gene expression, and trans-omic changes after trauma. These analyses include test and training set clustered prediction modeling and evaluation of endotypes related to beneficial outcome from the PAMPer trial(*47*). These advances in trans-omics studies have allowed for readouts closer to patient phenotype. However, to date, they have not been used to identify unsupervised molecular patient states or trajectories.

Our group and others have utilized longitudinal multivariate biomarker data to characterize illness and injury, have shown effective prediction of outcomes, and garnered biologic pathway insight(*42–59*). Inspired by this work, here we report trans-omics informed states that map trauma patient biology across time, and longitudinal patient trajectories that distinguish clinical phenotypes. Finally, comparing patient trajectories which begin in similar states offers a trans-omics explanation for divergence in patient outcomes. Injured patients were grouped via their underlying molecular signatures, rather than demographics or physiologic readouts. This allowed for an agnostic characterization in line with our clinical conjecture that patients often have similar demographic, injury or physiologic traits yet seemingly different underlying biology, responses to injury and treatment, and resultant outcomes.

Traditional statistical approaches do indeed often work for prediction after trauma. Unfortunately these predictors (i.e. NISS, BD, lactate, INR, etc.) while statistically valid miss multivariate patterns and do not work for those that are not fit perfectly to the prediction models. This is especially true in our data. For example Figure 8 does show higher NISS and lower BD in those who do poorly but this prediction did not work for 40+% of patients. For these types of patients, traditional models would be overfit or mis-specified and would not be accurate predictors. Our findings here combat this issue. Indeed we believe this to be the most unique aspect of this work, namely the ability to understand what drives divergent clinical trajectories and outcomes. We expect molecular-based trajectory data like these –when optimized, to augment our current patient classification, and outcome prediction. For example, from metabolomics there was high mortality in T1 compared to T2 which began in the same state but these survivors switched to a new metabolic state. In proteomics, T10 had no mortality compared to its shorter-lived trajectory T9 and survivors showed continued clot formation and stabilization. For combined trans-omics, 12/13 deaths were patients that began in the same states and followed the same trajectory. Yet similarly injured patients that began in the same states but switched into different omics trajectories survived.

For direct applicability to the clinic, a rapid means of quick state identification and transition propensity can be employed from similar work. This would allow for transomics-driven endotypes to provide prediction and decision support based on multivariate data which would otherwise be too complicated to discern. This then is a more formalistic modeling of experienced clinician gestalt. Future work to refine, rapidly identify states via quick (perhaps targeted field) measurement (with portable devices), dimension reduction, trajectory tracking and prediction is planned and essential to realize personalized trauma medicine by identifying intervenable mechanistic pathways and guiding individual treatment toward the unified goal of saving lives. Future work will center around codifying and clinically understanding the drivers of these discrete trajectories with the ultimate goal of rapid point of care precision medicine guided by state and trajectory identification. This therefore would allow for tailored treatment and clinical guidance of trajectories towards recovery.

Limitations with this study include sample size and sparsity in some outcomes (i.e. mortality). While sample collection was very good for this study there is some natural missingness of measurements based on censoring (i.e. death or discharge) and random missingness due to the harried nature of trauma care. These limitations have been mitigated in part by cross validation with an independent cohort of trauma patients as described in the methods, and will continue to be addressed with more sampling. Furthermore one powerful aspect of this work is the ability to clinically and biologically supervise results, giving clinical context to model outputs. Future studies and resulting data will improve and allow for testing and refinement of our models.

## MATERIALS AND METHODS

*See Supplementary files for extended methods*

### Experimental design

The overall hypothesis for this observational study was that plasma omics molecular biomarker data will give new insights into trauma patient classification, endotypes, and clinical outcome prediction. The objectives of this study were 1) to identify standard S/T omics signatures; and 2) identify trauma patient omics endotypes and trajectories. All patients enrolled in COMBAT with both metabolomic and proteomic data available were included, totaling 118 patients and 97 healthy controls, and none were treated as outliers. Plasma underwent LC-MS/MS proteomics and metabolomics. Blood was used to measure biochemical and coagulation data using standard clinical measurements, viscoelastic hemostatic assays, coagulation factor levels, and cytokine and chemokine levels using ELISA. Endpoints for the study were 168 hours post-injury, or the final timepoint of hospital care regardless of discharge status. For biological and technical replicates, N=1 plasma samples were collected from each patient at each timepoint.

### Proteomics

Workflows for mass spectrometry-based proteomics were used as previously described(*60*). Briefly, plasma samples were tryptic digested using S-Trap 96-well plates. Peptides were lyophilized, resuspended in 0.1% formic acid, and loaded onto Evotips. Peptides were analyzed using the Evosep One system coupled to a timsTOF Pro mass spectrometer (diaPASEF mode) via the nano-electrospray ion source. Raw DIA files were searched in Spectronaut using a project specific spectral library, and data were presented as units of relative intensity.

### Ultra-High-Pressure Liquid Chromatography-Mass Spectrometry metabolomics

Frozen plasma aliquots (10 µL) were extracted 1:25 in ice cold extraction solution (methanol:acetonitrile:water 5:3:2 v/v/v). Samples were vortexed for 30 min at 4℃, prior to centrifugation for 10 min at 15,000g at 4℃. Analyses were performed using a Vanquish UHPLC coupled online to a Q Exactive mass spectrometer (ThermoFisher). Samples were analyzed using a 1 min and 5 minute gradient-based method, spectra were searched in Maven, and data were presented as units of relative intensity.

### Bioinformatics and data processing

Omics variables with constant values were removed, and values at or below the limit of detection were imputed with 20% the minimum value for that analyte. R software and packages (versions and citations in supplemental files) were used for data analysis and generating all graphs. Mfuzz was used for C-means clustering, EnhancedVolcano created volcano plots, and pheatmap generated heatmaps. UMAP was used to create a 2D embedding of the omic manifold, followed by hierarchical clustering to identify patient states. Dual multiple factor analysis was employed on embeddings for hierarchical clustering to identify patient trajectories. The number of clusters was chosen using a combination of percent of variance explained, power effect size (η^2^), the gap statistic, and silhouette method. Metaboanalyst and Metascape were used for pathway enrichment.

### Statistical analysis

Data were log2-transformed to approximate and assume normal distribution for statistical analyses (**Figure S25**). Significant differences were determined as alpha P<0.05 using one-way ANOVA with TukeyHSD post-hoc analysis, Student’s t-test, or Pearson Chi-squared test. Omic comparisons were corrected using the Benjamini-Hochberg FDR method, and shown with median +/- IQR in boxplots or median +/- SEM in timeseries line graphs. Continuous clinical data with P < 0.05 was considered statistically significant, and shown using median +/- IQR. Linear mixed modeling was used to determine associations between analytes and clinical measurements (fixed effects age, sex, and time; random effects S/T group, individual patient); and to identify analytes and clinical measurements that had significant interactions with time per group (group*time; random effect of individual patient). Sidak post-hoc correction was performed on the estimated marginal means of mixed models. The partial slope was used as the primary measurement and for visualization in the circus plots. Ensemble methods for classification were built using SuperLearner with full details in SF10, receiver operating characteristic curves were calculated to show model performance using the area under the curve, and VIP of analytes were calculated using 5x repeated 5-fold cross-validation to avoid overfitting with RandomForest.

## Author contributions

Conceptualization: All authors

Data curation: ISL, CBE, EEM, MJC, KCH, ADA

Formal analysis: ISL, CBE, MJC, KCH, ADA

Funding acquisition: EEM, CCS, KCH, MC, ADA

Investigation: All authors

Methodology: MJC, CBE, ISL, ADA, KCH

Supervision: EEM, CCS, ADA, KCH

Validation: CBE, MJC, KCH, ADA

Visualization: CBE, ISL

Writing – original draft: MJC, CBE, ADA, KCH

Writing – review & editing: All authors

## Data and materials availability

All data are available in **Supplementary Table 1**. Raw Proteomics, Metabolomics data will be made publicly available via Metabolomics Workbench and PRIDE upon acceptance and final publication. Code will be uploaded to Zenodo upon final publication.

