## Supplementary Methods and Figures for "Trans-Omics analysis of post injury thrombo-inflammation identifies endotypes and trajectories in trauma patients"

#### Supplementary Materials and Methods - Extended

*Experimental design:* The overall design is an observational study using data gathered during the COMBAT clinical trial(*1*). The overall hypothesis for this study was that plasma omics data will give new insights into trauma patient classification, endotypes, and clinical outcome prediction. The objectives of this study were 1) to identify omic signatures of trauma patients as classified by standard S/T categories; and 2) identify new trauma patient trajectories as defined by unsupervised clustering of plasma omics. N=150 patients were calculated to be sufficient for statistical power in the COMBAT trial. In this study, all patients enrolled in COMBAT with both metabolomic and proteomic data available were included, totaling 118 patients and 97 healthy controls. Due to heterogeneity of baseline physiology and response to trauma, no patients were treated as outliers. During the COMBAT trial, patients were randomized to pre-hospital treatment with plasma or saline as a control in the field, after which blood was collected for the duration of hospital care at pre-determined timepoints(*1, 2*). Plasma underwent proteomics and metabolomics analysis via liquid chromatography coupled with tandem mass spectrometry (LC-MS/MS). Blood was used to measure biochemical and coagulation data using standard clinical measurements, viscoelastic hemostatic assays, coagulation factor levels, and cytokine and chemokine levels using ELISA. Endpoints for the study were 168 hours post-injury, or the final timepoint of hospital care regardless of discharge status (home, death, long-term care, other). For biological and technical replicates, N=1 plasma samples were collected from each patient at each timepoint.

*Proteomics:* Proteomics: Plasma samples were digested in the S-Trap 96-well plate (Protifi, Huntington, NY) was performed following the manufacturer's procedure. Briefly, around 50 μg of plasma proteins were first mixed with 5% SDS. Samples were reduced with 10 mM DTT at 55 °C for 30 min, cooled to room temperature, and then alkylated with 25 mM iodoacetamide in the dark for 30 min. Samples were reduced with 10 mM DTT at 55 °C for 30 min, cooled to room temperature, and then alkylated with 25 mM iodoacetamide in the dark for 30 min. Next, a final concentration of 1.2% phosphoric acid and then six volumes of binding buffer (90% methanol; 100 mM triethylammonium bicarbonate, TEAB; pH 7.1) were added to each sample. After gentle mixing, the protein solution was loaded to a S-Trap 96-well plate, spun at 1500 x g for 2 min, and the flow-through collected and reloaded onto the 96-well plate. This step was repeated three times, and then the 96-well plate was washed with 200 μL of binding buffer 3 times. Finally, 1 μg of sequencing-grade trypsin (Promega) and 125 μL of digestion buffer (50 mM TEAB) were added onto the filter and digested carried out at 37 °C for 6 hs. To elute peptides, three stepwise buffers were applied, with 200 μL of each with one more repeat, including 50 mM TEAB, 0.2% formic acid in H2O, and 50% acetonitrile and 0.2% formic acid in H2O. The peptide solutions were pooled, lyophilized and resuspended in 500 ul of 0.1 % FA. Those samples were diluted further 10-fold with 0.1 % FA.

A 20 ul of each sample was loaded onto individual Evotips for desalting and then washed 3 times with 200 μL 0.1% FA followed by the addition of 100 μL storage solvent (0.1% FA) to keep the Evotips wet until analysis. The Evosep One system (Evosep, Odense, Denmark) was used to separate peptides on a Pepsep column, (150 um inter diameter, 15 cm) packed with ReproSil C18 1.9 um, 120A resin. The system was coupled to the timsTOF Pro mass spectrometer (Bruker Daltonics, Bremen, Germany) via the nano-electrospray ion source (Captive Spray, Bruker Daltonics).

The mass spectrometer was operated in diaPASEF mode. We used a method with four windows in each 100 ms dia-PASEF scan. 32 of these scans covered the diagonal scan line for doubly and triply charged peptides in the m/z - ion mobility plane with narrow 25 m/z precursor windows.

We used a project-specific library generated from 24 high-pH reverse-phase peptide fractions acquired with PASEF technology. MS data were collected over an m/z range of 100 to 1,700. During each MS/MS data collection, each TIMS cycle was 1.17 seconds and included 1 MS and 10 PASEF MS/MS scans. Low-abundance precursor ions with an intensity above a threshold of 500 counts but below a target value of 20000 counts were repeatedly scheduled and otherwise dynamically excluded for 0.4 min.

Pooled plasma digest was separated by high pH reversed phase chromatography on a Gemini-NH C18, 4.6 x 250 mm analytical column containing 3 µM particles, flow rate was 0.6 mL/min. The solvent consisted of 20 mM ammonium bicarbonate (pH 10) as mobile phase (A) and 20 mM ammonium bicarbonate and 75% ACN (pH 10) as mobile-phase B. Sample separation was accomplished using the following linear gradient: from 0 to 5% B in 10 min, from 5 to 50% B in 80 min, from 50 to 100% B in 10 min, and held at 100% B for an additional 10 min. 96 fractions were collected along with the LC separation and were concatenated into 24 fractions by combining fractions 1, 25, 49, 73 and so on. Samples were dried in Speed-Vac , resuspended in 80 ul of 0.1% FA and 20ul of each fraction was loaded onto individual Evotips. Peaks were searched in Spectronaut at 1% FDR, and proteomics data are presented here in units of relative intensity.

*Ultra-High-Pressure Liquid Chromatography-Mass Spectrometry metabolomics:* Frozen plasma aliquots (10 µL) were extracted 1:25 in ice cold extraction solution (methanol:acetonitrile:water 5:3:2 v/v/v). Samples were vortexed for 30 min at 4°C, prior to centrifugation for 10 min at 15,000g at 4°C, as described(*3, 4*). Analyses were performed using a Vanquish UHPLC coupled online to a Q Exactive mass spectrometer (ThermoFisher). Samples were analyzed using a 1 min and 5 minute gradient-based method, spectra were searched in Maven, and data are presented as units of relative intensity.

*Bioinformatics and data processing:* Omics variables with constant values were removed, and values at or below the limit of detection were imputed with 20% the minimum value for that analyte. R software and packages (versions in supplemental material) were used for data analysis and generating all graphs(*5*). Mfuzz was used for C-means clustering, EnhancedVolcano created volcano plots, and pheatmap generated heatmaps(*6, 7*). UMAP was used to create a 2D embedding of the omic manifold (n_neighbors = 2, n_components = 20, min_dist = 0.3/0.5 metabolomics/proteomics, pca = 20, euclidean), followed by hierarchical clustering to identify patient states(*8*). Dual multiple factor analysis was employed (n_neighbors = 25/15, n_components = 40/100, min_dist = 0.5/0.7 metabolomics/proteomics, pca = 40/100, Euclidean distance) on embeddings for hierarchical clustering to identify patient trajectories(*9, 10*). This is conceptually similar to clustering patient-specific PCAs that has previously been reported to discern clinical inflammatory endotypes in blunt injury trauma patients(*11*). This method was chosen based on the "data table" structure of our longitudinal data, whereby patients had different numbers of timepoints collected for two reasons: 1) Given the hectic nature of trauma care, blood draws were not possible for some patients at some timepoints, and 2) patients were discharged at different times for different reasons. For 6 patients with N=1 datapoint (who usually experienced death at ED), these patients had 1 additional identical 'phantom' datapoint added to meet the model's requirements. For metabolomics patient trajectories, trajectory 1 was split into 2 trajectories, based on their location in the metabolomics meta-states of catabolism or biosynthesis. The number of clusters was chosen using a combination of percent of variance explained, power effect size ($\eta^2$), the gap statistic, and silhouette method(*12, 13*). Metaboanalyst and Metascape (using KEGG, PANTHER, STRING, MCODE, and DisGeNET databases) were used for pathway enrichment(*14-19*).

*R package versions and brief settings:* R statistical software (v4.1.1) and packages (uwot_0.1.11; clValid_0.7; plotly_4.10.0; circlize_0.4.15; dendextend_1.15.2; htmlwidgets_1.5.4; readxl_1.4.0; EnhancedVolcano_1.12.0; ggpubr_0.4.0; factoextra_1.0.7; matrixTests_0.1.9.1; caret_6.0-92; pheatmap_1.0.12; effectsize_0.7.0; NbClust_3.0.1; ggplot2_3.3.6; tidyverse_1.3.1; FactoMineR_2.4, dendextend_1.15.2) were used for statistical and bioinformatic analysis and for generating graphs(*5*).

*Statistical analysis:* Data were log2-transformed to approximate and assume normal distribution for statistical analyses (SF24). Significant differences were determined as $P<0.05$ using one-way ANOVA with TukeyHSD post-hoc analysis, Student's t-test, or Pearson Chi-squared test. Omic comparisons were corrected using the Benjamini-Hochberg FDR method, and shown with median +/- IQR in boxplots or median +/- SEM in timeseries line graphs. Continuous clinical data with $P < 0.05$ was considered statistically significant, and shown using median +/- IQR. Linear mixed modeling was used to determine associations between analytes and clinical measurements (fixed effects age, sex, and time; random effects S/T group, individual patient); and to identify analytes and clinical measurements that had significant interactions with time per group (group*time; random effect of individual patient). Sidak post-hoc correction was performed on the estimated marginal means of mixed models. The partial slope was used as the primary measurement and for visualization in the circus plots. Ensemble methods for classification were built using SuperLearner with full details in SF10, receiver operating characteristic curves were calculated to show model performance using the area under the curve, and VIP of analytes were calculated using 5x repeated 5-fold cross-validation with RandomForest.

#### Supplementary FIGURES

##### *Supplementary FIGURE 1*

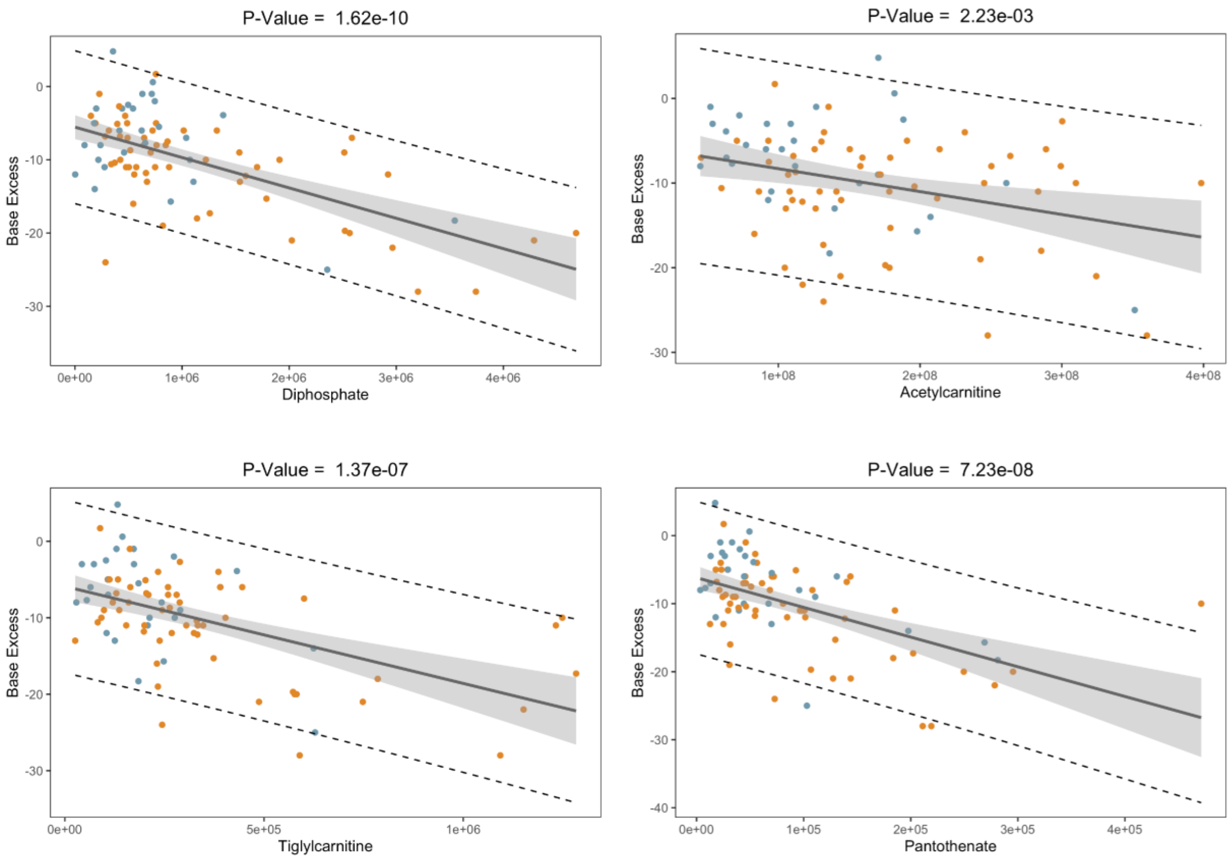

**Supplemental Figure 1 (Figure S1):** Base Excess (BE) linear regression imputation model derived from training on an independent cohort of 333 trauma patients, and used to impute missing BE value at ED for 20 patients.

Linear Regression Equation:

$$Y = -(4.582e\text{-}01)X_{\text{intercept}} - (2.231e\text{-}06)X_{\text{Diphosphate}} - (1.659e\text{-}05)X_{\text{Pantothenate}} - (1.921e\text{-}08)X_{\text{Acetylcarnitine}} - (5.194e\text{-}06)X_{\text{Tiglylcarnitine}}$$

*Supplementary FIGURE 2*

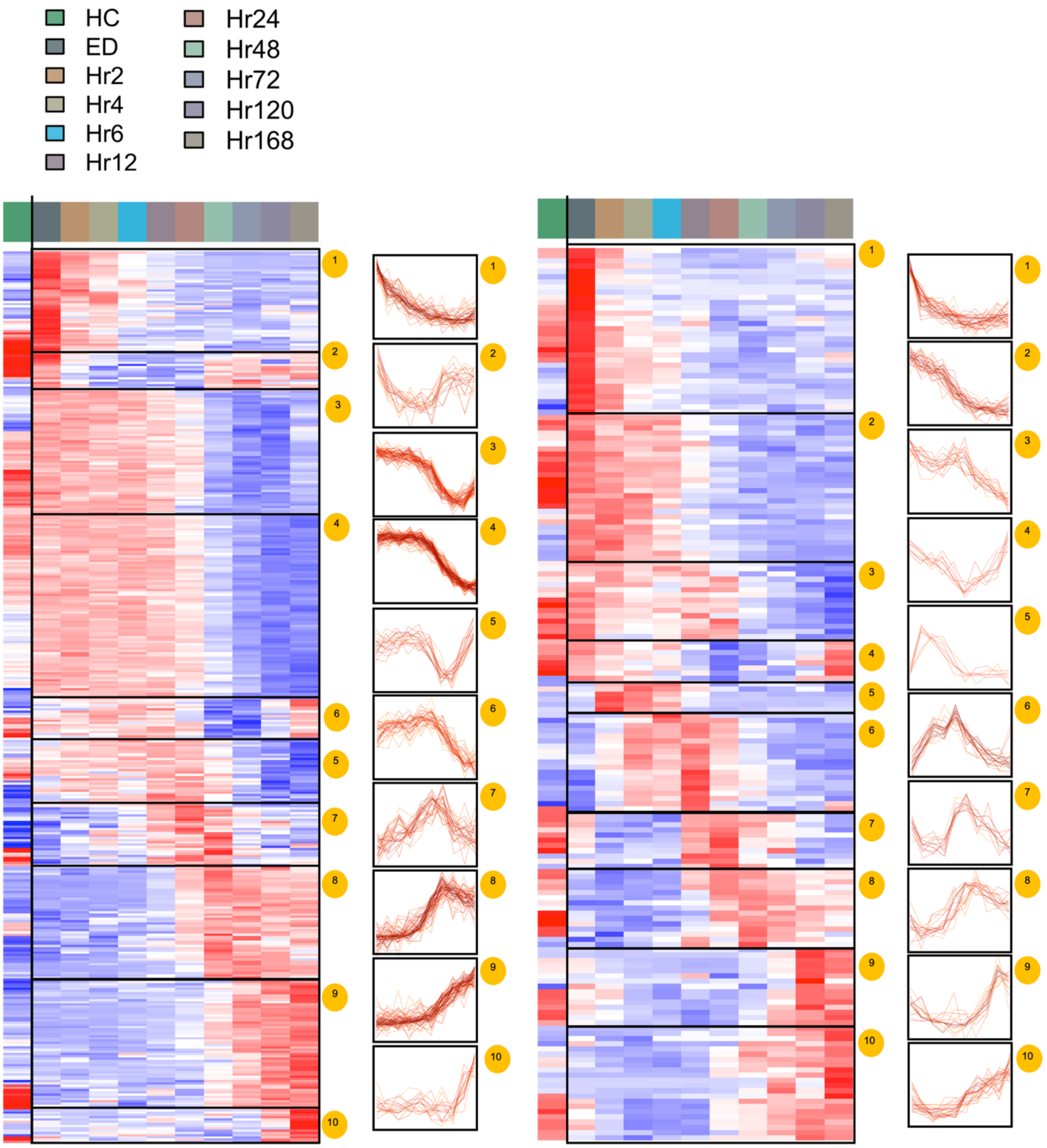

**Supplemental Figure 2 (Figure S2):** C-means clustering results showing line trends of each analyte within its respective cluster. C-means was calculated with 75% zero row threshold using Knn imputation and a minimum membership of 0.6, and were standardized separately into 10 clusters.

*Supplementary FIGURE 3*

| Patient State | Pathway | Total | Expected | p.value |
|---|---|---|---|---|
| 10 | Alanine, aspartate and glutamate metabolism | 28 | 0.52387 | 3.37E-07 |
| 10 | Citrate cycle (TCA cycle) | 20 | 0.37419 | 8.06E-07 |
| 10 | Arginine biosynthesis | 14 | 0.26194 | 2.85E-06 |
| 2 | Arginine and proline metabolism | 38 | 0.29419 | 0.0025694 |
| 2 | Tryptophan metabolism | 41 | 0.31742 | 0.0032042 |
| 2 | Cysteine and methionine metabolism | 33 | 0.25548 | 0.025397 |
| 2 | Thiamine metabolism | 7 | 0.054194 | 0.053051 |
| 9 | Sphingolipid metabolism | 21 | 0.12194 | 0.0059461 |
| 9 | Histidine metabolism | 16 | 0.092903 | 0.08938 |
| 3 | Arginine biosynthesis | 14 | 0.08129 | 0.0026318 |
| 3 | Pantothenate and CoA biosynthesis | 19 | 0.11032 | 0.0048712 |
| 3 | Vitamin B6 metabolism | 9 | 0.052258 | 0.05119 |
| 1 | Taurine and hypotaurine metabolism | 8 | 0.015484 | 0.015414 |
| 1 | Cysteine and methionine metabolism | 33 | 0.063871 | 0.06256 |
| 1 | Fatty acid biosynthesis | 47 | 0.090968 | 0.088292 |
| 6 | Biosynthesis of unsaturated fatty acids | 36 | 0.30194 | 0.0001512 9 |
| 6 | Porphyrin and chlorophyll metabolism | 30 | 0.25161 | 0.024751 |

| | | | | |
|---|---|---|---|---|
| 6 | Pyrimidine metabolism | 39 | 0.3271 | 0.040405 |
| 7 | Glutathione metabolism | 28 | 0.10839 | 0.0045154 |
| 7 | Purine metabolism | 65 | 0.25161 | 0.023294 |
| 7 | Steroid hormone biosynthesis | 85 | 0.32903 | 0.038599 |
| 7 | Pentose phosphate pathway | 22 | 0.085161 | 0.082324 |
| 5 | Tryptophan metabolism | 41 | 0.23806 | 0.0012932 |
| 5 | Tyrosine metabolism | 42 | 0.24387 | 0.022877 |
| 8 | Primary bile acid biosynthesis | 46 | 0.17806 | 0.011982 |
| 8 | Valine, leucine and isoleucine biosynthesis | 8 | 0.030968 | 0.03062 |
| 4 | Aminoacyl-tRNA biosynthesis | 48 | 0.40258 | 5.33E-10 |
| 4 | Phenylalanine, tyrosine and tryptophan biosynthesis | 4 | 0.033548 | 0.00038616 |
| 4 | Glycine, serine and threonine metabolism | 33 | 0.27677 | 0.0021764 |

**Supplemental Figure 3 (Figure S3):** C-means metabolite cluster enrichment analysis, from MetaboAnalyst pathway enrichment workflow.

#### Supplementary FIGURE 4

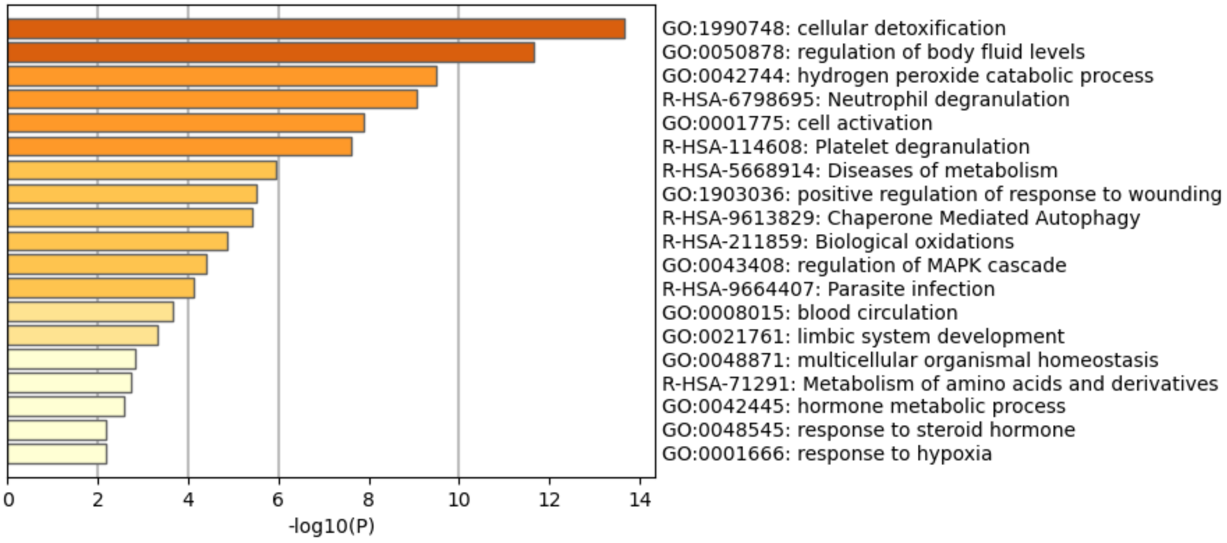

Cluster 1

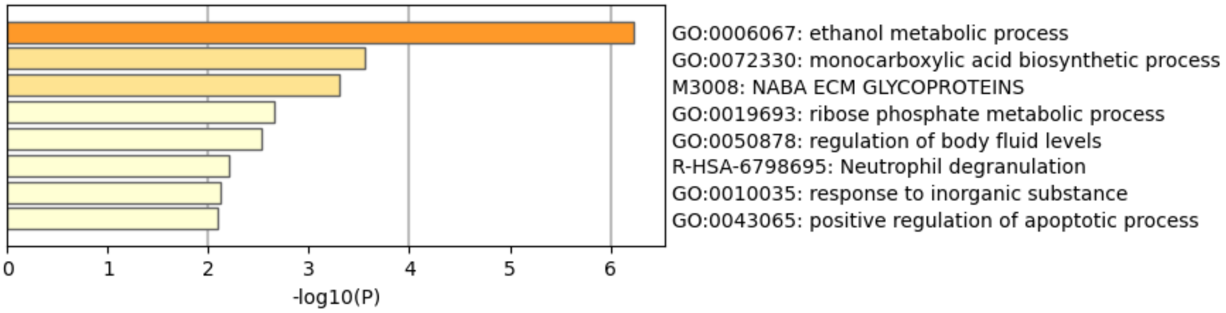

Cluster 2

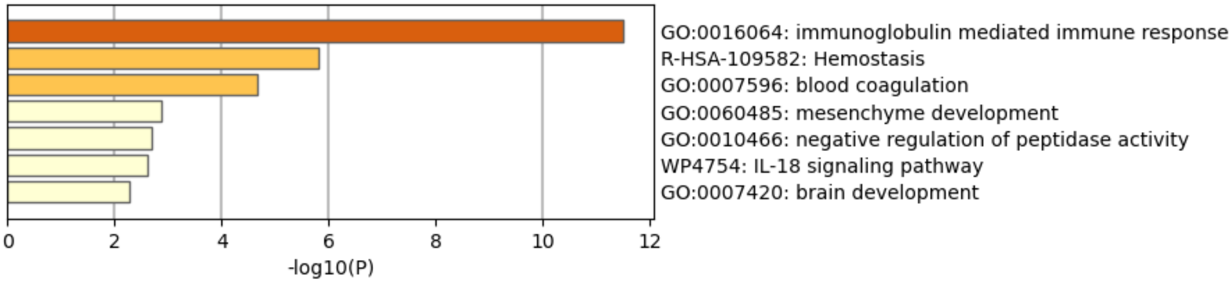

Cluster 3

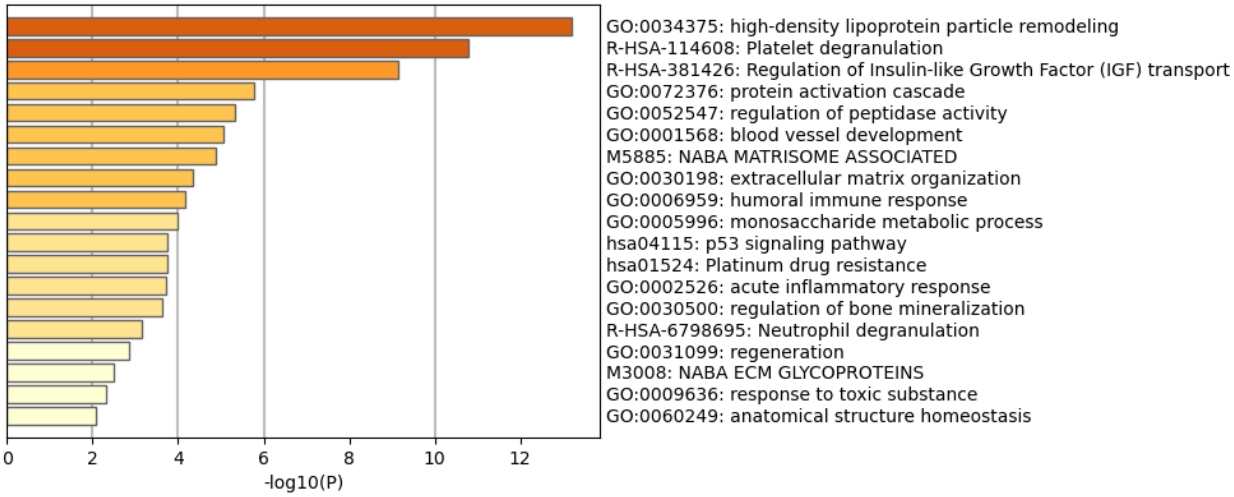

Cluster 4

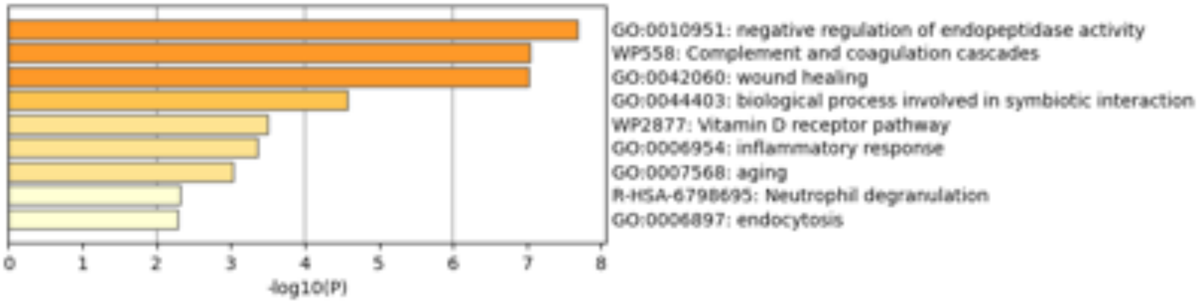

Cluster 5

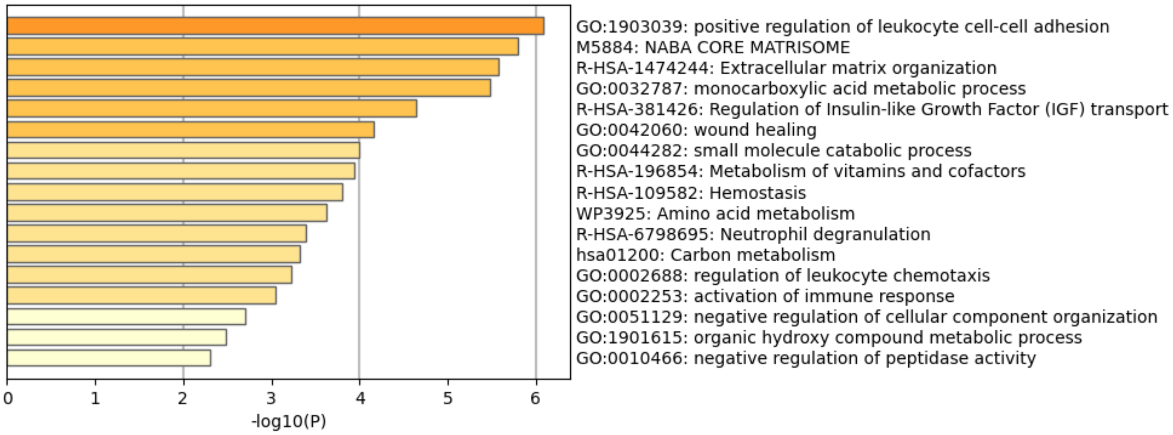

Cluster 6

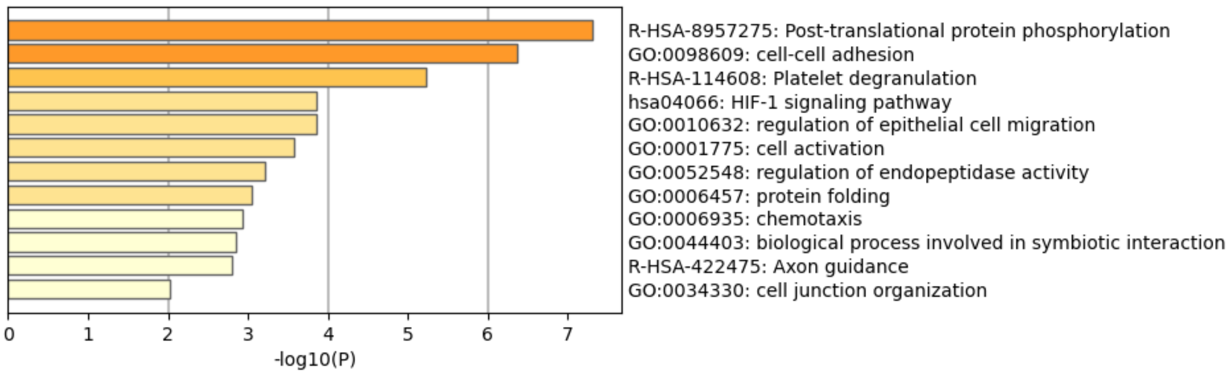

Cluster 7

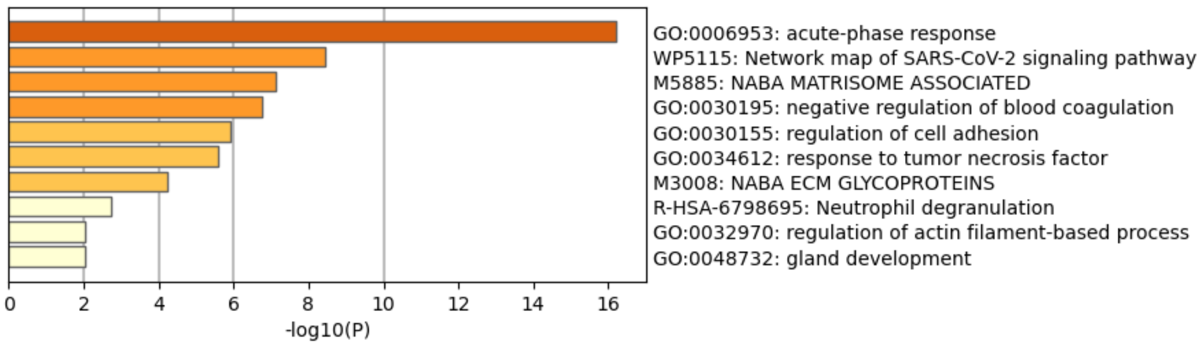

Cluster 8

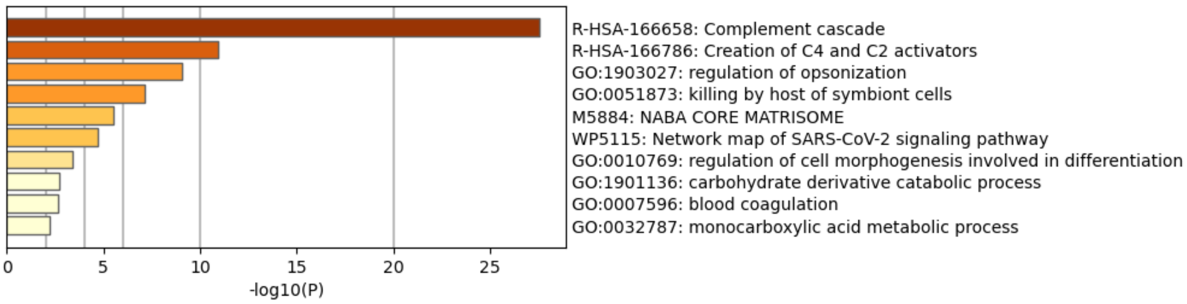

Cluster 9

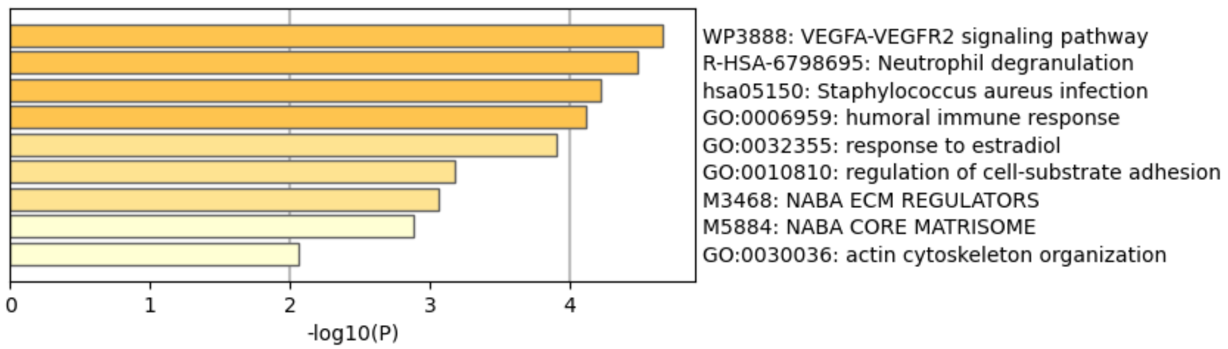

Cluster 10

**Supplemental Figure 4 (Figure S4):** Enrichment results of C-means proteomic clusters, attained from using Metascape.

*Supplementary FIGURE 5*

| ClinicalAssay | Protein | PartialSlope | R2 |
|---|---|---|---|
| SBP | CPN1 | 9.807758294 | 0.37552506 |
| DBP | C2 | 7.46621483 | 0.300786777 |
| Temp | SERPINA3 | 0.508907977 | 0.562514381 |
| HR | LYZ | 6.180419384 | 0.392899347 |
| pH | HRG | 0.07403739 | 0.621905503 |
| pa_CO2 | C3 | 4.731910367 | 0.586635643 |
| pa_O2 | PGLYRP2 | 39.19729917 | 0.355294442 |
| Bicarb | ITIH1 | 2.592738374 | 0.683869556 |
| BE | ITIH4.2 | 2.125017155 | 0.716471273 |
| VCO2 | KNG1.2 | 1.989839474 | 0.650574296 |
| Lactate_labs | AFM | 1.251125139 | 0.781054548 |
| WBC | SERPINA1 | 3.709804775 | 0.65446397 |
| HGB | BCHE | 0.87606852 | 0.658540018 |
| HCT | C4BPB | 2.463795108 | 0.599067697 |
| Potassium | CFB | 0.519380428 | 0.543898388 |
| ion_Calcium | A2M | 0.031982564 | 0.279992627 |
| Plt_count | PROC | 31.01085753 | 0.747229504 |

| ClinicalAssay | Protein | PartialSlope | R2 |
|---|---|---|---|
| INR | COPS4 | 0.261753841 | 0.560201865 |
| PTT | TXNDC17 | 4.514303356 | 0.505034723 |
| Fibrinogen | FGA | 118.2810445 | 0.843529748 |
| DDimer | DEFA1 | 1.790062505 | 0.789567743 |
| cn_r | FGB | 1.044453406 | 0.309035339 |
| cn_angle | F5 | 4.505538993 | 0.123661274 |
| cn_ma | FIBG.2 | 7.15813377 | 0.343068768 |
| cn_ly30 | F2 | 1.751599762 | 0.87536724 |
| cn_ly60 | SERPINF1 | 3.442844416 | 0.740501104 |
| rbc.units | C1QC | 1.753595796 | 0.721085225 |
| plt.units | C1QB | 0.210532437 | 0.307251612 |
| NISS | SAA4 | 0.158860081 | 0.999824392 |

Linear mixed model patial slope and $R^2$ values to value the relatioship (association) between proteomics and clinical measurements.

| ClinicalAssay | Protein | PartialSlope | R2 |
|---|---|---|---|
| SBP | Dodecanoic.acid | 2.740877056 | 0.35249409 |
| DBP | Hexadecanoic.acid | 1.888959464 | 0.306854428 |
| Temp | delta8tetrahydrocannabinol | 0.506565202 | NA |
| HR | DErythrose.4phosphate | 14.27561023 | 0.400294138 |
| pH | Bilirubin | 0.036451062 | 0.680460806 |
| pa_CO2 | N1Acetylspermine | 7.896577566 | 0.461605622 |
| pa_O2 | UDP | 31.45344918 | 0.328147138 |
| Bicarb | Decanoic.acid | 1.050515647 | 0.658553653 |
| BE | Nonanoylcarnitine | 0.695691554 | 0.706403644 |
| VCO2 | LysoPE00204 | 0.630251234 | 0.639263979 |
| Lactate_labs | Alanine | 1.14222235 | 0.779034642 |
| WBC | Testosterone.acetate | 0.751773065 | 0.632235781 |
| HGB | Heptanoic.acid | 0.353572856 | 0.659569797 |
| HCT | OAcetylneuraminic.acid | 0.774869559 | 0.603690463 |
| Potassium | Hexanoic.acid | 0.309662846 | 0.526056374 |
| ion_Calcium | LValine | 0.037907782 | 0.296262643 |
| Plt_count | Urate | 9.111764942 | 0.746025167 |
| INR | N1Acetylspermidine | 0.107672297 | 0.649312369 |

| ClinicalAssay | Protein | PartialSlope | R2 |
|---|---|---|---|
| PTT | Fumarate | 4.088270127 | 0.446432866 |
| Fibrinogen | Leucine | 14.65580898 | 0.803739129 |
| DDimer | Hexanoylcarnitine | 0.791857097 | 0.819133694 |
| cn_r | Tiglylcarnitine | 0.55714281 | 0.328213495 |
| cn_angle | LArginine | 1.601617285 | 0.122642931 |
| cn_ma | NNDimethylglycine | 1.857340736 | 0.319326288 |
| cn_ly30 | ProlylArginine | 2.811001591 | 0.879218046 |
| cn_ly60 | Hexenoylcarnitine | 3.02721969 | 0.699649143 |
| rbc.units | Allantoate | 1.332191719 | 0.714112533 |
| plt.units | sorbic.acid | 0.085635243 | 0.278038354 |
| NISS | Nonanoic.acid | 0.073133094 | 0.999820563 |

Linear mixed model patial slope and $R^2$ values to value the relatioship (association) between metabolomics and clinical measurements.

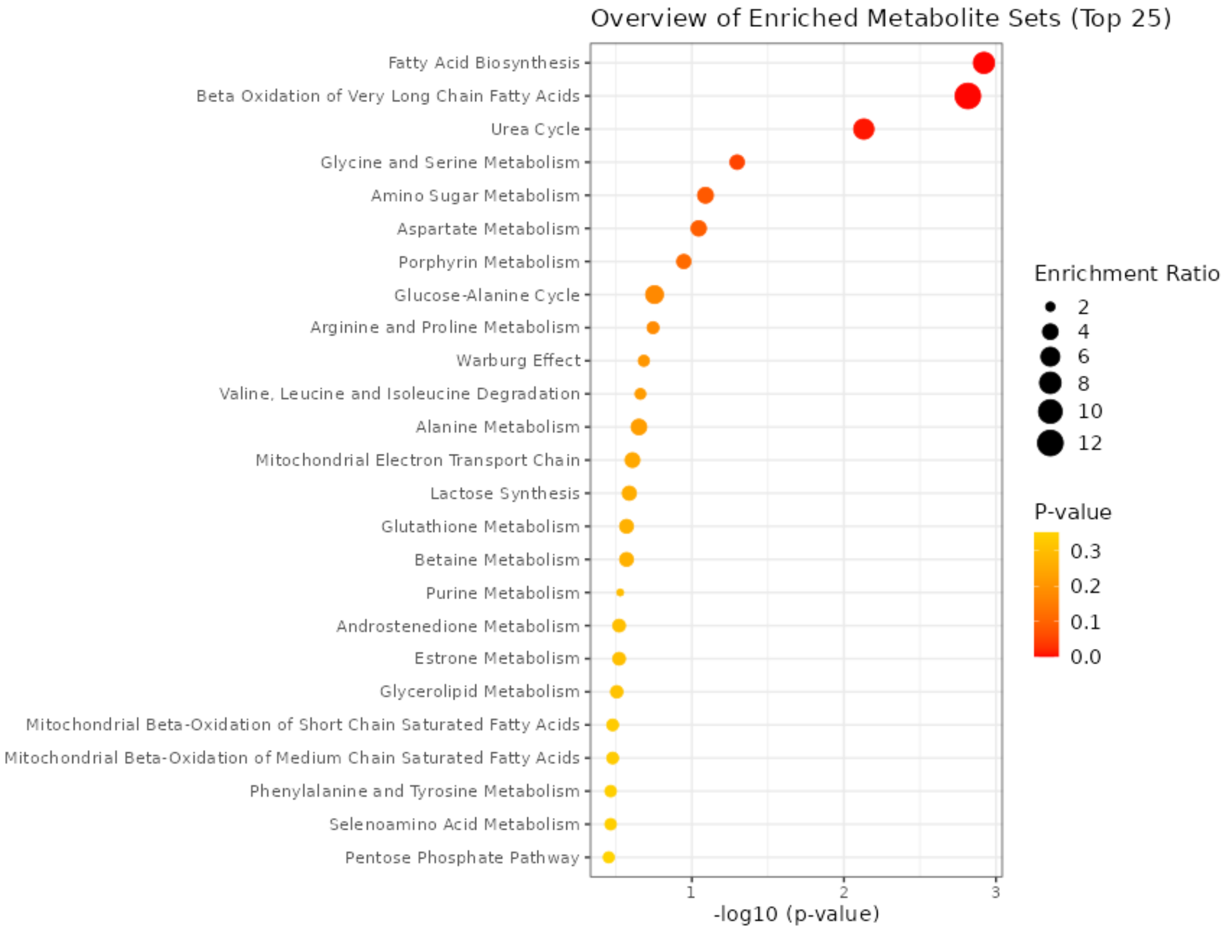

Metaboanalyst pathway enrichment results of highly associated metabolites to identify enrichment terms.

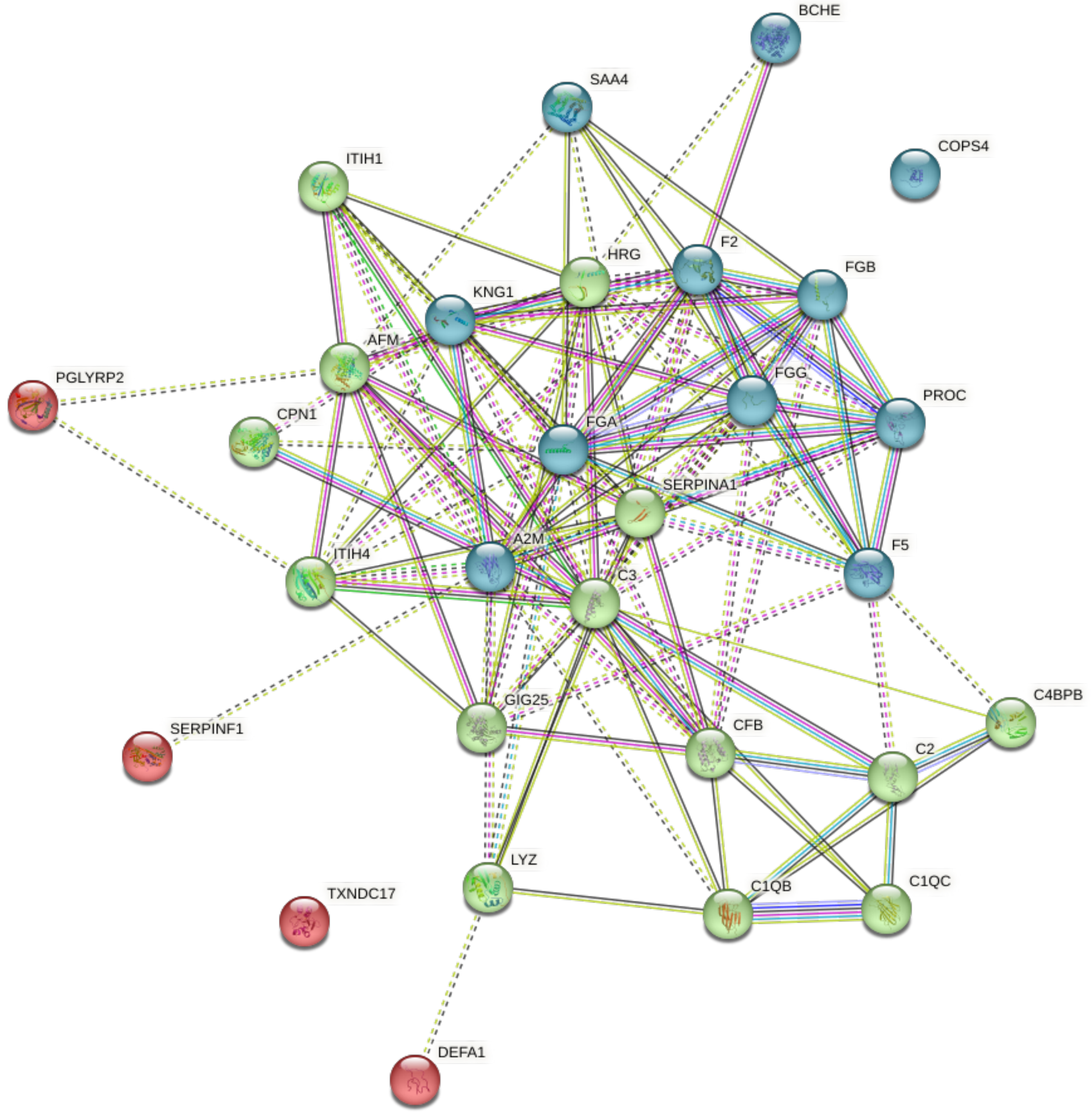

STRING-DB image of K-means clustering (K=3) of highly associated proteins to identify protein-protein interaction enrichment terms.

**Supplemental Figure 5 (Figure S5):** Partial slope and $R^2$ values calculated from linear mixed model used to identify highly associated proteins and metabolites with clinical measurements, as well as enrichment results and network identification using STRING-DB network analysis.

#### Supplementary FIGURE 6

ANOVA results for SF6

| Protein | statistic | pvalue | fdr |
|---|---|---|---|
| FABP1 | 31.8771508 | 1.39E-14 | 1.40E-11 |
| H2AC4 | 25.4415051 | 2.43E-12 | 1.22E-09 |
| GAPDH | 20.8092467 | 1.41E-10 | 3.63E-08 |
| PFN1 | 20.7876203 | 1.44E-10 | 3.63E-08 |
| LGALS1 | 20.1294254 | 2.64E-10 | 5.31E-08 |
| FABP5 | 18.747091 | 9.62E-10 | 1.61E-07 |
| H4-16 | 18.4778667 | 1.24E-09 | 1.79E-07 |
| SOD1 | 17.8136014 | 2.35E-09 | 2.95E-07 |
| YWHAZ | 16.9462627 | 5.45E-09 | 5.15E-07 |
| GSTA1 | 16.9401689 | 5.48E-09 | 5.15E-07 |
| TPI1 | 16.9133233 | 5.63E-09 | 5.15E-07 |
| ADH4 | 16.7444753 | 6.64E-09 | 5.57E-07 |
| ACTA2 | 16.2239816 | 1.11E-08 | 8.60E-07 |
| PRDX6 | 15.9772625 | 1.42E-08 | 1.02E-06 |
| ABHD14B | 15.873681 | 1.57E-08 | 1.03E-06 |
| FABP4 | 15.8373809 | 1.63E-08 | 1.03E-06 |
| ACTB | 15.6412979 | 1.99E-08 | 1.18E-06 |
| ADH1B | 14.9726787 | 3.90E-08 | 2.18E-06 |
| TAGLN | 14.3331413 | 7.51E-08 | 3.84E-06 |

| **Protein** | **statistic** | **pvalue** | **fdr** |
|---|---|---|---|
| CFL1 | 14.3172322 | 7.63E-08 | 3.84E-06 |
| TUBB4A | 14.0785754 | 9.76E-08 | 4.68E-06 |
| LDHA | 14.0113729 | 1.05E-07 | 4.79E-06 |
| ALDH9A1 | 13.8348723 | 1.26E-07 | 5.14E-06 |
| PEBP1 | 13.8250381 | 1.27E-07 | 5.14E-06 |
| MDH1 | 13.8206981 | 1.28E-07 | 5.14E-06 |
| AKR1A1 | 13.5904957 | 1.62E-07 | 6.28E-06 |
| GRHPR | 13.5419998 | 1.71E-07 | 6.36E-06 |
| SERPINB9 | 13.293061 | 2.21E-07 | 7.78E-06 |
| ENO1 | 13.2810963 | 2.24E-07 | 7.78E-06 |
| PAFAH1B2 | 13.0664719 | 2.81E-07 | 9.13E-06 |
| ACTG1 | 13.0661583 | 2.81E-07 | 9.13E-06 |
| FTL | 13.0098403 | 2.98E-07 | 9.39E-06 |
| EIF5A | 12.5907461 | 4.65E-07 | 1.42E-05 |
| GNMT | 12.3051529 | 6.31E-07 | 1.87E-05 |
| ADH1C | 12.2666905 | 6.57E-07 | 1.89E-05 |
| H3-3A | 12.1931425 | 7.11E-07 | 1.99E-05 |
| EEF1A1 | 11.9948428 | 8.81E-07 | 2.40E-05 |
| APRT | 11.6482622 | 1.28E-06 | 3.39E-05 |
| CDH5 | 11.3414234 | 1.79E-06 | 4.62E-05 |

| Protein | statistic | pvalue | fdr |
|---|---|---|---|
| TXN | 11.2230852 | 2.04E-06 | 5.12E-05 |
| ALDOB | 11.1198924 | 2.28E-06 | 5.60E-05 |
| FBP2 | 11.0619342 | 2.43E-06 | 5.82E-05 |
| FKBP1A | 11.0077846 | 2.58E-06 | 6.03E-05 |
| TUBB | 10.7530747 | 3.41E-06 | 7.81E-05 |
| RIDA | 10.6003573 | 4.04E-06 | 9.04E-05 |
| RHOA | 10.3785943 | 5.17E-06 | 0.00011311 |
| RAN | 10.3574961 | 5.29E-06 | 0.00011333 |
| GLO1 | 10.1204309 | 6.89E-06 | 0.00014456 |
| PPIA | 10.0536518 | 7.43E-06 | 0.0001526 |
| ARHGDIA | 9.80299524 | 9.84E-06 | 0.00019696 |
| CRIP1 | 9.79072863 | 9.97E-06 | 0.00019696 |
| SERPINB1 | 9.73025487 | 1.07E-05 | 0.00020678 |
| SELENBP1 | 9.61432053 | 1.22E-05 | 0.00023123 |
| FTCD | 9.43887175 | 1.48E-05 | 0.00027679 |
| ADH1A | 9.36742187 | 1.61E-05 | 0.0002947 |
| LDHB | 9.29225153 | 1.75E-05 | 0.00031524 |
| CAP1 | 9.20110753 | 1.94E-05 | 0.00034355 |
| NME1 | 9.15299839 | 2.05E-05 | 0.00035393 |
| ARF3 | 9.14474007 | 2.07E-05 | 0.00035393 |

| Protein | statistic | pvalue | fdr |
|---|---|---|---|
| RGN | 9.08717987 | 2.21E-05 | 0.00037165 |
| FTH1 | 9.06497978 | 2.27E-05 | 0.00037178 |
| C11orf54 | 9.05813981 | 2.29E-05 | 0.00037178 |
| MB | 8.93641099 | 2.63E-05 | 0.00042053 |
| UBE2N | 8.90118523 | 2.74E-05 | 0.000431 |
| BLVRB | 8.81776828 | 3.01E-05 | 0.00046696 |
| TUBA1B | 8.74624188 | 3.27E-05 | 0.00049925 |
| UBB | 8.71100953 | 3.41E-05 | 0.00051213 |
| CMBL | 8.3624034 | 5.09E-05 | 0.00075447 |
| TAGLN2 | 8.12503764 | 6.71E-05 | 0.00096783 |
| S100A8 | 8.11758866 | 6.77E-05 | 0.00096783 |
| UBE2V1 | 8.11065492 | 6.82E-05 | 0.00096783 |
| CSTB | 8.06190557 | 7.22E-05 | 0.00100524 |
| DDT | 8.05421245 | 7.29E-05 | 0.00100524 |
| HSP90AB1 | 8.02037725 | 7.58E-05 | 0.00103154 |
| ACTBL2 | 7.98737645 | 7.88E-05 | 0.00105771 |
| PCBD1 | 7.96213416 | 8.11E-05 | 0.00106461 |
| SULT2A1 | 7.95924826 | 8.14E-05 | 0.00106461 |
| S100A9 | 7.92008145 | 8.52E-05 | 0.00110013 |
| MPST | 7.90194224 | 8.70E-05 | 0.00110946 |

| Protein | statistic | pvalue | fdr |
|---|---|---|---|
| BHMT2 | 7.77144101 | 0.00010139 | 0.00126245 |
| ACAT2 | 7.77014205 | 0.00010155 | 0.00126245 |
| HSPB1 | 7.71457799 | 0.00010838 | 0.00133096 |
| PSME2 | 7.66924329 | 0.0001143 | 0.00138674 |
| EIF4A1 | 7.40195015 | 0.00015654 | 0.00187661 |
| TKFC | 7.25879814 | 0.00018538 | 0.00219619 |
| BASP1 | 7.15055702 | 0.00021073 | 0.00246745 |
| PSMA4 | 7.08143474 | 0.00022873 | 0.00264746 |
| AKR1C3 | 7.05209974 | 0.00023683 | 0.00271013 |
| NIT2 | 7.01703783 | 0.0002469 | 0.00279363 |
| FAH | 6.99219665 | 0.0002543 | 0.00284537 |
| AKR1C4 | 6.96810057 | 0.00026169 | 0.00289589 |
| QPRT | 6.94826295 | 0.00026794 | 0.00293281 |
| PGK1 | 6.84538947 | 0.00030286 | 0.00326746 |
| CNDP2 | 6.83662341 | 0.00030604 | 0.00326746 |
| PTMA | 6.83059664 | 0.00030825 | 0.00326746 |
| CAPG | 6.76036861 | 0.00033519 | 0.00351603 |
| CRYL1 | 6.70332184 | 0.00035883 | 0.00372516 |
| PSAT1 | 6.62553692 | 0.00039381 | 0.00404662 |
| GNAI2 | 6.6170215 | 0.00039785 | 0.00404678 |

| Protein | statistic | pvalue | fdr |
|---|---:|---:|---:|
| UGP2 | 6.57878043 | 0.00041649 | 0.00419403 |
| ADH5 | 6.52060791 | 0.00044656 | 0.00445235 |
| H1-4 | 6.49283223 | 0.00046169 | 0.00455804 |
| F10 | 6.45712579 | 0.0004819 | 0.00471142 |
| PGAM1 | 6.37479356 | 0.00053201 | 0.00515125 |
| ALDH1A1 | 6.34421679 | 0.00055193 | 0.0052933 |
| ESD | 6.29406185 | 0.00058627 | 0.0055696 |
| CLIC1 | 6.28568279 | 0.00059222 | 0.0055735 |
| PRDX1 | 6.26481313 | 0.00060729 | 0.00566245 |
| ASPD | 6.20193452 | 0.00065511 | 0.00605223 |
| GCHFR | 6.18506335 | 0.00066858 | 0.0061205 |
| HBB | 6.17196634 | 0.00067922 | 0.00616197 |
| EPHX2 | 6.13563267 | 0.00070967 | 0.00638073 |
| AKR7A3 | 6.08604424 | 0.00075348 | 0.00670879 |
| HAAO | 6.07947694 | 0.00075949 | 0.00670879 |
| DPYS | 6.06940055 | 0.00076879 | 0.00673194 |
| FHL1 | 6.026398 | 0.00080983 | 0.00700587 |
| ABRACL | 6.01862636 | 0.00081748 | 0.00700587 |
| ARPC4 | 6.0151268 | 0.00082095 | 0.00700587 |
| ADH6 | 5.99257769 | 0.00084366 | 0.00713918 |

| **Protein** | **statistic** | **pvalue** | **fdr** |
|---|---:|---:|---:|
| HPD | 5.97179358 | 0.00086515 | 0.00726008 |
| PTGR1 | 5.96374686 | 0.00087362 | 0.00727058 |
| BHMT | 5.91896891 | 0.00092233 | 0.00761297 |
| LAP3 | 5.89886993 | 0.00094507 | 0.0077373 |
| MAT1A | 5.79768245 | 0.00106854 | 0.00867754 |
| IGKV1D-8 | 5.77401521 | 0.0010997 | 0.00885919 |
| HBA1 | 5.75970196 | 0.00111899 | 0.00894307 |
| PNPO | 5.65198532 | 0.00127564 | 0.01011474 |
| PSMA3 | 5.58304707 | 0.00138739 | 0.01091487 |
| QDPR | 5.55688844 | 0.00143235 | 0.0111812 |
| HAGH | 5.53981756 | 0.00146248 | 0.01132859 |
| HINT1 | 5.4040128 | 0.00172626 | 0.01326978 |
| PBLD | 5.35152478 | 0.00184069 | 0.01404226 |
| GPD1 | 5.32939016 | 0.00189123 | 0.01431931 |
| PSMA7 | 5.23907989 | 0.00211241 | 0.01587461 |
| GAMT | 5.22625125 | 0.00214589 | 0.01600674 |
| YWHAQ | 5.16207304 | 0.00232161 | 0.01719019 |
| PRDX2 | 5.15586978 | 0.00233935 | 0.0171951 |
| DPYD | 5.11589978 | 0.00245699 | 0.01792893 |
| GSTM3 | 5.03355663 | 0.00271858 | 0.01969503 |

| Protein | statistic | pvalue | fdr |
|---|---|---|---|
| BDH2 | 4.99089272 | 0.00286503 | 0.02053705 |
| ENO3 | 4.98789986 | 0.00287559 | 0.02053705 |
| MAN2A1 | 4.97161763 | 0.00293378 | 0.02080507 |
| PSMB6 | 4.96096664 | 0.00297249 | 0.02093216 |
| RAB1A | 4.93268438 | 0.0030778 | 0.02152324 |
| ARG1 | 4.89541948 | 0.00322235 | 0.02237865 |
| APOC3 | 4.88648383 | 0.00325802 | 0.02247139 |
| ROBO4 | 4.88031257 | 0.00328288 | 0.02248887 |
| CAPNS1 | 4.85493758 | 0.00338717 | 0.02304648 |
| MMP9 | 4.83914773 | 0.00345375 | 0.02327138 |
| GSTP1 | 4.83451848 | 0.00347352 | 0.02327138 |
| IGF1 | 4.83078273 | 0.00348955 | 0.02327138 |
| CD163 | 4.80918271 | 0.00358376 | 0.02374238 |
| DEFA1 | 4.7238395 | 0.00398182 | 0.02620713 |
| CTH | 4.71449243 | 0.00402805 | 0.02633925 |
| ASS1 | 4.69601015 | 0.00412106 | 0.02670242 |
| CA1 | 4.69295984 | 0.00413662 | 0.02670242 |
| CUL3 | 4.66691734 | 0.0042719 | 0.02740001 |
| ACOC | 4.65424385 | 0.00433934 | 0.02765643 |
| PSMA1 | 4.63902241 | 0.00442176 | 0.02800451 |

| Protein | statistic | pvalue | fdr |
|---|---|---|---|
| HBD | 4.61795718 | 0.00453845 | 0.02856388 |
| DBI | 4.60316384 | 0.00462225 | 0.02891061 |
| H2AJ | 4.59251306 | 0.00468355 | 0.02911319 |
| MASP2 | 4.54451069 | 0.00497019 | 0.03070542 |
| PSMB5 | 4.48876037 | 0.00532543 | 0.03269946 |
| COL18A1 | 4.48218709 | 0.00536898 | 0.03276702 |
| KHK | 4.4395477 | 0.00566029 | 0.03433682 |
| CA3 | 4.41903634 | 0.00580606 | 0.0350102 |
| H3C13 | 4.40075165 | 0.0059392 | 0.03559986 |
| PARK7 | 4.37264619 | 0.00614989 | 0.03664462 |
| WARS1 | 4.28142314 | 0.00688718 | 0.0407964 |
| PGM3 | 4.27212567 | 0.00696717 | 0.04102886 |
| GPX4 | 4.25817022 | 0.00708899 | 0.04150355 |
| GDI1 | 4.24961347 | 0.00716475 | 0.04170461 |
| AMDHD1 | 4.2427398 | 0.00722619 | 0.04182055 |
| AKR1C2 | 4.21953738 | 0.00743756 | 0.04279787 |
| PREP | 4.2114705 | 0.00751251 | 0.04298348 |
| DCXR | 4.20110224 | 0.00760995 | 0.04329501 |
| PSMB1 | 4.19268414 | 0.00769 | 0.04350465 |
| EEF1G | 4.1767793 | 0.00784357 | 0.04412556 |

| Protein | statistic | pvalue | fdr |
|---|---|---|---|
| HSP90AA1 | 4.16090496 | 0.00799993 | 0.04465031 |
| GSR | 4.15833646 | 0.00802553 | 0.04465031 |
| HDGF | 4.09438317 | 0.00869012 | 0.04766223 |
| LASP1 | 4.09378019 | 0.00869665 | 0.04766223 |
| IGLC7 | 4.09265001 | 0.00870889 | 0.04766223 |
| SDC1 | 4.07861573 | 0.00886234 | 0.04823987 |
| SLC9A3R1 | 4.05637247 | 0.00911115 | 0.04926577 |
| SORD | 4.05307282 | 0.00914866 | 0.04926577 |
| PRG4 | 4.04018006 | 0.0092967 | 0.04979669 |

Significantly different proteins among S/T groups at ED

| Protein | conf.level | Ctrlmean | Exptmean | FC | fdrpval | logFC |
|---|---|---|---|---|---|---|
| QPRT | 0.95 | 0.74127121 | 21.4146147 | 28.8890416 | 0.00552575 | 4.85245044 |
| AHNAK | 0.95 | 8.20287534 | 175.461075 | 21.3901916 | 0.02145991 | 4.4188775 |
| RGN | 0.95 | 0.89766994 | 12.7494664 | 14.2028442 | 0.00022454 | 3.82810796 |
| GSTA1 | 0.95 | 45.7677291 | 616.866041 | 13.4781876 | 1.52E-06 | 3.75255461 |
| WARS1 | 0.95 | 1.89279219 | 24.7710661 | 13.0870501 | 0.00483791 | 3.71006803 |
| RHOA | 0.95 | 1.54843517 | 19.687697 | 12.7145762 | 2.14E-05 | 3.66841147 |
| GNMT | 0.95 | 3.67868463 | 40.1325783 | 10.9094914 | 4.62E-06 | 3.44751194 |
| H3-3A | 0.95 | 2.22546089 | 22.8573952 | 10.2708591 | 1.52E-06 | 3.36048495 |
| GLO1 | 0.95 | 2.45394427 | 25.1064084 | 10.2310426 | 2.30E-05 | 3.35488127 |
| UBE2N | 0.95 | 1.21326763 | 12.3522222 | 10.1809542 | 0.0018069 | 3.34780088 |
| AKR1C3 | 0.95 | 6.66609652 | 67.685463 | 10.1536878 | 0.00065871 | 3.3439319 |
| SULT2A1 | 0.95 | 17.3887553 | 168.581088 | 9.69483353 | 0.00041969 | 3.27721613 |
| ALDH9A1 | 0.95 | 5.84857704 | 54.9161298 | 9.38965656 | 1.03E-05 | 3.23107239 |
| PSME2 | 0.95 | 2.69231946 | 25.0230207 | 9.29422421 | 0.00031999 | 3.21633445 |
| ADH4 | 0.95 | 109.00895 | 1002.97764 | 9.20087429 | 3.59E-06 | 3.20177096 |
| RIDA | 0.95 | 13.024172 | 119.594726 | 9.18252046 | 2.42E-05 | 3.1988902 |
| FABP1 | 0.95 | 107.028639 | 900.574306 | 8.41433015 | 1.44E-08 | 3.07284843 |
| H2AC4 | 0.95 | 45.725066 | 380.039211 | 8.31139777 | 1.62E-10 | 3.05509112 |
| CAPG | 0.95 | 4.80220829 | 39.2340848 | 8.17000898 | 0.00028416 | 3.03033766 |
| ADH1A | 0.95 | 72.3596775 | 573.929115 | 7.93161516 | 0.00022532 | 2.98761468 |

| Protein | conf.level | Ctrlmean | Exptmean | FC | fdrpval | logFC |
|---|---|---|---|---|---|---|
| ABHD14B | 0.95 | 7.04063836 | 55.5940977 | 7.89617288 | 8.76E-07 | 2.98115358 |
| FBP2 | 0.95 | 16.9322395 | 131.00875 | 7.73723702 | 0.00012447 | 2.95181847 |
| ADH1C | 0.95 | 21.3348464 | 164.450547 | 7.70807268 | 2.80E-06 | 2.94637018 |
| TUBB4A | 0.95 | 6.75028727 | 51.7499514 | 7.6663332 | 4.62E-06 | 2.9385367 |
| QDPR | 0.95 | 1.65214046 | 12.6509636 | 7.65731722 | 0.00240904 | 2.93683902 |
| PAFAH1B2 | 0.95 | 2.32208763 | 17.701339 | 7.62302798 | 2.80E-06 | 2.93036417 |
| PNPO | 0.95 | 1.39974553 | 10.6323963 | 7.59594946 | 0.00164934 | 2.92523031 |
| ADH1B | 0.95 | 287.403164 | 2173.57596 | 7.56281153 | 9.52E-07 | 2.91892267 |
| FABP5 | 0.95 | 9.02708623 | 65.0672873 | 7.20800551 | 4.10E-08 | 2.84960011 |
| DDT | 0.95 | 45.151302 | 324.086838 | 7.17779607 | 0.00031095 | 2.84354093 |
| HAGH | 0.95 | 3.54441342 | 25.3443232 | 7.1504986 | 0.00487098 | 2.83804384 |
| AKR1C2 | 0.95 | 7.20656587 | 50.3810415 | 6.99099162 | 0.01194659 | 2.8054971 |
| GCHFR | 0.95 | 3.42900362 | 23.9642621 | 6.98869547 | 0.00125394 | 2.80502318 |
| RAB1A | 0.95 | 14.4208887 | 97.5537227 | 6.7647511 | 0.01206943 | 2.75803685 |
| ARHGDIA | 0.95 | 8.27617828 | 55.0561654 | 6.65236581 | 4.99E-05 | 2.7338675 |
| ABRACL | 0.95 | 2.64828715 | 17.5247075 | 6.61737437 | 0.00059704 | 2.7262589 |
| HINT1 | 0.95 | 1.11916248 | 7.34101937 | 6.55938657 | 0.00301486 | 2.7135609 |
| C11orf54 | 0.95 | 12.0325 | 76.3847166 | 6.34820002 | 0.00036599 | 2.66634758 |
| EIF4A1 | 0.95 | 1.90904724 | 12.0878589 | 6.33188047 | 0.01609781 | 2.66263402 |
| CD93 | 0.95 | 52.1666579 | 328.434876 | 6.29587726 | 0.01807405 | 2.65440742 |

| Protein | conf.level | Ctrlmean | Exptmean | FC | fdrpval | logFC |
|---|---|---|---|---|---|---|
| H2AJ | 0.95 | 1.15381274 | 7.1463326 | 6.19366763 | 0.00697086 | 2.63079397 |
| PDIA6 | 0.95 | 0.69591541 | 4.28942602 | 6.16371758 | 0.0333729 | 2.62380076 |
| SERPINB9 | 0.95 | 4.04962461 | 24.8788097 | 6.14348542 | 4.45E-06 | 2.61905738 |
| EIF5A | 0.95 | 6.58626944 | 40.0207636 | 6.07639331 | 6.77E-06 | 2.60321526 |
| PTMA | 0.95 | 3.34626349 | 19.9293538 | 5.9557037 | 0.00059704 | 2.57427198 |
| PSMA4 | 0.95 | 4.63701678 | 26.6355591 | 5.74411531 | 0.00209257 | 2.52208471 |
| PSMB1 | 0.95 | 11.8439458 | 67.3389048 | 5.68551275 | 0.01423853 | 2.50729046 |
| H4-16 | 0.95 | 28.3144012 | 158.977885 | 5.61473589 | 5.33E-08 | 2.48921816 |
| GDI1 | 0.95 | 1.38996937 | 7.70019641 | 5.53983173 | 0.02981027 | 2.46984215 |
| GSTZ1 | 0.95 | 2.94856226 | 16.1307997 | 5.47073395 | 0.02151633 | 2.4517344 |
| CRIP1 | 0.95 | 4.90332877 | 26.8148147 | 5.46869605 | 1.30E-05 | 2.45119688 |
| CMBL | 0.95 | 14.5762933 | 79.3373068 | 5.44289999 | 0.00103744 | 2.44437553 |
| DCXR | 0.95 | 56.5019095 | 301.749385 | 5.34051658 | 0.01976388 | 2.4169793 |
| HAAO | 0.95 | 5.30866803 | 28.1044564 | 5.29406929 | 0.01444406 | 2.40437708 |
| TXN | 0.95 | 11.3241819 | 56.5098768 | 4.99019507 | 0.0001589 | 2.31909621 |
| FABP4 | 0.95 | 31.8615093 | 158.647093 | 4.97927111 | 1.43E-07 | 2.31593457 |
| CMPK1 | 0.95 | 2.82313795 | 13.9931506 | 4.95659471 | 0.02270563 | 2.3093493 |
| ADH5 | 0.95 | 17.2601634 | 85.1302848 | 4.93218302 | 0.00059704 | 2.30222634 |
| PTGR1 | 0.95 | 15.074711 | 74.1795845 | 4.92079647 | 0.00500148 | 2.29889185 |
| H3C13 | 0.95 | 0.90454187 | 4.42459057 | 4.89152655 | 0.00994986 | 2.29028477 |

| **Protein** | **conf.level** | **Ctrlmean** | **Exptmean** | **FC** | **fdrpval** | **logFC** |
|---|---|---|---|---|---|---|
| APRT | 0.95 | 6.77074439 | 32.9617872 | 4.86826637 | 0.00095726 | 2.28340811 |
| MDH1 | 0.95 | 15.6648808 | 74.6992098 | 4.76857824 | 6.13E-07 | 2.25355919 |
| ALDOB | 0.95 | 152.06315 | 722.870629 | 4.75375283 | 0.00028087 | 2.24906689 |
| GRHPR | 0.95 | 19.9345298 | 94.7252257 | 4.75181641 | 5.64E-06 | 2.2484791 |
| PEBP1 | 0.95 | 85.6275118 | 397.261939 | 4.6394194 | 1.01E-05 | 2.21394427 |
| GPX1 | 0.95 | 3.98119537 | 18.4131159 | 4.62502193 | 0.02231308 | 2.20946021 |
| ARG1 | 0.95 | 19.3200221 | 89.0706852 | 4.61027864 | 0.00563608 | 2.20485395 |
| ADH6 | 0.95 | 22.4926207 | 102.664649 | 4.56437028 | 0.00119112 | 2.19041583 |
| CORO1A | 0.95 | 0.79786985 | 3.59539485 | 4.50624229 | 0.01265563 | 2.17192489 |
| FAH | 0.95 | 36.7065722 | 164.57549 | 4.48354287 | 0.00301486 | 2.16463919 |
| AKR1C4 | 0.95 | 15.1913427 | 67.404287 | 4.4370197 | 0.00584329 | 2.14959096 |
| FTL | 0.95 | 63.7761998 | 282.305771 | 4.42650663 | 8.41E-06 | 2.14616858 |
| SOD1 | 0.95 | 66.2347225 | 292.610211 | 4.41777666 | 5.00E-08 | 2.14332049 |
| GAMT | 0.95 | 8.71085962 | 38.0399049 | 4.36695189 | 0.01776464 | 2.12662664 |
| H1-4 | 0.95 | 6.28012951 | 27.1832907 | 4.32846021 | 0.00049329 | 2.1138539 |
| GAPDH | 0.95 | 196.925274 | 851.131574 | 4.32210431 | 4.10E-08 | 2.11173389 |
| FTCD | 0.95 | 47.7850762 | 205.778758 | 4.30633943 | 0.00014081 | 2.10646204 |
| PSMA3 | 0.95 | 11.8238401 | 49.955139 | 4.2249505 | 0.01755125 | 2.07893444 |
| SLC9A3R1 | 0.95 | 2.46744143 | 10.4149904 | 4.22096762 | 0.04633129 | 2.07757376 |
| PRDX6 | 0.95 | 52.0648135 | 218.323592 | 4.19330403 | 6.64E-08 | 2.06808744 |

| Protein | conf.level | Ctrlmean | Exptmean | FC | fdrpval | logFC |
|---|---|---|---|---|---|---|
| CTH | 0.95 | 7.70967819 | 32.3243919 | 4.19270314 | 0.01928759 | 2.06788068 |
| GNAI2 | 0.95 | 2.44149935 | 9.93926258 | 4.07096671 | 0.01683856 | 2.02537142 |
| NIT2 | 0.95 | 13.9699063 | 56.7822728 | 4.06461372 | 0.00174303 | 2.02311825 |
| RPLP2 | 0.95 | 2.21034812 | 8.94259082 | 4.04578389 | 0.04050639 | 2.01641926 |
| PSAT1 | 0.95 | 11.360153 | 45.1268002 | 3.97237611 | 0.00037728 | 1.99000223 |
| FHL1 | 0.95 | 4.19973144 | 16.5216348 | 3.93397412 | 0.0008705 | 1.97598747 |
| TPI1 | 0.95 | 52.0049214 | 204.177063 | 3.92611041 | 6.57E-07 | 1.97310074 |
| MMP9 | 0.95 | 7.05047992 | 27.5419138 | 3.90638851 | 0.01683856 | 1.96583544 |
| PFN1 | 0.95 | 69.2307955 | 269.987502 | 3.89981799 | 2.52E-08 | 1.96340679 |
| HPD | 0.95 | 43.5527893 | 166.896132 | 3.83204232 | 0.00091966 | 1.93811349 |
| YWHAZ | 0.95 | 13.9171784 | 53.3166299 | 3.83099421 | 4.19E-08 | 1.93771885 |
| TAGLN | 0.95 | 15.7458921 | 60.1666793 | 3.82110324 | 9.52E-07 | 1.93398924 |
| GPT | 0.95 | 47.6981569 | 180.222357 | 3.77839247 | 0.03949754 | 1.91777257 |
| CLTC | 0.95 | 1.33069736 | 5.02154751 | 3.77362101 | 0.03380315 | 1.91594954 |
| ACSS2 | 0.95 | 0.75919339 | 2.85927738 | 3.76620425 | 0.04249003 | 1.91311124 |
| CKB | 0.95 | 0.77530246 | 2.91216987 | 3.75617263 | 0.03827796 | 1.90926337 |
| TUBB | 0.95 | 30.8323779 | 115.167246 | 3.73526966 | 0.0001589 | 1.9012124 |
| PTPA | 0.95 | 3.21398429 | 11.9836023 | 3.72858149 | 0.02863737 | 1.89862687 |
| CFL1 | 0.95 | 24.4533352 | 90.2960256 | 3.69258529 | 1.52E-06 | 1.88463124 |
| UBE2V1 | 0.95 | 15.5129881 | 56.878881 | 3.66653288 | 0.00407715 | 1.87441648 |

| Protein | conf.level | Ctrlmean | Exptmean | FC | fdrpval | logFC |
|---|---|---|---|---|---|---|
| SHMT1 | 0.95 | 37.5949667 | 137.818847 | 3.66588559 | 0.00584329 | 1.87416176 |
| LASP1 | 0.95 | 2.31644497 | 8.40299201 | 3.62753794 | 0.02332434 | 1.8589907 |
| CAPNS1 | 0.95 | 3.56808179 | 12.6970256 | 3.55850184 | 0.00407715 | 1.83126998 |
| TKFC | 0.95 | 32.9291422 | 116.366672 | 3.53385071 | 0.00065656 | 1.82124109 |
| AKR7A3 | 0.95 | 20.6215978 | 72.4407564 | 3.51285855 | 0.02242957 | 1.81264549 |
| SELENBP1 | 0.95 | 60.6874874 | 212.533904 | 3.50210418 | 2.14E-05 | 1.808222 |
| SLC2A1 | 0.95 | 19.0423841 | 65.729 | 3.45172116 | 0.04056442 | 1.78731592 |
| IGLV5-52 | 0.95 | 2.85629088 | 9.83530124 | 3.44338223 | 0.0333729 | 1.78382633 |
| GPX4 | 0.95 | 1.87545689 | 6.4347752 | 3.43104405 | 0.03493722 | 1.77864765 |
| LGALS1 | 0.95 | 23.0289736 | 78.7324062 | 3.41884131 | 2.36E-09 | 1.77350746 |
| LDHA | 0.95 | 56.6141909 | 191.22559 | 3.37769714 | 4.62E-06 | 1.75603997 |
| ACTG1 | 0.95 | 9.43100189 | 31.7271442 | 3.36413295 | 1.32E-06 | 1.75023472 |
| ALDH1A1 | 0.95 | 107.646253 | 361.393769 | 3.357235 | 0.00066743 | 1.74727352 |
| CDC42 | 0.95 | 4.55895826 | 15.2598426 | 3.3472214 | 0.0224256 | 1.74296398 |
| CLIC1 | 0.95 | 11.6927265 | 38.6506104 | 3.3055259 | 0.00174275 | 1.72487982 |
| PCBD1 | 0.95 | 12.3903337 | 40.6584381 | 3.28146433 | 0.0001645 | 1.71433975 |
| EEF1G | 0.95 | 17.7143491 | 57.6872777 | 3.25652822 | 0.00543858 | 1.70333473 |
| RAN | 0.95 | 21.7600646 | 70.6134456 | 3.24509356 | 7.61E-05 | 1.69826007 |
| KHK | 0.95 | 6.23918601 | 19.912284 | 3.19148747 | 0.00742951 | 1.67422898 |
| AMDHD1 | 0.95 | 12.0350089 | 38.041293 | 3.16088616 | 0.0333729 | 1.66032908 |

| Protein | conf.level | Ctrlmean | Exptmean | FC | fdrpval | logFC |
|---|---|---|---|---|---|---|
| ENO1 | 0.95 | 42.4386214 | 133.575423 | 3.14749675 | 7.92E-07 | 1.65420489 |
| PSMA1 | 0.95 | 7.35327735 | 23.0769934 | 3.13832762 | 0.02145991 | 1.64999597 |
| NDRG2 | 0.95 | 12.2077383 | 37.5464044 | 3.07562329 | 0.03293887 | 1.62087881 |
| FTH1 | 0.95 | 15.0125569 | 45.5644051 | 3.03508625 | 2.29E-05 | 1.60173751 |
| BASP1 | 0.95 | 1.50121731 | 4.51457662 | 3.00727722 | 0.00382758 | 1.58845787 |
| BHMT2 | 0.95 | 86.4100352 | 257.486219 | 2.97981848 | 0.00061961 | 1.57522445 |
| CTRB1 | 0.95 | 6.8094391 | 19.934641 | 2.92750118 | 0.01976388 | 1.54966975 |
| EPHX2 | 0.95 | 13.2880359 | 37.8537313 | 2.84870778 | 0.00277611 | 1.51030764 |
| CSTB | 0.95 | 17.8050023 | 49.4702809 | 2.77844844 | 0.0002458 | 1.47427947 |
| TUBA1B | 0.95 | 48.0587547 | 133.272528 | 2.77311655 | 0.00012475 | 1.47150825 |
| ACY1 | 0.95 | 53.857116 | 146.261125 | 2.71572515 | 0.01755125 | 1.44133748 |
| HSPB1 | 0.95 | 19.5851176 | 52.6045512 | 2.68594513 | 0.00474995 | 1.42542983 |
| YWHAQ | 0.95 | 10.5801414 | 28.3808476 | 2.68246392 | 0.00284624 | 1.42355877 |
| CRYZ | 0.95 | 20.7297828 | 55.1325156 | 2.65958 | 0.01656255 | 1.41119843 |
| ACTBL2 | 0.95 | 110.831869 | 288.559061 | 2.6035748 | 7.10E-05 | 1.38049386 |
| BPGM | 0.95 | 35.86588 | 92.0650397 | 2.56692544 | 0.00640561 | 1.36004139 |
| HNRNPD | 0.95 | 5.39354248 | 13.8183249 | 2.5620128 | 0.01482556 | 1.35727768 |
| UGDH | 0.95 | 15.2684179 | 38.8478754 | 2.5443288 | 0.02559911 | 1.34728512 |
| CAP1 | 0.95 | 7.05384128 | 17.4213209 | 2.46976367 | 0.00020015 | 1.304373 |
| AKR1A1 | 0.95 | 85.5654899 | 208.112757 | 2.43220435 | 3.31E-06 | 1.28226445 |

| Protein | conf.level | Ctrlmean | Exptmean | FC | fdrpval | logFC |
|---|---|---|---|---|---|---|
| FKBP1A | 0.95 | 51.6269555 | 125.511434 | 2.43112212 | 2.77E-05 | 1.28162236 |
| TAGLN2 | 0.95 | 280.08485 | 679.285251 | 2.42528381 | 0.00037519 | 1.27815358 |
| MAT1A | 0.95 | 33.2306543 | 79.9586301 | 2.4061708 | 0.00095726 | 1.26673906 |
| ACAT2 | 0.95 | 7.36460329 | 17.2129093 | 2.33724867 | 0.00099876 | 1.22481124 |
| MPST | 0.95 | 22.7690051 | 53.0560643 | 2.33018808 | 0.00040863 | 1.22044641 |
| HSPA8 | 0.95 | 77.0169961 | 179.267735 | 2.32763862 | 0.0261578 | 1.21886709 |
| ACTA2 | 0.95 | 246.263028 | 570.580629 | 2.31695611 | 1.60E-07 | 1.21223072 |
| DSP | 0.95 | 21.182178 | 48.2706468 | 2.27883302 | 0.02559911 | 1.18829522 |
| ACTB | 0.95 | 272.460795 | 619.333614 | 2.27311094 | 6.13E-07 | 1.1846681 |
| GSTM3 | 0.95 | 13.4026961 | 30.4278892 | 2.27028122 | 0.00882105 | 1.18287101 |
| GOT2 | 0.95 | 7.90547084 | 17.9023039 | 2.26454619 | 0.01014149 | 1.17922197 |
| ARPC4 | 0.95 | 9.80804305 | 21.5144429 | 2.19355103 | 0.00125394 | 1.13326827 |
| CNDP2 | 0.95 | 18.5071862 | 39.7973495 | 2.15037279 | 0.00284624 | 1.10458679 |
| BHMT | 0.95 | 78.2233905 | 168.065141 | 2.14852795 | 0.00709787 | 1.10334855 |
| PPIA | 0.95 | 89.3900771 | 188.315909 | 2.10667576 | 1.16E-05 | 1.07496829 |
| SORD | 0.95 | 70.3789143 | 147.855658 | 2.10085165 | 0.03632372 | 1.07097429 |
| SPYA | 0.95 | 31.8343404 | 65.9621517 | 2.07204393 | 0.049082 | 1.05105459 |
| UGP2 | 0.95 | 34.6529558 | 71.7985447 | 2.07193133 | 0.00826312 | 1.05097619 |
| GSTT1 | 0.95 | 12.5260192 | 25.7741298 | 2.05764731 | 0.04545826 | 1.04099572 |
| DPYS | 0.95 | 18.5207796 | 38.0725236 | 2.05566528 | 0.00149952 | 1.03960537 |

| Protein | conf.level | Ctrlmean | Exptmean | FC | fdrpval | logFC |
|---|---|---|---|---|---|---|
| FAHD2A | 0.95 | 21.2038639 | 43.5586548 | 2.05427911 | 0.02720771 | 1.03863221 |
| ESD | 0.95 | 40.5674006 | 81.2780229 | 2.00353046 | 0.00065871 | 1.00254445 |
| PRDX2 | 0.95 | 97.3059673 | 194.836696 | 2.00230984 | 0.00233629 | 1.00166524 |

**Proteins significantly elevated in the high trauma groups at ED**

| Protein | conf.level | Ctrlmean | Exptmean | FC | fdrpval | logFC |
|---|---|---|---|---|---|---|
| HSPA4 | 0.95 | 81.0776296 | 35.1032845 | 0.43295894 | 0.02224003 | -1.2076979 |
| TYMP | 0.95 | 63.9567187 | 18.8319788 | 0.2944488 | 0.03380315 | -1.7639113 |
| MSN | 0.95 | 390.744134 | 109.42112 | 0.28003266 | 0.04056442 | -1.836333 |

**Proteins significantly elevated in the low trauma groups at ED**

| Protein | conf.level | Ctrlmean | Exptmean | FC | fdrpval | logFC |
|---|---|---|---|---|---|---|
| GSTA1 | 0.95 | 101.652322 | 669.412354 | 6.58531296 | 0.02113389 | 2.719252 |
| FABP1 | 0.95 | 187.009875 | 970.713561 | 5.1907075 | 0.00010937 | 2.37593119 |
| ADH4 | 0.95 | 221.086072 | 1054.87581 | 4.77133542 | 0.01359904 | 2.25439311 |
| FBP2 | 0.95 | 31.4349802 | 137.383349 | 4.37039719 | 0.03406824 | 2.1277644 |
| PEBP1 | 0.95 | 107.98284 | 435.97971 | 4.03749066 | 0.03406824 | 2.01345892 |
| TXN | 0.95 | 16.5104174 | 59.7238192 | 3.61734156 | 0.03406824 | 1.85492983 |
| AKR7A3 | 0.95 | 22.5993574 | 81.0254698 | 3.58529972 | 0.03720925 | 1.84209373 |
| ALDOB | 0.95 | 223.27671 | 756.438465 | 3.38789686 | 0.02113389 | 1.76038995 |
| ARF3 | 0.95 | 31.1255926 | 92.42095 | 2.96929126 | 0.03406824 | 1.57011862 |
| GAPDH | 0.95 | 306.542374 | 855.052237 | 2.78934435 | 0.03406824 | 1.47992605 |
| TPI1 | 0.95 | 79.7653479 | 202.296567 | 2.53614598 | 0.03406824 | 1.34263779 |
| LDHA | 0.95 | 80.0373525 | 190.961136 | 2.3859002 | 0.03478692 | 1.2545337 |
| TUBB | 0.95 | 50.7851265 | 108.488294 | 2.13622179 | 0.04713769 | 1.09506144 |

**Proteins elevated in the HS groups at ED**

| Protein | conf.level | Ctrlmean | Exptmean | FC | fdrpval | logFC |
|---|---|---|---|---|---|---|
| ROBO4 | 0.95 | 64.4827078 | 22.6181644 | 0.35076325 | 0.03406824 | -1.5114305 |

**Proteins elevated in the LS groups at ED**

SERPINA1

| contrast | estimate | SE | df | lower.CL | upper.CL | p.value |
|---|---|---|---|---|---|---|
| HSHT - HSLT | 0.16813789 | 0.06264722 | 110.509589 | 0.00471912 | 0.33155666 | 0.04121545 |
| HSHT - LSHT | 0.0180864 | 0.05541827 | 112.908821 | -0.1264282 | 0.162601 | 0.98793441 |
| HSHT - LSLT | 0.1850268 | 0.05392821 | 131.256026 | 0.04469215 | 0.32536145 | 0.00442182 |
| HSLT - LSHT | -0.1500515 | 0.06512191 | 110.161412 | -0.3199338 | 0.01983086 | 0.10330381 |
| HSLT - LSLT | 0.01688891 | 0.06385867 | 122.274092 | -0.1494447 | 0.1832225 | 0.99349173 |
| LSHT - LSLT | 0.1669404 | 0.05678415 | 128.421296 | 0.01913183 | 0.31474898 | 0.02009683 |

FGA

| contrast | estimate | SE | df | lower.CL | upper.CL | p.value |
|---|---|---|---|---|---|---|
| HSHT - HSLT | 0.19508798 | 0.07675093 | 109.122925 | -0.0051601 | 0.39533603 | 0.05913762 |
| HSHT - LSHT | 0.06450724 | 0.06800531 | 110.710201 | -0.1128835 | 0.24189794 | 0.77868643 |
| HSHT - LSLT | 0.25812273 | 0.06713013 | 135.37278 | 0.08350181 | 0.43274366 | 0.00104991 |
| HSLT - LSHT | -0.1305807 | 0.0797659 | 108.989142 | -0.338699 | 0.07753753 | 0.36233577 |
| HSLT - LSLT | 0.06303475 | 0.07902109 | 125.642819 | -0.1427163 | 0.26878576 | 0.85535693 |
| LSHT - LSLT | 0.1936155 | 0.07055742 | 132.299252 | 0.01002615 | 0.37720484 | 0.03446728 |

S100A9

| contrast | estimate | SE | df | lower.CL | upper.CL | p.value |
|---|---|---|---|---|---|---|
| HSHT - HSLT | 0.90859049 | 0.20055642 | 110.583164 | 0.38543324 | 1.43174774 | 8.739E-05 |
| HSHT - LSHT | 0.2511347 | 0.17739196 | 113.012997 | -0.2114451 | 0.7137145 | 0.49224284 |
| HSHT - LSLT | 1.14159437 | 0.17248397 | 130.992202 | 0.69273639 | 1.59045235 | 5.0708E-09 |

| | | | | | | |
|---|---|---|---|---|---|---|
| HSLT - LSHT | -0.6574558 | 0.20848269 | 110.226267 | -1.2013159 | -0.1135957 | 0.01102761 |
| HSLT - LSLT | 0.23300388 | 0.20432289 | 122.080729 | -0.2992105 | 0.7652183 | 0.66534242 |
| LSHT - LSLT | 0.89045967 | 0.18163941 | 128.180163 | 0.41764243 | 1.36327691 | 1.6594E-05 |

**Timeseries Sidak correction on estimated marginal means of mixed model**

**Supplemental Figure 6 (Figure S6):** Statistical results for main Figure 3 proteomics results: ANOVA with TukeyHSD post-hoc results comparing S/T groups at ED; Volcano results comparing HS vs. LS and HT vs. LT at ED; linear mixed model results for timeseries graphs.

#### *Supplementary FIGURE 7*

| **Metabolite** | **statistic** | **pvalue** | **fdr** |
|---|---|---|---|
| sn-Glycerol.3-phosphate | 21.1075833 | 1.08E-10 | 4.54E-08 |
| Xanthine | 19.0854925 | 6.99E-10 | 1.47E-07 |
| Spermidine | 17.4877898 | 3.21E-09 | 4.51E-07 |
| Creatine | 15.5428878 | 2.19E-08 | 2.31E-06 |
| Succinate | 14.9658775 | 3.93E-08 | 3.31E-06 |
| N1-Acetylspermidine | 13.4775807 | 1.82E-07 | 1.28E-05 |
| Phosphocreatine | 12.603237 | 4.59E-07 | 2.76E-05 |
| Spermine | 12.2988529 | 6.35E-07 | 3.34E-05 |
| Amoxicillin | 12.1345724 | 7.58E-07 | 3.54E-05 |
| Pantothenate | 11.8144439 | 1.07E-06 | 4.50E-05 |
| Tiglylcarnitine | 11.5224304 | 1.47E-06 | 5.62E-05 |
| Taurine | 11.2056932 | 2.07E-06 | 6.83E-05 |
| Hypoxanthine | 11.1904557 | 2.11E-06 | 6.83E-05 |
| D-Glucose.6-phosphate | 10.6982569 | 3.62E-06 | 0.000109 |
| Gerberinol | 10.572807 | 4.16E-06 | 0.00011688 |
| Nicotinamide | 9.60919795 | 1.22E-05 | 0.00032209 |
| Mannitol | 9.31600684 | 1.71E-05 | 0.00042258 |
| sn-glycero-3-Phosphoethanolamine | 8.97830577 | 2.51E-05 | 0.00057107 |
| Hydroxyisovaleroyl.carnitine | 8.95439856 | 2.58E-05 | 0.00057107 |

| Metabolite | statistic | pvalue | fdr |
|---|---|---|---|
| furosemide | 8.42099327 | 4.76E-05 | 0.00100214 |
| Propionylcarnitine | 7.75751908 | 0.00010306 | 0.00206611 |
| Diphosphate | 7.63316358 | 0.00011925 | 0.00214823 |
| Sucrose | 7.61363482 | 0.00012201 | 0.00214823 |
| Choline | 7.61048257 | 0.00012246 | 0.00214823 |
| L-Glutamate | 7.5685618 | 0.00012865 | 0.00216646 |
| Orthophosphate | 7.33603398 | 0.00016921 | 0.00273982 |
| Malate | 7.1266803 | 0.00021678 | 0.00338009 |
| Withaperuvin.E | 6.90646568 | 0.00028161 | 0.00423419 |
| IMP | 6.31320792 | 0.00057292 | 0.00831715 |
| Dehydroascorbate | 6.21896847 | 0.00064179 | 0.00900645 |
| Hexenoylcarnitine | 5.96625343 | 0.00087098 | 0.01182843 |
| Butyrylcarnitine | 5.72237113 | 0.00117094 | 0.01540523 |
| S-Adenosyl-L-homocysteine | 5.40787432 | 0.00171813 | 0.02191916 |
| Putrescine | 5.24476913 | 0.00209773 | 0.02597488 |
| D-Ribose | 5.11765312 | 0.00245171 | 0.02949057 |
| (+)-Quercitol | 5.08543586 | 0.00255066 | 0.02982859 |
| Fumarate | 4.97377286 | 0.00292601 | 0.03329329 |
| Lactate | 4.76529962 | 0.00378314 | 0.04176296 |
| Butanoic.acid | 4.74716507 | 0.00386878 | 0.04176296 |

| Metabolite | statistic | pvalue | fdr |
|---|---|---|---|
| D-Arabitol | 4.61440465 | 0.00455843 | 0.04797752 |

#### Significantly different metabolites among S/T groups at ED

| Metabolite | Ctrlmean | Exptmean | FC | pval | fdrpval |
|---|---|---|---|---|---|
| Withaperuvin.E | 1861.67634 | 26181.4006 | 14.0633472 | 0.0006733 | 0.01771614 |
| sn-glycero-3-Phosphoethanolamine | 17033.267 | 139035.996 | 8.16261473 | 0.00048296 | 0.01355503 |
| sn-Glycerol.3-phosphate | 150709.632 | 1096326.25 | 7.27442717 | 7.39E-08 | 1.55E-05 |
| D-Glucose.6-phosphate | 61985.7276 | 376272.216 | 6.07030409 | 2.99E-06 | 0.00020982 |
| Mannitol | 6664732.02 | 36647408.5 | 5.49870699 | 0.00081473 | 0.01905551 |
| Succinate | 1460903.99 | 7215451.93 | 4.93903224 | 3.99E-05 | 0.00167886 |
| Spermine | 57369.3782 | 273773.507 | 4.77211913 | 1.23E-05 | 0.00064889 |
| Spermidine | 74946.9971 | 326553.262 | 4.35712269 | 1.99E-07 | 2.79E-05 |
| Creatine | 31250185.2 | 101584681 | 3.25069053 | 1.87E-08 | 7.88E-06 |
| Phosphocreatine | 21688.0429 | 67053.3791 | 3.09172106 | 1.98E-06 | 0.00016711 |
| Sucrose | 245399.677 | 685120.228 | 2.79185464 | 4.57E-05 | 0.00174966 |
| Gerberinol | 1059624.56 | 2838585.7 | 2.67885987 | 1.89E-05 | 0.00088313 |
| Malate | 1977491.96 | 5137243.94 | 2.59785832 | 0.00101495 | 0.0224892 |
| Taurine | 239976.539 | 542997.954 | 2.26271267 | 1.16E-06 | 0.0001226 |
| Xanthine | 818126.992 | 1724898.85 | 2.10835099 | 0.00159334 | 0.03194265 |
| Amoxicillin | 155458.298 | 323673.489 | 2.0820599 | 4.63E-06 | 0.00027872 |

**Metabolites elevated in HT at ED**

| Metabolite | Ctrlmean | Exptmean | FC | pval | fdrpval |
|---|---|---|---|---|---|
| furosemide | 48299.4875 | 5351365.71 | 110.795497 | 6.08E-05 | 0.00197019 |
| Withaperuvin.E | 3620.47191 | 29185.3814 | 8.06120919 | 0.00112187 | 0.01628648 |
| Mannitol | 5704661.17 | 44211469.7 | 7.75006058 | 4.35E-05 | 0.0018297 |
| Succinate | 1186392.76 | 8778582.38 | 7.39938972 | 1.89E-06 | 0.00011353 |
| Amoxicillin | 67086.6796 | 468518.293 | 6.98377526 | 0.00012709 | 0.00347471 |
| Gerberinol | 597523.697 | 3787340.2 | 6.3383933 | 0.00021312 | 0.00498465 |
| Lactate | 77659951.9 | 438528411 | 5.64677675 | 0.0007181 | 0.01259668 |
| (+)-Quercitol | 67807.2186 | 341548.611 | 5.03705385 | 0.00236315 | 0.02763575 |
| Hypoxanthine | 1428435.36 | 6863248.42 | 4.80473155 | 6.45E-08 | 7.65E-06 |
| sn-Glycerol.3-phosphate | 269108.586 | 1151411.92 | 4.2786146 | 4.65E-07 | 3.26E-05 |
| Xanthine | 544450.821 | 2255557.08 | 4.14281141 | 2.03E-10 | 8.54E-08 |
| Spermine | 75908.1252 | 296939.336 | 3.91182545 | 5.66E-05 | 0.00197019 |
| sn-glycero-3-Phosphoethanolamine | 40056.9467 | 136581.823 | 3.4096913 | 0.00356754 | 0.03754831 |
| Malate | 1830036.7 | 5991497.67 | 3.27397679 | 0.0021333 | 0.02641533 |
| Tokinolide.A | 54310.4694 | 165772.23 | 3.05230707 | 0.00107316 | 0.01628648 |
| Spermidine | 111559.447 | 334905.188 | 3.00203342 | 7.24E-06 | 0.00038114 |
| Propionylcarnitine | 3599337.82 | 10293953.7 | 2.85995764 | 0.0002375 | 0.00526248 |
| Pantothenate | 47640.9221 | 133482.639 | 2.80184836 | 6.98E-08 | 7.65E-06 |

| **Metabolite** | **Ctrlmean** | **Exptmean** | **FC** | **pval** | **pval10** |
|---|---|---|---|---|---|
| Hydroxypropionylcarnitine | 24490.5347 | 62875.5555 | 2.56734107 | 0.00050817 | 0.00972453 |
| Tiglylcarnitine | 190371.459 | 485743.635 | 2.55155704 | 7.27E-08 | 7.65E-06 |
| Diphosphate | 645447.052 | 1619418.82 | 2.50898786 | 0.00059118 | 0.01082119 |
| D-Ribose | 1119022.65 | 2637323.28 | 2.35680956 | 0.00032898 | 0.00659518 |
| Creatine | 47151346.6 | 96926757.5 | 2.05565195 | 0.00109916 | 0.01628648 |

**Metabolites elevated in HS AT ED**

| **Metabolite** | **Ctrlmean** | **Exptmean** | **FC** | **pval** | **pval10** |
|---|---|---|---|---|---|
| LPC(28:0) | 54698.7223 | 22377.129 | 0.40909784 | 0.00399383 | 2.39860996 |
| IMP | 47568.711 | 8499.11084 | 0.17867019 | 0.00082235 | 3.08494592 |

**Metabolites elevated in LT at ED**

L-Threonine

| contrast | estimate | SE | df | lower.CL | upper.CL | p.value |
|---|---|---|---|---|---|---|
| HSHT - HSLT | -0.2101661 | 0.09895753 | 105.284219 | -0.4684987 | 0.04816654 | 0.15229041 |
| HSHT - LSHT | -0.0961575 | 0.08770533 | 103.513854 | -0.3251788 | 0.1328638 | 0.69254362 |
| HSHT - LSLT | -0.4309785 | 0.08977433 | 141.608763 | -0.6643738 | -0.1975833 | 2.3438E-05 |
| HSLT - LSHT | 0.11400857 | 0.1028751 | 105.934895 | -0.1545245 | 0.38254165 | 0.685286 |
| HSLT - LSLT | -0.2208124 | 0.1046446 | 133.056097 | -0.4930763 | 0.05145145 | 0.15518864 |
| LSHT - LSLT | -0.334821 | 0.0940751 | 138.834523 | -0.5794558 | -0.0901862 | 0.00283919 |

Spermidine

| contrast | estimate | SE | df | lower.CL | upper.CL | p.value |
|---|---|---|---|---|---|---|
| HSHT - HSLT | 0.83216502 | 0.29435649 | 105.264921 | 0.06373324 | 1.60059681 | 0.02833719 |
| HSHT - LSHT | 0.79775906 | 0.26088342 | 103.475556 | 0.11652078 | 1.47899734 | 0.01483082 |
| HSHT - LSLT | 1.06313117 | 0.26708543 | 141.626569 | 0.36876359 | 1.75749876 | 0.00062743 |
| HSLT - LSHT | -0.034406 | 0.30601076 | 105.919623 | -0.8331823 | 0.76437041 | 0.99948839 |
| HSLT - LSLT | 0.23096615 | 0.31131504 | 133.087309 | -0.5790096 | 1.04094192 | 0.87995054 |
| LSHT - LSLT | 0.26537211 | 0.2798776 | 138.855561 | -0.4624259 | 0.99317008 | 0.77887359 |

LPE(18:1)

| contrast | estimate | SE | df | lower.CL | upper.CL | p.value |
|---|---|---|---|---|---|---|
| HSHT - HSLT | -0.4532173 | 0.20276949 | 109.747176 | -0.9822091 | 0.07577451 | 0.12025381 |
| HSHT - LSHT | -0.2112576 | 0.17955879 | 111.746564 | -0.6795673 | 0.25705203 | 0.64287121 |
| HSHT - LSLT | -1.0127891 | 0.17613828 | 133.697551 | -1.471037 | -0.5545412 | 3.4505E-07 |
| HSLT - LSHT | 0.24195965 | 0.21074853 | 109.507043 | -0.3078668 | 0.79178605 | 0.66064589 |
| HSLT - LSLT | -0.5595718 | 0.20784195 | 124.184907 | -1.1008255 | -0.0183182 | 0.03981252 |
| LSHT - LSLT | -0.8015315 | 0.18526784 | 130.692951 | -1.2836714 | -0.3193916 | 0.00017371 |

Creatine

| contrast | estimate | SE | df | lower.CL | upper.CL | p.value |
|---|---|---|---|---|---|---|
| HSHT - HSLT | 0.69840116 | 0.25886601 | 110.723538 | 0.02315475 | 1.37364757 | 0.0397255 |
| HSHT - LSHT | 0.22949874 | 0.22890957 | 113.206936 | -0.3674067 | 0.82640416 | 0.74820739 |

| | | | | | | |
|---|---|---|---|---|---|---|
| HSHT - LSLT | 1.2259864 | 0.22223232 | 130.47269 | 0.64763771 | 1.80433508 | 1.0638E-06 |
| HSLT - LSHT | -0.4689024 | 0.26910707 | 110.35103 | -1.1708986 | 0.2330938 | 0.30692377 |
| HSLT - LSLT | 0.52758524 | 0.26345062 | 121.706787 | -0.1586726 | 1.2138431 | 0.19270829 |
| LSHT - LSLT | 0.99648766 | 0.23408162 | 127.7076 | 0.38713072 | 1.60584459 | 0.00023082 |

**Timeseries Sidak correction on estimated marginal means of mixed model**

**Supplemental Figure 7 (Figure S7):** Statistical results for main Figure 3 for metabolomics: ANOVA with TukeyHSD post-hoc results comparing S/T groups at ED; Volcano results comparing HS vs. LS and HT vs. LT at ED; linear mixed model results for timeseries graphs.

*Supplementary FIGURE 8*

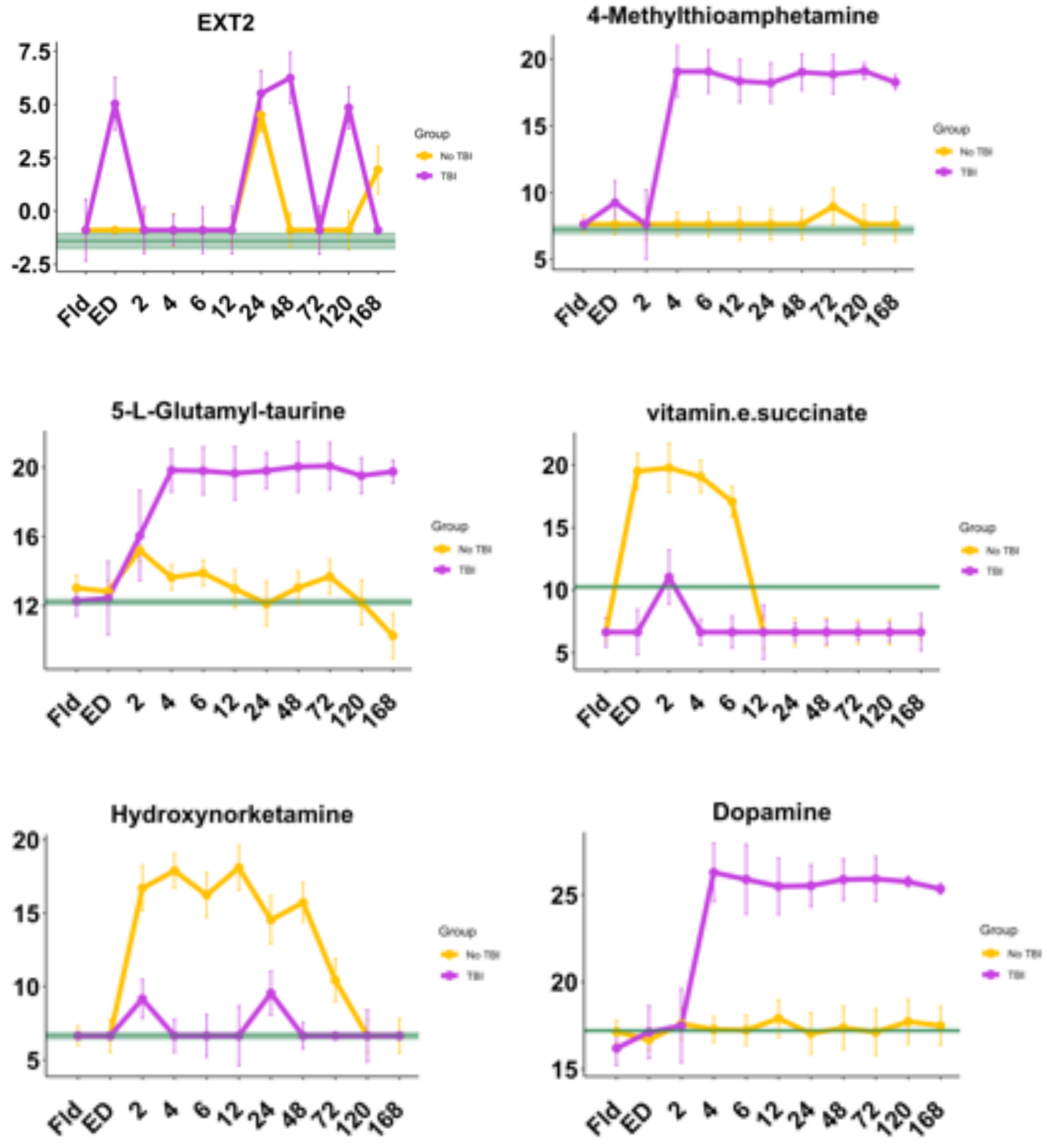

TBI in HSHT group

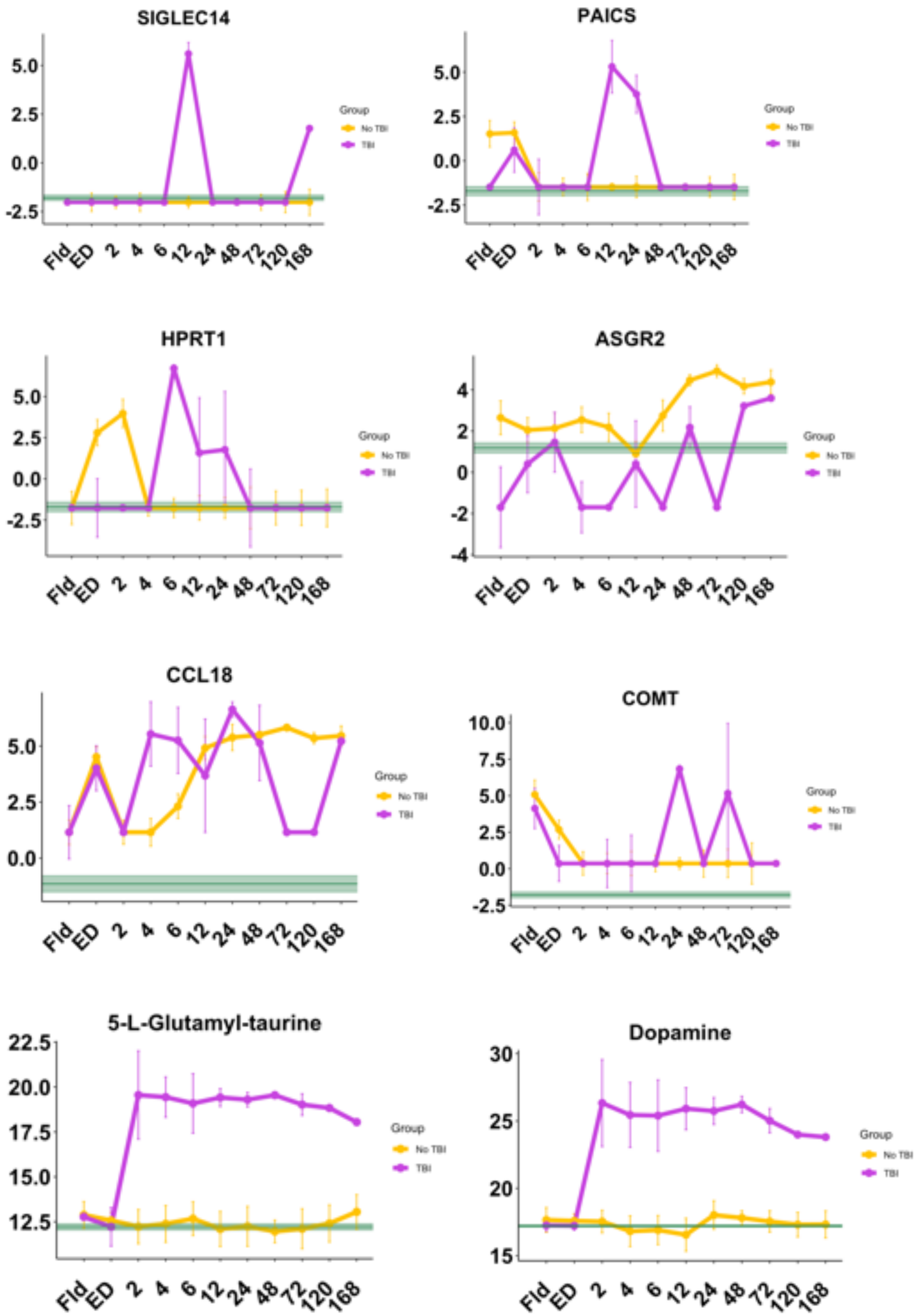

TBI in LSHT group

**Supplemental Figure 8 (Figure S8):** Timeseries expression patterns of significantly different proteins and metabolites of HT patients with TBI vs. no TBI.

*Supplementary FIGURE 9*

| Analyte | Overall VIP (ALI) |
|---|---|
| Creatine | 0.505493099 |
| CD163 | 0.480336142 |
| UBE2V1 | 0.476482583 |
| HBB | 0.415782482 |
| COL6A1 | 0.380514143 |
| UBB | 0.321015924 |
| PRDX1 | 0.275188976 |
| BLVRB | 0.27142977 |
| ACTN4 | 0.267683219 |
| Phosphocreatine | 0.263950285 |
| HBA1 | 0.260852234 |
| PRDX2 | 0.258616066 |
| LGlutamate | 0.236241193 |
| CA1 | 0.230647989 |
| snGlycerol.3phosphate | 0.211491355 |
| ATIC | 0.209205333 |
| HBD | 0.203422025 |
| S100A4 | 0.200997647 |

| Analyte | Overall VIP (ALI) |
| --- | --- |
| SERPINB1 | 0.195584265 |
| Methylglutarylcarnitine | 0.189615388 |

| Analyte | Overall VIP (Death) |
| --- | --- |
| Succinate | 1.39253468 |
| Malate | 0.848976109 |
| Spermidine | 0.587652959 |
| Putrescine | 0.495136648 |
| PRDX1 | 0.473797769 |
| Pantothenate | 0.417058577 |
| Spermine | 0.391357226 |
| Nonanoic.acid | 0.32669947 |
| PRDX2 | 0.305812086 |
| Diphosphate | 0.255147725 |
| NQO2 | 0.240083372 |
| Butyrylcarnitine | 0.214433257 |
| lovastatin | 0.198968385 |
| Hexenoylcarnitine | 0.195819693 |

| Analyte | Overall VIP (Death) |
| --- | --- |

| | |
|---|---|
| Nicotinamide | 0.19502469 |
| isobutyrylcarnitine | 0.183483378 |
| Isovalerylcarnitine | 0.163432513 |
| Alphalinolenyl.carnitine | 0.155009569 |
| Fumarate | 0.154890767 |
| GNAI2 | 0.151096442 |

| **Analyte** | **Overall VIP (ICU-free days)** |
|---|---|
| NME1 | 0.404136307 |
| MDH1 | 0.390024599 |
| Succinate | 0.311230923 |
| HBB | 0.309873989 |
| SOD1 | 0.301271699 |
| H2AC4 | 0.289427281 |
| TUBB | 0.288100299 |
| LDHA | 0.287323312 |
| HBA1 | 0.286353204 |
| FTL | 0.268498602 |

| **Analyte** | **Overall VIP (ICU-free days)** |
|---|---|
| PPIA | 0.259004567 |

| | |
|---|---|
| Spermine | 0.257693125 |
| SULT2A1 | 0.249505546 |
| DDT | 0.248729271 |
| UBB | 0.24192026 |
| LGALS1 | 0.234164738 |
| GAPDH | 0.231692085 |
| CMBL | 0.230495434 |
| PEBP1 | 0.229947602 |
| snGlycerol.3phosphate | 0.22670003 |

| **Analyte** | **Overall VIP (INR)** |
|---|---|
| Spermine | 0.94585515 |
| PRDX2 | 0.555240984 |
| GAPDH | 0.548134631 |
| RAN | 0.497226622 |
| Creatine | 0.41319906 |
| CDH5 | 0.39490895 |
| **Analyte** | **Overall VIP (INR)** |
| TPI1 | 0.390444566 |
| GSTA1 | 0.387530769 |

| | |
|---|---|
| SOD1 | 0.367296055 |
| FAH | 0.349657858 |
| PCBD1 | 0.344593777 |
| LGlutamate | 0.339463048 |
| Nobiletin | 0.327357851 |
| TXN | 0.314283754 |
| Prostaglandin.E2 | 0.311452881 |
| ENO1 | 0.287519144 |
| PRDX6 | 0.276623788 |
| CFL1 | 0.261960788 |
| H2AC4 | 0.25668572 |
| FABP1 | 0.254215123 |

| **Analyte** | **Overall VIP (MTX)** |
|---|---|
| Fumarate | 0.664533157 |
| Succinate | 0.644947586 |

| **Analyte** | **Overall VIP (MTX)** |
|---|---|
| Hexenoylcarnitine | 0.46750399 |
| N1Acetylspermidine | 0.395688357 |
| Valerylcarnitine | 0.357246569 |

| | |
|---|---|
| PRDX1 | 0.353981193 |
| COL18A1 | 0.331223143 |
| Lysine | 0.323779451 |
| Hypoxanthine | 0.28584183 |
| Isovalerylcarnitine | 0.276020309 |
| Xanthine | 0.274667022 |
| Malate | 0.220514147 |
| Putrescine | 0.213695743 |
| Spermine | 0.206130952 |
| Psychosine | 0.203155841 |
| KNG1.1 | 0.177600422 |
| SLC4A1 | 0.173636077 |
| Hexadecadienoylcarnitine | 0.160691983 |
| ECH1 | 0.157509287 |
| Propionylcarnitine | 0.151390003 |

| **Analyte** | **Overall VIP (Vent-free days)** |
|---|---|
| snGlycerol.3phosphate | 1.283569666 |
| Spermidine | 0.953882031 |
| TPI1 | 0.803891534 |

| | |
|---|---|
| PRDX6 | 0.729399322 |
| GSTA1 | 0.719629039 |
| MDH1 | 0.667600071 |
| Spermine | 0.656114552 |
| FABP1 | 0.632446349 |
| GAPDH | 0.605978663 |
| Succinate | 0.601837075 |
| Dehydroascorbate | 0.577622562 |
| FBP2 | 0.485000125 |
| SULT2A1 | 0.414279897 |
| Malate | 0.411999451 |
| ABHD14B | 0.398517225 |
| SOD1 | 0.337700763 |
| TUBB | 0.329721094 |
| **Analyte** | **Overall VIP (Vent-free days)** |
| ADH1B | 0.30585915 |
| TXN | 0.300628801 |
| CAP1 | 0.278949603 |

**VIP scores from RandomForest prediction of clinical outcomes.**

| Model (Death) | Risk | Coef |
|---|---|---|
| SL.ranger_All | 0.09063742 | 0.1894109 |
| SL.kernelKnn_All | 0.08177215 | 0.8105891 |

| Model (ALI) | Risk | Coef |
|---|---|---|
| SL.ranger_All | 0.2301309 | 0 |
| SL.kernelKnn_All | 0.231519 | 0.1779251 |
| SL.ksvm_All | 0.2243849 | 0.8220749 |

| Model (ICU-free days) | Risk | Coef |
|---|---|---|
| SL.xgboost_All | 0.2516347 | 0.00876711 |
| SL.ksvm_All | 0.1680393 | 0.99123289 |

| Model (Vent-free days) | Risk | Coef |
|---|---|---|
| SL.glmnet_All | 0.1241692 | 0.1059206 |

| | | |
|---|---|---|
| SL.ksvm_All | 0.1081759 | 0.8940794 |

| **Model (MTX)** | **Risk** | **Coef** |
|---|---|---|
| SL.ksvm_All | 0.07192628 | 1 |

| **Model (INR)** | **Risk** | **Coef** |
|---|---|---|
| SL.ranger_All | 0.2067699 | 0.274331 |
| SL.ksvm_All | 0.2021235 | 0.4941587 |
| SL.kernelKnn_All | 0.2102532 | 0.2315104 |

**Supplemental Figure 9 (Figure S9):** VIP analyte scores from RandomForest in predicting clinical outcomes, above. Below, ensemble model composition trained on a training set (75% split) for predicting outcomes of Death, Acute Lung Injury (ALI), Ventilator-free days, ICU-free days, coagulopathy by International Normalized Ratio (INR), and the need for massive transfusion (MTX) in a test set (25% split). From SuperLearner documentation: “Risk is a measure of model accuracy or performance. We want our models to minimize the estimated risk, which means the model is making the fewest mistakes in its prediction. It's basically the mean-squared error in a regression model, but you can customize it if you want.” Also: “The coefficient column tells us the weight or importance of each individual learner in the overall ensemble. By default the weights are always greater than or equal to 0 and sum to 1. In this case we only have one algorithm so the coefficient has to be 1. If a coefficient is 0 it means that the algorithm isn't being used in the SuperLearner ensemble.” For methods used: ranger = RandomForest; kernelKnn = K-Nearest Neighbors; ksvm = Support Vector Machine; xgboost = XGBoost; glmnet = Generalized Linear Model.

#### Supplementary FIGURE 10

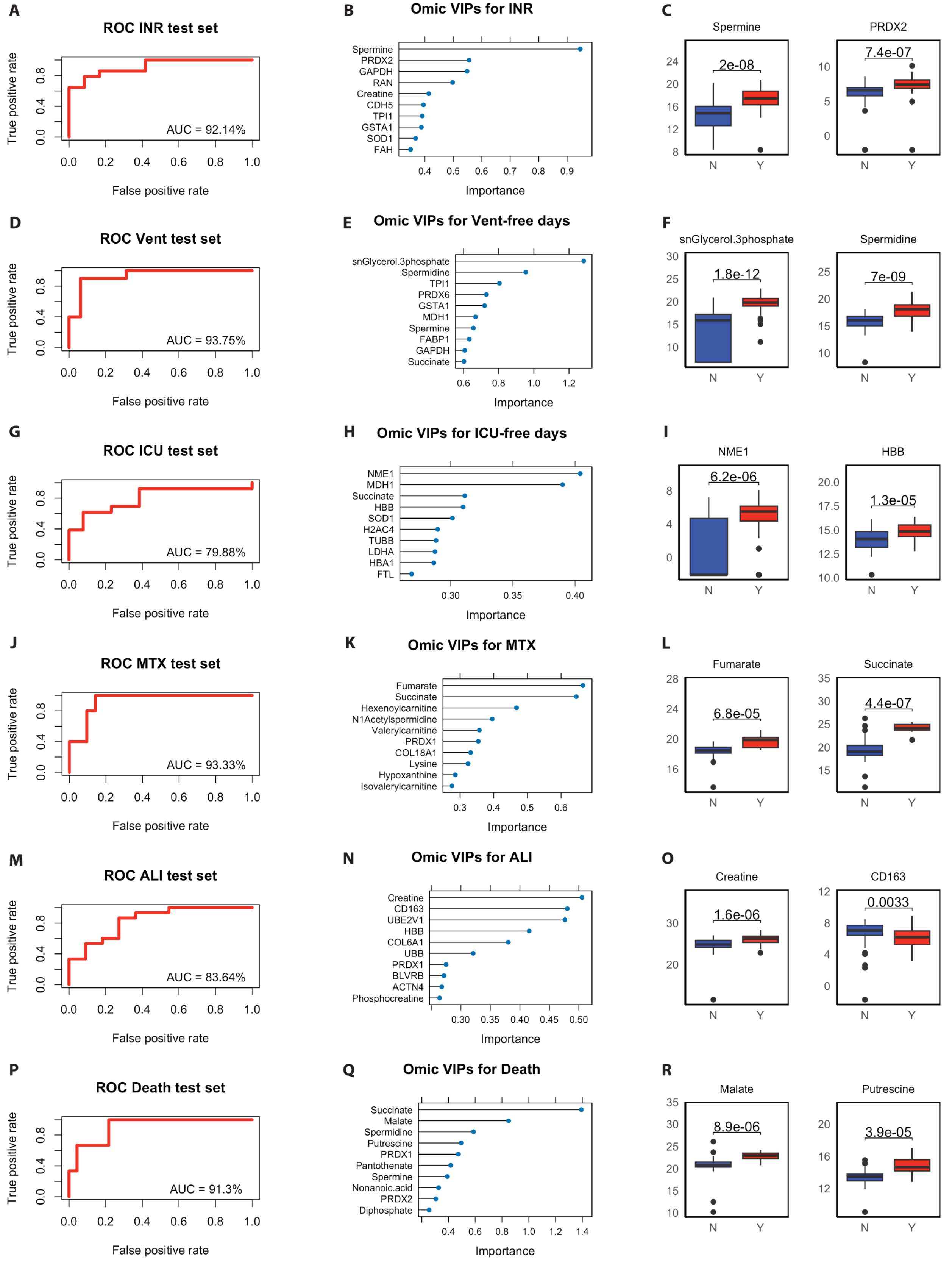

**Supplemental Figure 10 (Figure S10) Ensemble learning to predict patient outcomes:** ROC-AUC, VIP analysis, and top VIPs of COMBAT omics data training set predicting outcomes in COMBAT omics test set. Ensemble models were trained and tested on separate sets of ED data to predict patient outcomes, and identify important analytes (VIP analysis from RandomForest) contributing to outcomes. ROC analysis of Ensemble model performance with AUC (1st column), VIP analysis (2nd column), and boxplots of top VIP analytes (3rd column, t-test) identified to predict (**A-C**) coagulopathy by INR, (**D-F**) Ventilator-free days, (**G-I**) ICU-free days, (**J-K**) the need for massive transfusion (MTX), (**M-O**) acute lung injury (ALI), and (**P-R**) death.

*Supplementary FIGURE 11*

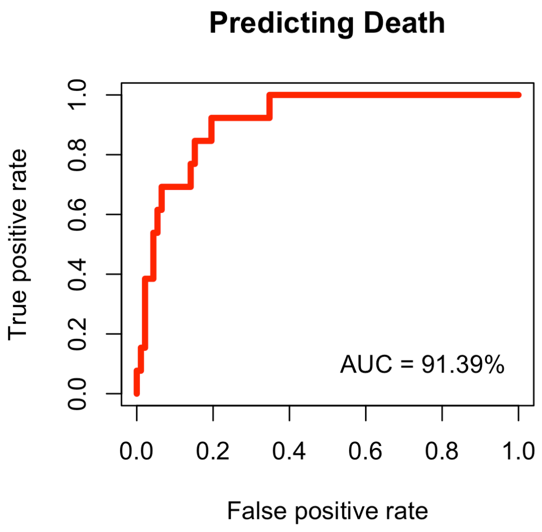

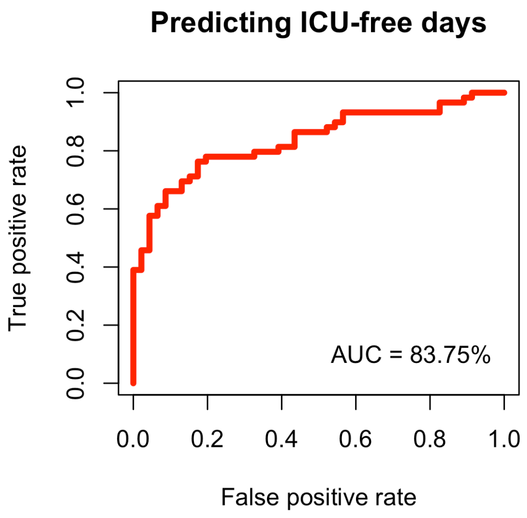

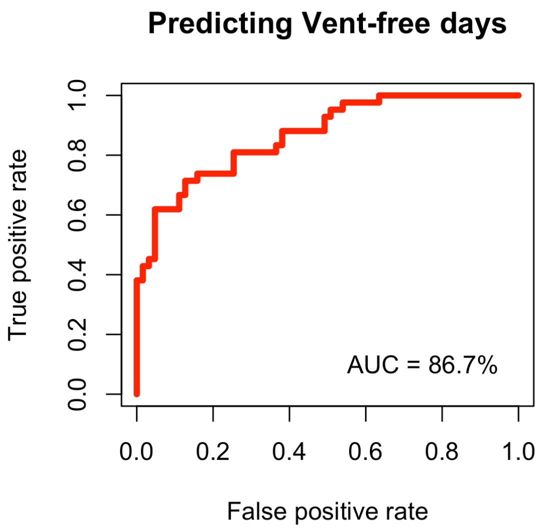

| Model (Death) | Risk | Coef |
|---|---|---|
| SL.ranger_All | 0.08202811 | 0.47334635 |
| SL.kernelKnn_All | 0.09030822 | 0.06722673 |
| SL.xgboost_All | 0.08239314 | 0.45942692 |

| Model (Vent-free days) | Risk | Coef |
|---|---|---|
| SL.ranger_All | 0.2076651 | 0.44311812 |
| SL.xgboost_All | 0.246541 | 0.06985625 |
| SL.biglasso_All | 0.2078765 | 0.48702563 |

| Model (ICU-free days) | Risk | Coef |
|---|---|---|
| SL.ranger_All | 0.1850735 | 0.48827214 |
| SL.kernelKnn_All | 0.2191781 | 0.02471672 |
| SL.xgboost_All | 0.1855299 | 0.48701114 |

**Supplemental Figure 11 (Figure S11):** ROC-AUC, and ensemble model composition trained on an independent training dataset set of trauma patients for predicting outcomes of Death, ICU-free days, and Ventilator-free days in the COMBAT test set.

*Supplementary FIGURE 12*

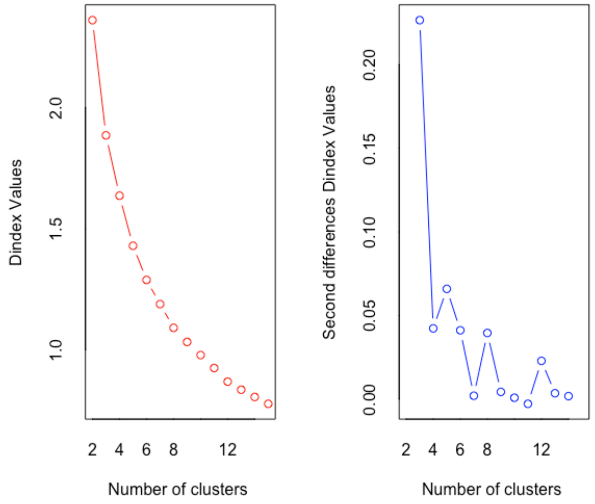

Ward method to minimize the total within-cluster variance. From NbClust():

*** : The Hubert index is a graphical method of determining the number of clusters.

In the plot of Hubert index, we seek a significant knee that corresponds to a significant increase of the value of the measure i.e the significant peak in Hubert index second differences plot.

*** : The D index is a graphical method of determining the number of clusters.

In the plot of D index, we seek a significant knee (the significant peak in Dindex second differences plot) that corresponds to a significant increase of the value of the measure.

*******************************************************************

* Among all indices:

* 10 proposed 2 as the best number of clusters

* 8 proposed 3 as the best number of clusters

* 1 proposed 4 as the best number of clusters

* 1 proposed 6 as the best number of clusters

* 3 proposed 8 as the best number of clusters

* 1 proposed 12 as the best number of clusters

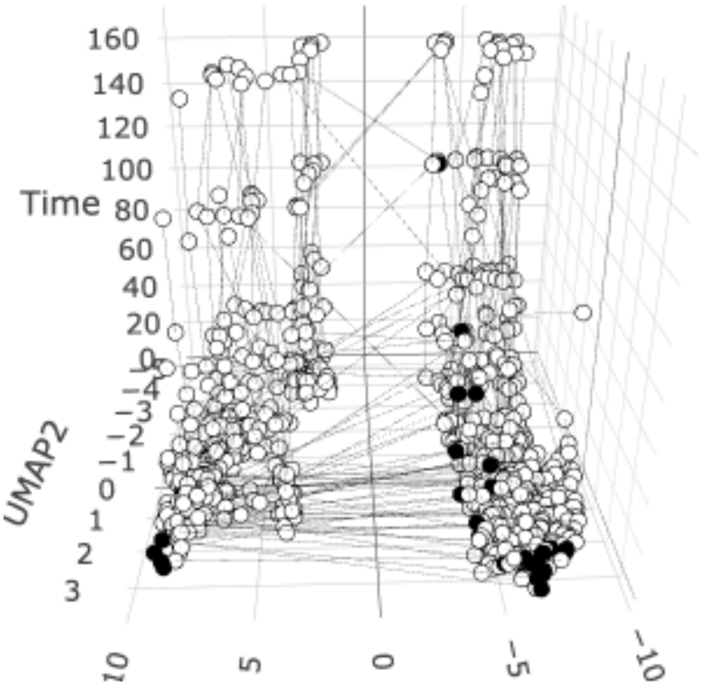

3D UMAP of metabolic patient-timepoints colored by association with death

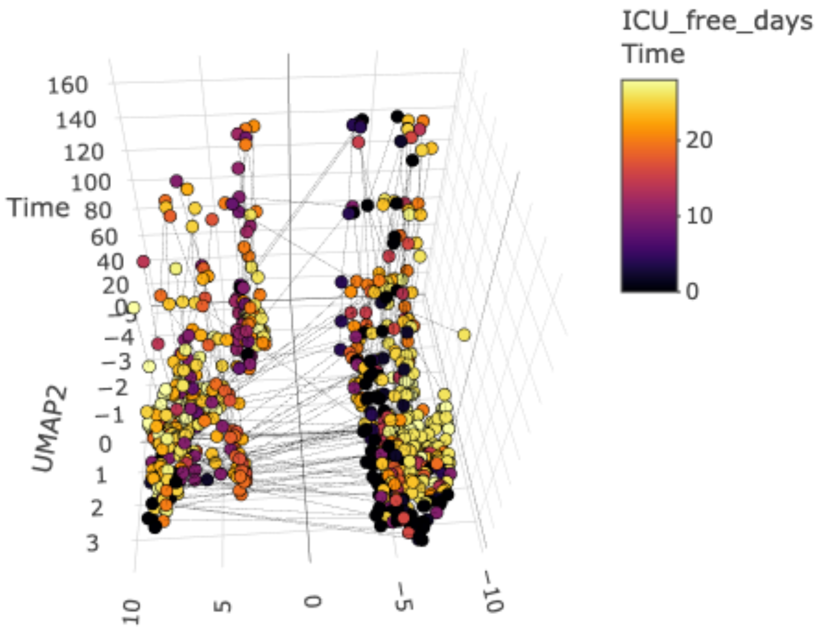

3D UMAP of metabolic patient-timepoints colored by association with ICU-free days (fewer = more severely injured)

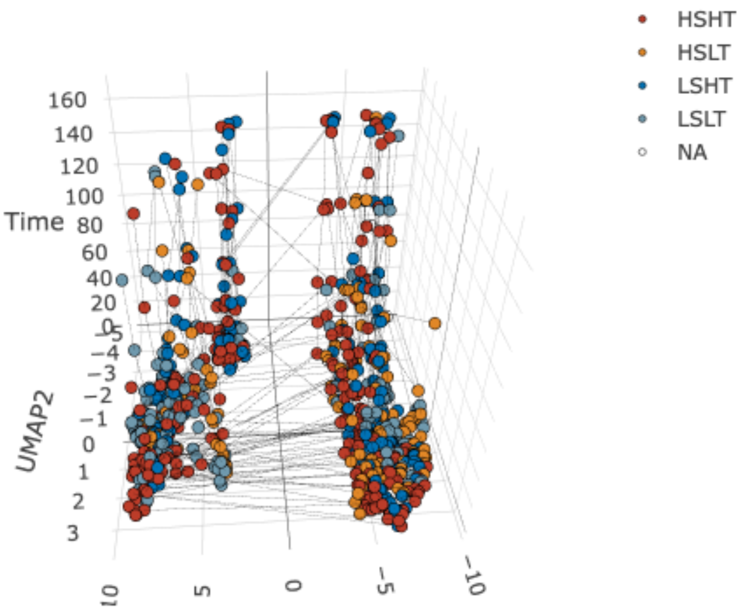

3D UMAP of metabolic patient-timepoints colored by association with shock and tissue injury

**Supplemental Figure 12 (Figure S12):** Metabolomic patient state determination of cluster number, and basic characterization and distribution metabolic patient-timepoints associated with more severe- and less severe- injury.

#### *Supplementary FIGURE 13*

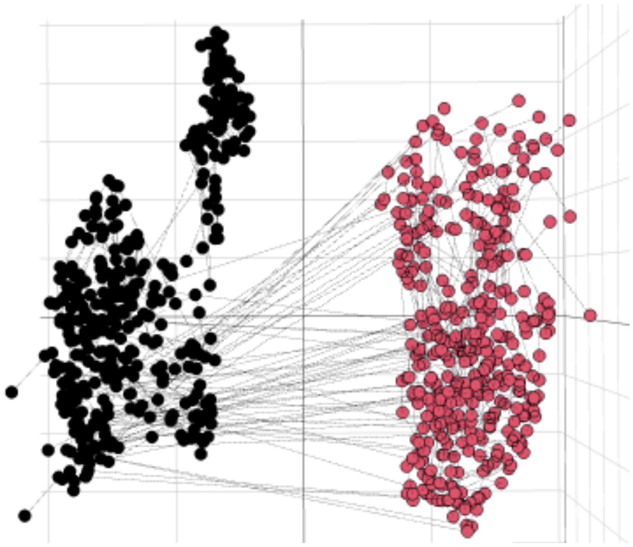

**Left** **Right**

(Biosynthesis) (Catabolism)

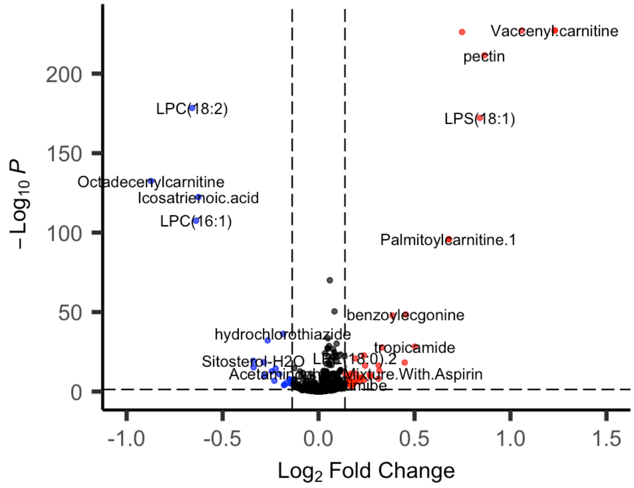

**Metabolites significantly elevated in Biosynthesis**

| Metabolite | fdrpval | logFC |
|---|---|---|
| LPC(18:2) | 3.79E-179 | -0.6585397 |
| Octadecenylcarnitine | 3.45E-133 | -0.8728294 |
| Icosatrienoic.acid | 4.25E-123 | -0.6262291 |
| LPC(16:1) | 3.18E-108 | -0.6382225 |
| hydrochlorothiazide | 4.67E-37 | -0.1846542 |
| ADP | 6.26E-33 | -0.2654047 |
| Sitosterol-H2O | 4.74E-20 | -0.339887 |
| lisinopril | 3.58E-19 | -0.2856864 |
| L-Cysteine | 5.03E-16 | -0.3382385 |
| LPS(18:0) | 2.01E-15 | -0.1268777 |
| AMP | 3.67E-15 | -0.2238388 |
| Eremosulphoxinolide.B | 2.09E-13 | -0.2440441 |
| Hydroxyundecanoyl.carnitine | 2.15E-13 | -0.0965483 |
| Cervonyl.carnitine | 2.00E-12 | -0.08117 |
| S-allyl-L-cysteine | 7.71E-12 | -0.0404638 |
| Guanine | 1.07E-11 | -0.2041742 |
| Trimeprazine | 1.12E-10 | -0.2832774 |
| Umbelliferone | 4.28E-10 | -0.0905936 |
| L-Asparagine | 1.43E-09 | -0.0292867 |
| 4-Methylamphetamine | 1.64E-09 | -0.0676538 |

**Metabolites significantly elevated in Catabolism**

| Metabolite | fdrpval | logFC |
|---|---|---|
| Vaccenyl.carnitine | 0.00E+00 | 1.23094378 |
| Stearoylcarnitine | 0.00E+00 | 1.05819157 |
| Oleoylcarnitine | 0.00E+00 | 1.23094378 |
| Heptadecanoyl.carnitine | 6.65E-227 | 0.74752275 |
| pectin | 2.72E-212 | 0.86440342 |
| LPS(18:1) | 6.30E-173 | 0.84052097 |
| Palmitoylcarnitine.1 | 1.50E-96 | 0.67886337 |
| Palmitoylcarnitine.2 | 1.50E-96 | 0.67886337 |
| Diallyl.disulfide | 9.73E-71 | 0.05875937 |
| xi-8-Acetonyldihydrosanguinarine | 3.73E-51 | 0.08298324 |
| benzoylecgonine | 3.58E-49 | 0.4544165 |
| Huperzine.B | 1.62E-48 | 0.38575068 |
| Indole | 2.80E-34 | 0.04887106 |
| LPC(18:1) | 7.04E-31 | 0.09387344 |
| Indole-3-acetaldehyde | 2.13E-29 | 0.05465967 |

| | | |
|---|---|---|
| tropicamide | 5.38E-29 | 0.50176504 |
| Psychosine | 1.95E-28 | 0.33074453 |
| 3-Methyleneoxindole | 4.44E-28 | 0.04677066 |
| 2-Hydroxyibuprofen | 1.22E-27 | 0.04841038 |
| LPC(16:0) | 1.91E-27 | 0.07578637 |

**Supplemental Figure 13 (Figure S13):** Identification of two patient “meta” metabolomic states, with metabolites involved in biosynthesis of macromolecules (L) and those involved in catabolism (R) enriched in different states.

### Supplementary FIGURE 14

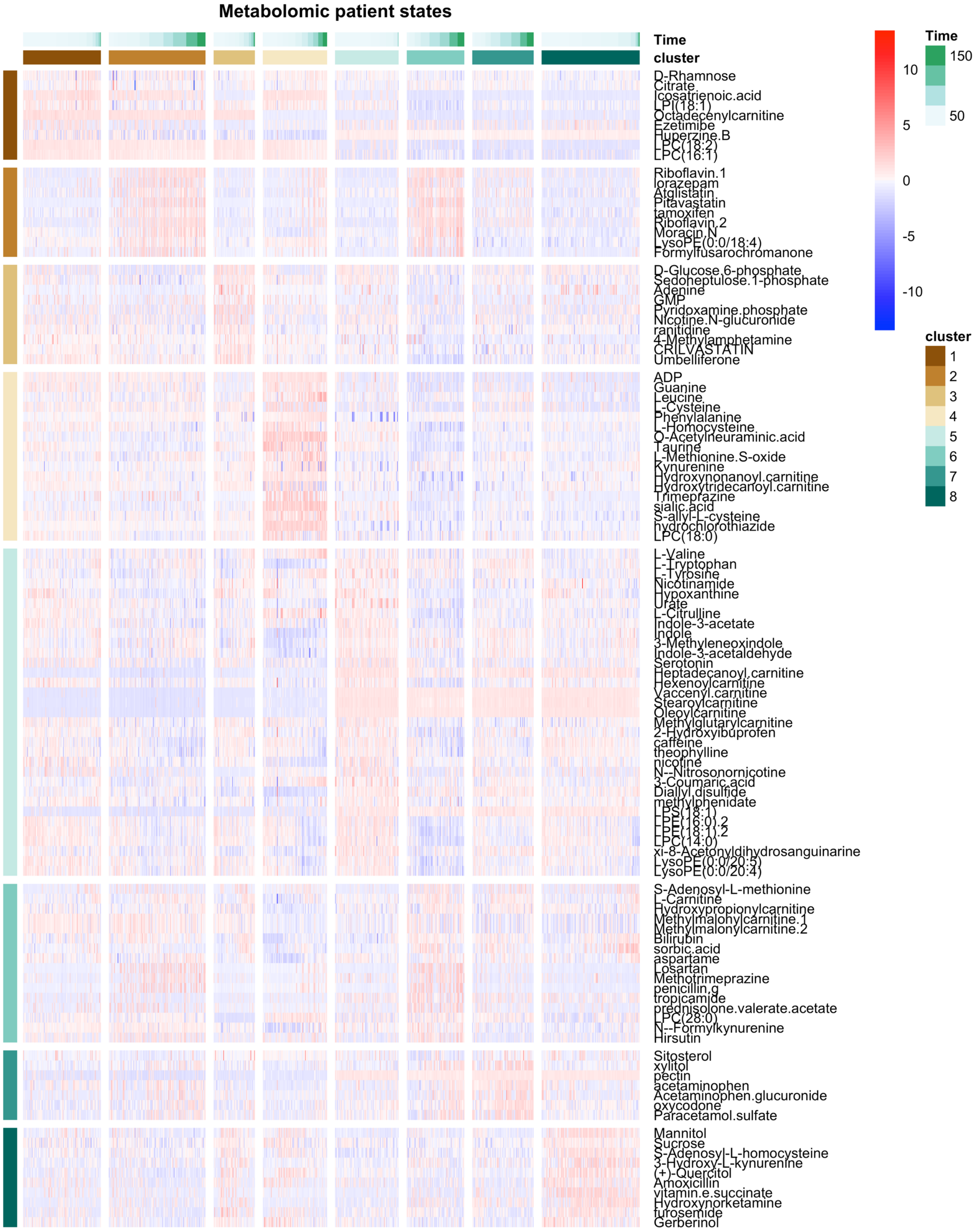

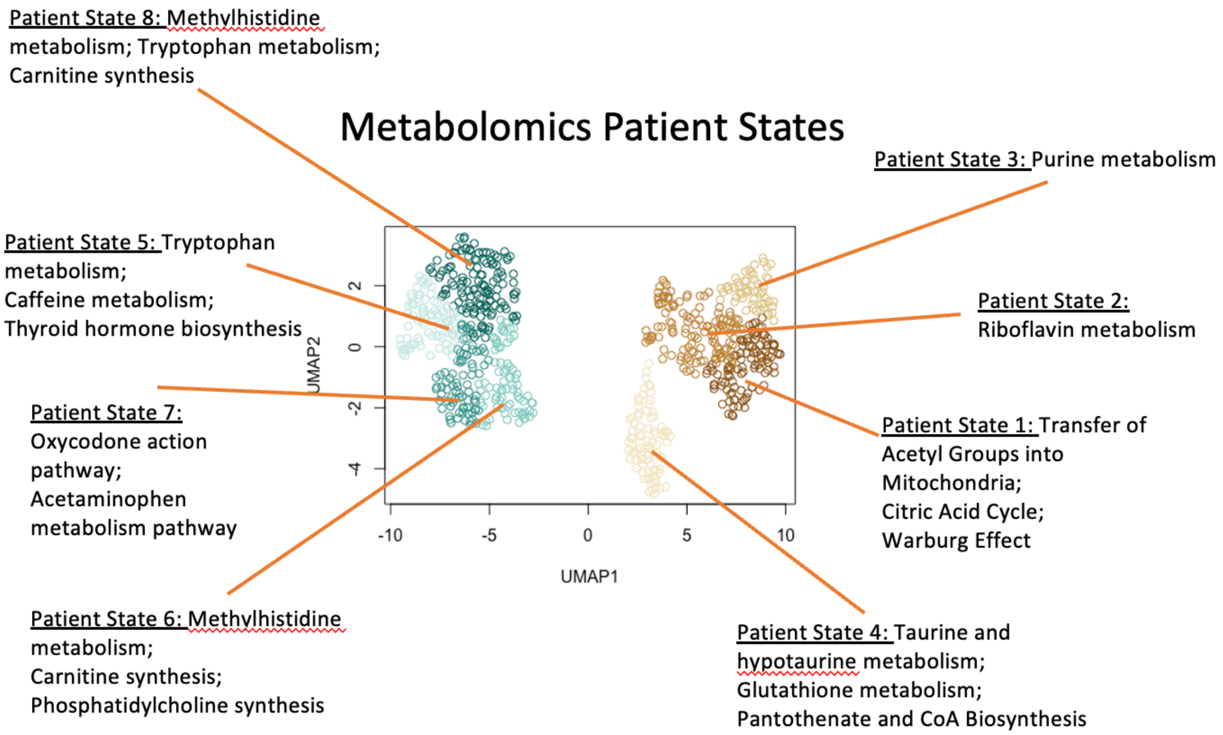

**Supplemental Figure 14 (Figure S14):** Heatmap of longitudinal patient state signature expression across all patient states. 2D UMAP of patient-timepoints colored by patient state, with overview of metabolic state as determined by pathway enrichment of analytes expressed at higher levels within each patient state.

#### Supplementary FIGURE 15

##### Vitals

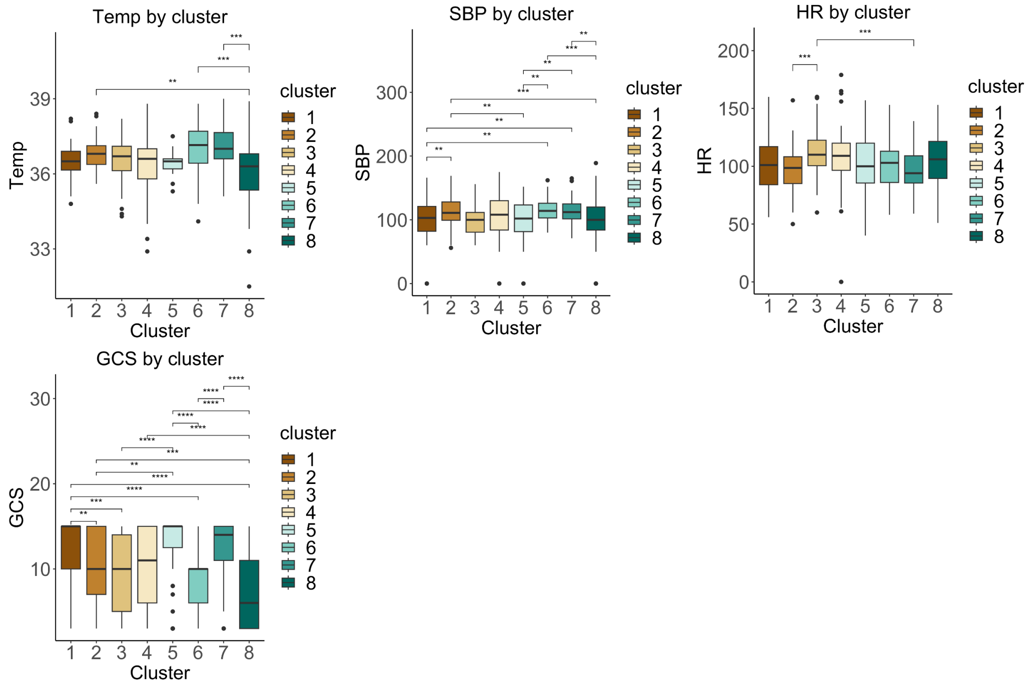

##### Blood chemistry

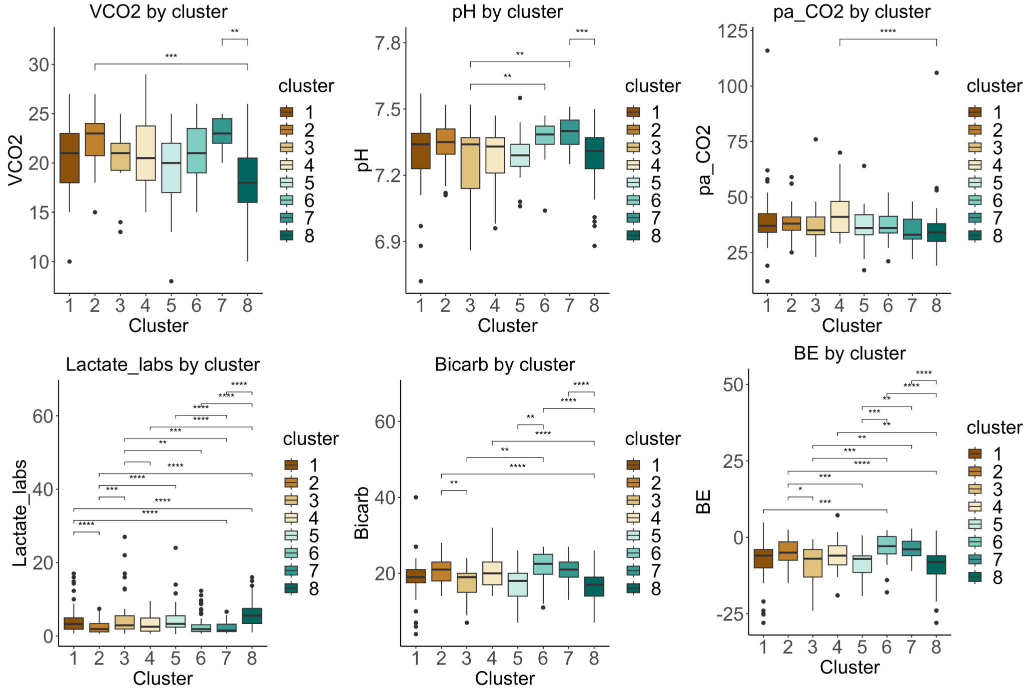

#### Blood products

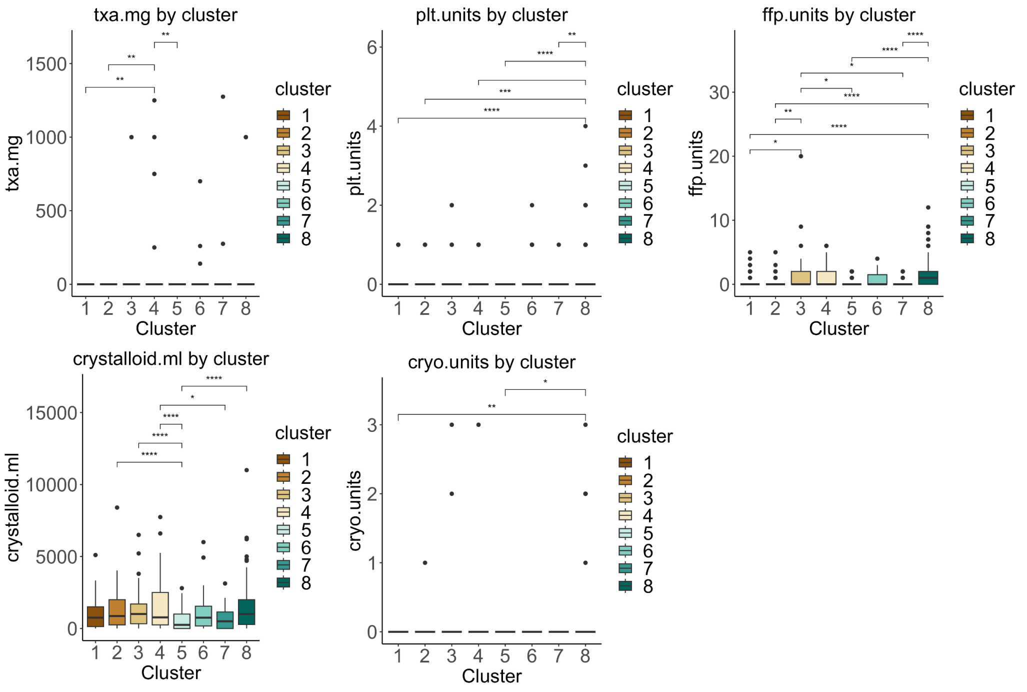

#### Blood count

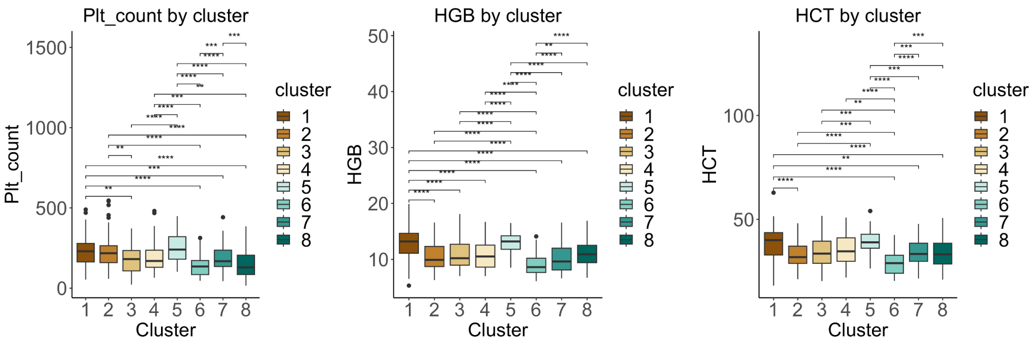

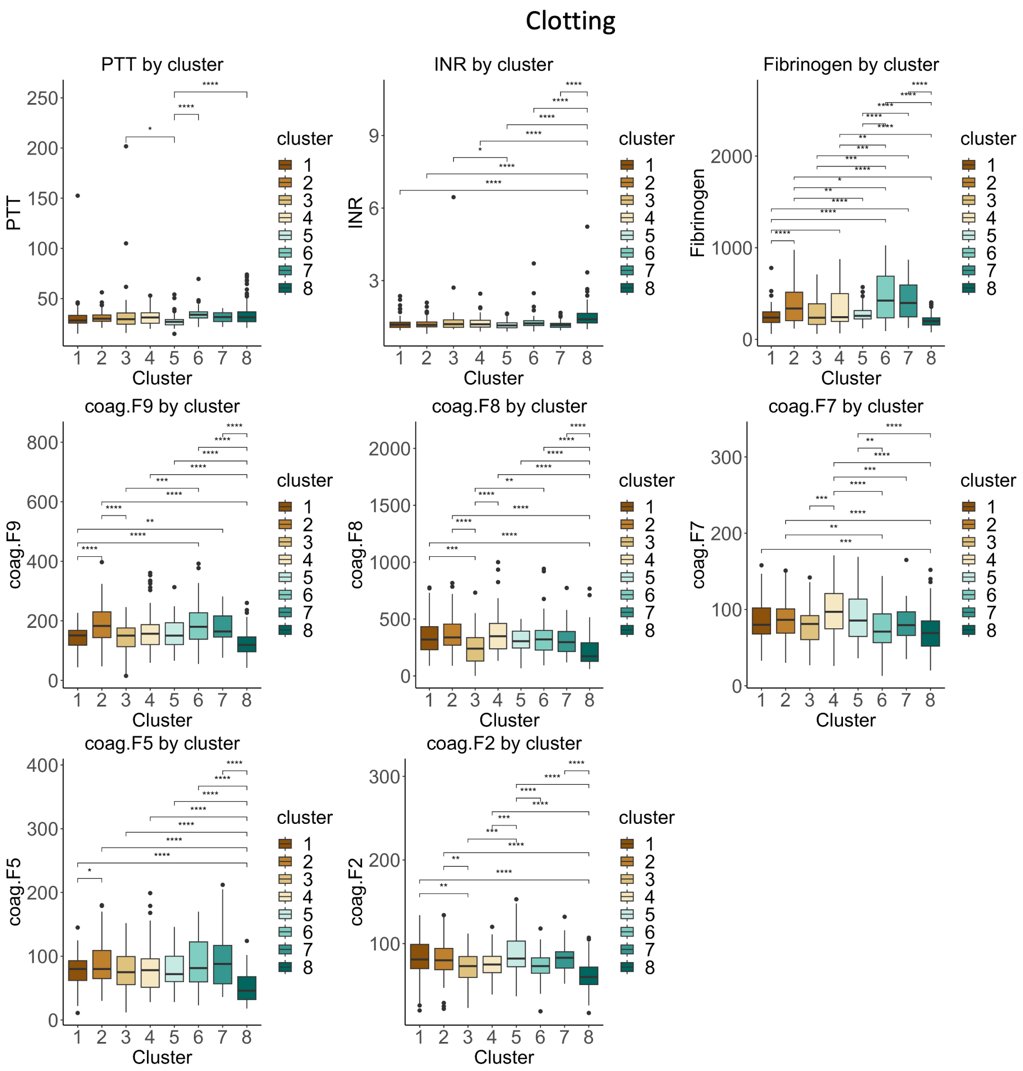
Clotting
PTT by cluster
INR by cluster
Fibrinogen by cluster
coag.F9 by cluster
coag.F8 by cluster
coag.F7 by cluster
coag.F5 by cluster
coag.F2 by cluster
Cluster
cluster

#### TEGs

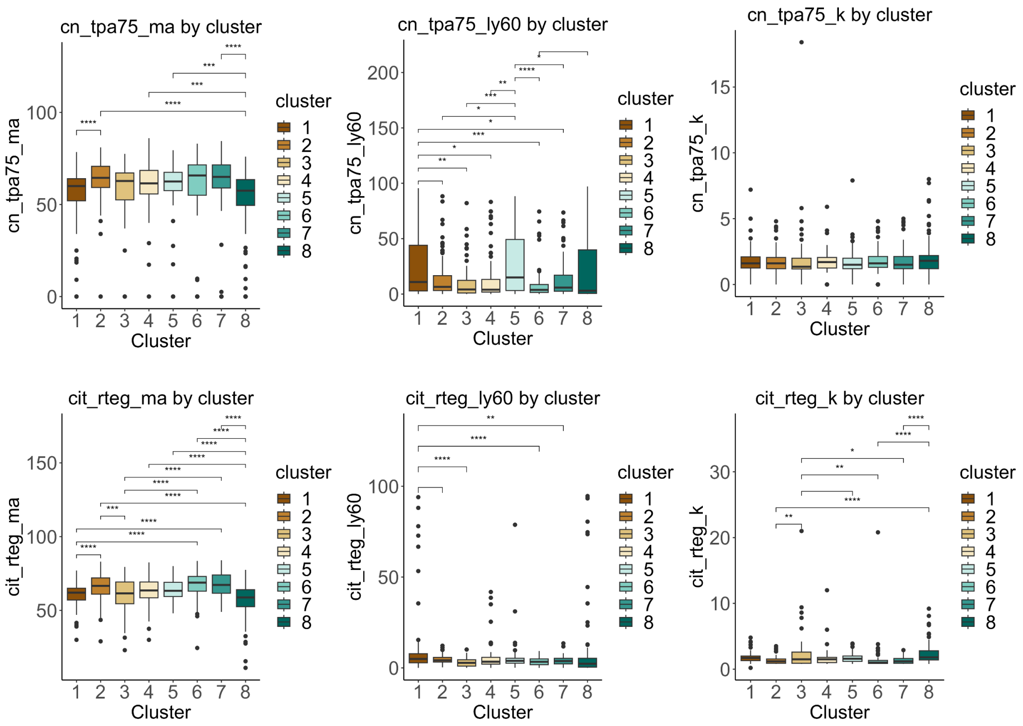
cn_tpa75_ma by cluster
cn_tpa75_ly60 by cluster
cn_tpa75_k by cluster
cit_rteg_ma by cluster
cit_rteg_ly60 by cluster
cit_rteg_k by cluster
cn_tpa75_ma
cn_tpa75_ly60
cn_tpa75_k
cit_rteg_ma
cit_rteg_ly60
cit_rteg_k
Cluster
cluster
1
2
3
4
5
6
7
8

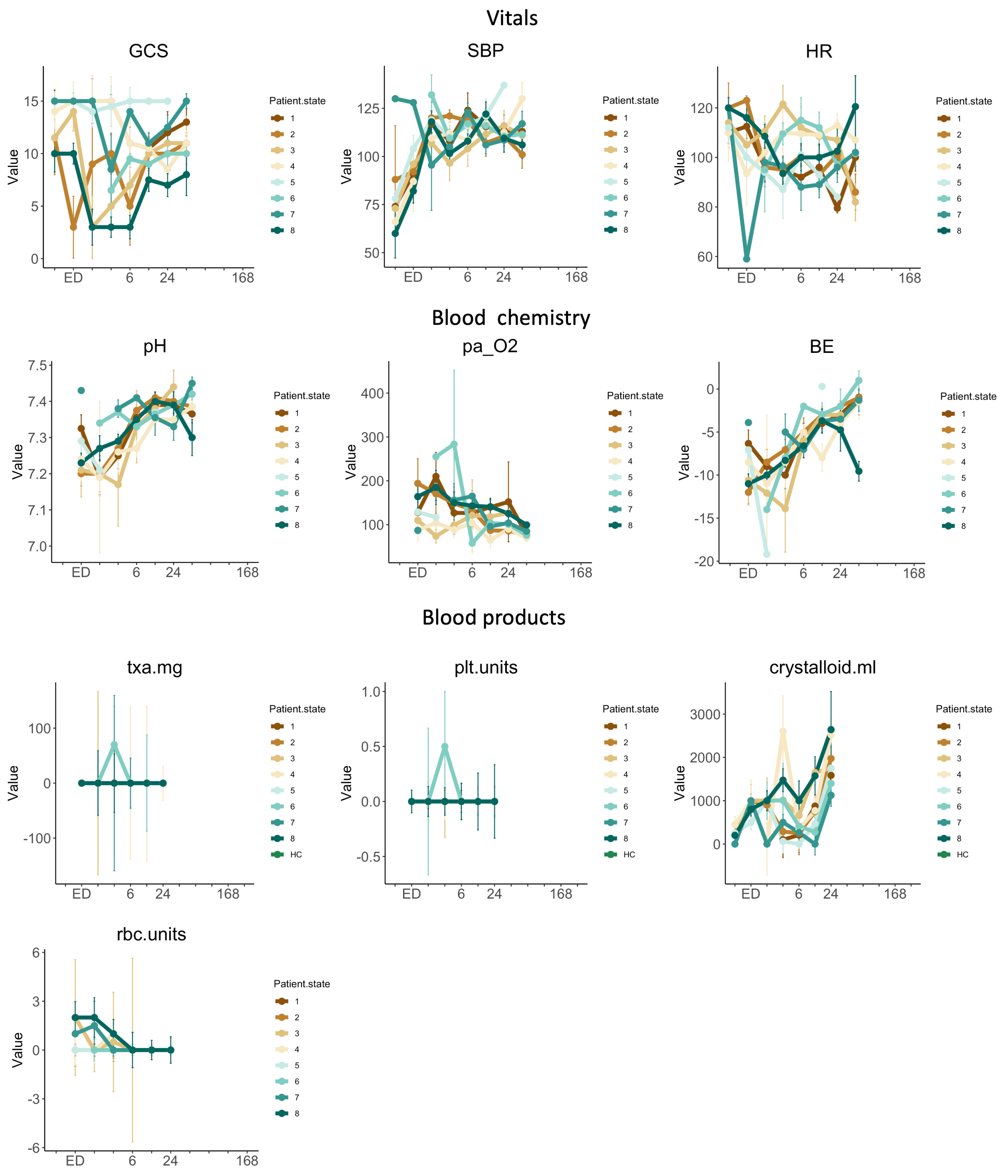
Vitals
GCS
SBP
HR
Blood chemistry
pH
pa_O2
BE
Blood products
txa.mg
plt.units
crystalloid.ml
rbc.units
Value
ED
6
24
168
Patient.state
1
2
3
4
5
6
7
8
HC

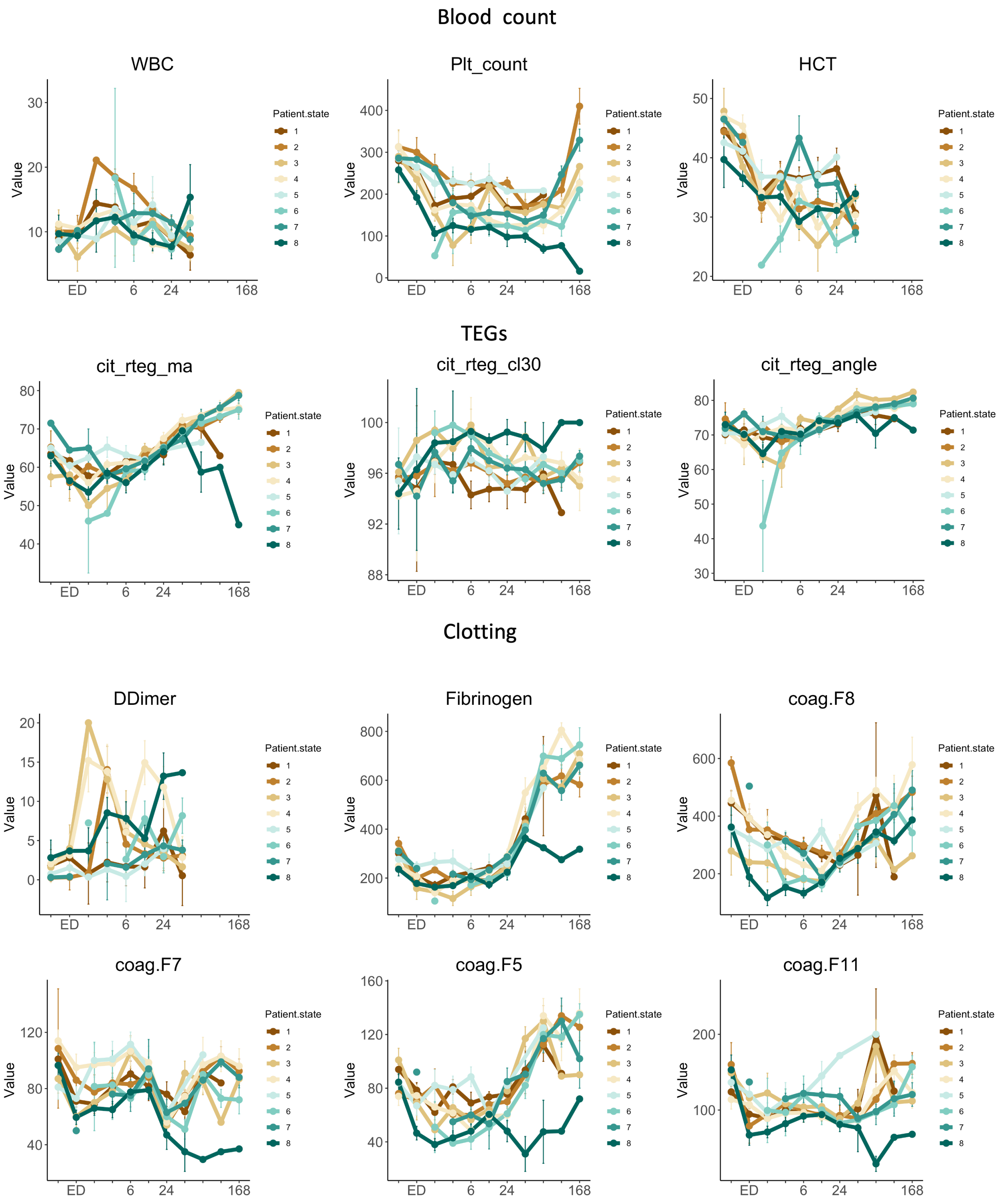
Blood count
WBC
Plt_count
HCT
TEGs
cit_rteg_ma
cit_rteg_cl30
cit_rteg_angle
Clotting
DDimer
Fibrinogen
coag.F8
coag.F7
coag.F5
coag.F11
Value
ED
6
24
168
Patient.state
1
2
3
4
5
6
7
8

| Clinical assay | obs.tot | obs.groups | df.between | df.within | statistic | pvalue |
|---|---|---|---|---|---|---|
| HGB | 672 | 8 | 7 | 249.869192 | 32.0299243 | 1.52E-31 |
| Fibrinogen | 623 | 8 | 7 | 225.797311 | 29.301126 | 1.40E-28 |
| Lactate_labs | 599 | 8 | 7 | 203.467424 | 21.0689236 | 3.19E-21 |
| Plt_count | 662 | 8 | 7 | 247.22832 | 19.6762279 | 7.08E-21 |
| coag.F5 | 637 | 8 | 7 | 237.403293 | 18.798225 | 6.96E-20 |
| cit_rteg_g | 732 | 8 | 7 | 278.043837 | 18.1806284 | 7.16E-20 |
| coag.F9 | 637 | 8 | 7 | 238.782345 | 18.1203512 | 2.90E-19 |
| cit_rteg_ma | 732 | 8 | 7 | 281.437955 | 15.571945 | 3.10E-17 |
| HCT | 495 | 8 | 7 | 174.913501 | 15.6259642 | 7.73E-16 |
| BD | 367 | 8 | 7 | 112.681753 | 17.0857449 | 3.06E-15 |
| GCS | 462 | 8 | 7 | 171.93738 | 14.6225072 | 7.05E-15 |
| coag.F2 | 637 | 8 | 7 | 242.048161 | 13.1189201 | 2.61E-14 |
| cit_rteg_k | 730 | 8 | 7 | 272.959453 | 11.580108 | 6.97E-13 |
| coag.F8 | 637 | 8 | 7 | 242.09736 | 11.6085949 | 1.03E-12 |
| cn_tpa150_ly30 | 705 | 8 | 7 | 272.661567 | 11.0116386 | 2.98E-12 |
| coag.plasm | 637 | 8 | 7 | 242.180652 | 11.1782633 | 2.99E-12 |
| cit_rteg_angle | 732 | 8 | 7 | 278.026487 | 10.4938502 | 1.06E-11 |
| crystalloid.ml | 561 | 8 | 7 | 180.876472 | 8.55799156 | 4.78E-09 |
| coag.F7 | 637 | 8 | 7 | 241.232997 | 8.1898913 | 5.96E-09 |
| INR | 650 | 8 | 7 | 242.693277 | 8.15902868 | 6.37E-09 |
| BE | 320 | 8 | 7 | 101.767401 | 9.15607727 | 9.74E-09 |
| Bicarb | 303 | 8 | 7 | 93.8725486 | 9.20273689 | 1.27E-08 |
| cn_ma | 730 | 8 | 7 | 280.097883 | 7.37208916 | 3.93E-08 |
| rbc.units | 492 | 8 | 7 | 172.95433 | 7.71901669 | 4.11E-08 |
| PTT | 645 | 8 | 7 | 237.134172 | 7.42145772 | 4.54E-08 |
| cn_tpa75_ly60 | 730 | 8 | 7 | 285.410339 | 7.25584342 | 5.22E-08 |
| cit_rteg_cl60 | 730 | 8 | 7 | 277.091131 | 6.91300711 | 1.37E-07 |
| SBP | 596 | 8 | 7 | 223.753583 | 6.70031148 | 3.27E-07 |
| ffp.units | 561 | 8 | 7 | 185.891975 | 6.81426761 | 3.31E-07 |
| plt.units | 492 | 8 | 7 | 148.008595 | 6.93377258 | 3.87E-07 |
| cit_rteg_ly60 | 730 | 8 | 7 | 282.733108 | 6.40891815 | 5.19E-07 |
| dly60 | 730 | 8 | 7 | 286.741177 | 6.26147598 | 7.62E-07 |
| cn_tpa75_r | 730 | 8 | 7 | 284.021274 | 6.26417345 | 7.64E-07 |
| cn_tpa75_ma | 730 | 8 | 7 | 278.941274 | 6.16724945 | 1.01E-06 |
| cn_tpa75_ly30 | 703 | 8 | 7 | 269.018501 | 5.93050903 | 1.99E-06 |
| DDimer | 417 | 8 | 7 | 146.361415 | 6.15212814 | 2.57E-06 |
| cit_rteg_cl30 | 732 | 8 | 7 | 286.120234 | 5.80888293 | 2.60E-06 |
| cn_r | 730 | 8 | 7 | 284.205473 | 5.79008563 | 2.75E-06 |
| dly30 | 730 | 8 | 7 | 283.536401 | 5.418062 | 7.55E-06 |

| Clinical assay | obs.tot | obs.groups | df.between | df.within | statistic | pvalue |
|---|---|---|---|---|---|---|
| pH | 303 | 8 | 7 | 95.9687864 | 5.57294251 | 2.08E-05 |
| pa_O2 | 303 | 8 | 7 | 100.267526 | 4.95330589 | 7.69E-05 |
| Potassium | 213 | 8 | 7 | 52.5624021 | 5.11380688 | 0.00017649 |
| cit_rteg_ly30 | 732 | 8 | 7 | 281.810249 | 4.18231839 | 0.00021111 |
| VCO2 | 171 | 8 | 7 | 45.2548878 | 4.78735603 | 0.00043084 |
| coag.F11 | 636 | 8 | 7 | 239.943212 | 3.78078305 | 0.000654 |
| Temp | 299 | 8 | 7 | 103.735413 | 3.92151175 | 0.00078221 |
| Creatinine_labs | 162 | 8 | 7 | 44.9295733 | 4.31524808 | 0.00100345 |
| DBP | 563 | 8 | 7 | 210.463873 | 3.34780305 | 0.00209867 |
| HR | 601 | 8 | 7 | 220.277179 | 3.19387961 | 0.00305657 |
| rteg_ma | 118 | 7 | 6 | 25.0586145 | 4.22317554 | 0.00453107 |
| cn_angle | 730 | 8 | 7 | 279.44464 | 2.88185501 | 0.00637527 |
| WBC | 444 | 8 | 7 | 160.23503 | 2.82056331 | 0.00848894 |
| ion_Calcium | 295 | 8 | 7 | 76.2161037 | 2.9200336 | 0.00923469 |
| cn_tpa75_angle | 730 | 8 | 7 | 278.412881 | 2.67940653 | 0.01063581 |
| pa_CO2 | 303 | 8 | 7 | 95.7843568 | 2.48750586 | 0.02164424 |
| rteg_g | 118 | 7 | 6 | 25.192887 | 2.91246698 | 0.02690931 |
| cn_ly30 | 730 | 8 | 7 | 283.174119 | 2.22806125 | 0.03216521 |
| rteg_angle | 118 | 7 | 6 | 26.1213869 | 2.76878465 | 0.03220545 |
| rteg_ly30 | 116 | 7 | 6 | 26.5167044 | 2.6335622 | 0.0389193 |
| cn_k | 730 | 8 | 7 | 280.176091 | 2.03525686 | 0.05083452 |
| cryo.units | 492 | 8 | 7 | 307.590718 | 2.00843396 | 0.05376067 |
| txa.mg | 492 | 8 | 7 | 182.134915 | 2.0217336 | 0.05458295 |
| rteg_k | 118 | 7 | 6 | 27.3458195 | 1.87413166 | 0.12162678 |
| Glucose | 137 | 8 | 7 | 32.5504206 | 1.78305899 | 0.12456626 |
| cit_rteg_r | 730 | 8 | 7 | 283.139595 | 1.36122403 | 0.22167926 |
| rteg_act | 118 | 7 | 6 | 24.5619012 | 1.34788065 | 0.2744042 |
| cit_rteg_sp | 732 | 8 | 7 | 290.814853 | 1.18428625 | 0.31157353 |
| rteg_sp | 118 | 7 | 6 | 24.0340256 | 1.20261248 | 0.33877534 |
| Calcium | 170 | 8 | 7 | 44.6394656 | 1.1241626 | 0.36525973 |
| rteg_r | 118 | 7 | 6 | 24.8373631 | 1.1410685 | 0.36836764 |
| cit_rteg_act | 732 | 8 | 7 | 285.926203 | 1.06171705 | 0.38836222 |
| cn_tpa75_k | 730 | 8 | 7 | 278.312693 | 0.37823469 | 0.91467911 |

**Supplemental Figure 15 (Figure S15):** Boxplots and line charts showing trends and statistical comparisons (ANOVA TukeyHSD posthoc) of clinical measurements per metabolomics patient state.

*Supplementary FIGURE 16*

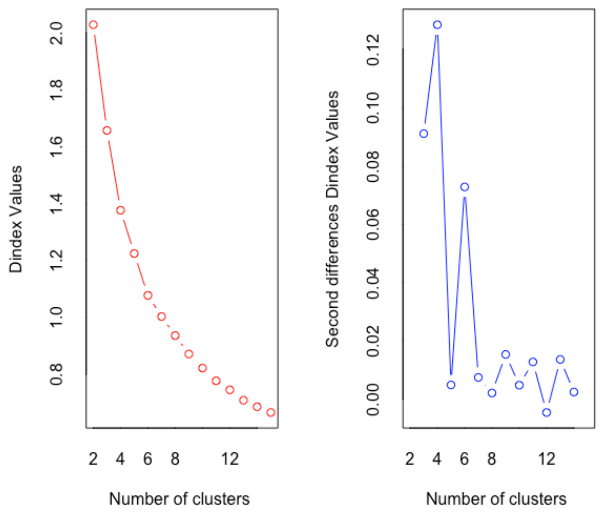

Ward method to minimize the total within-cluster variance. From NbClust():

*** : The Hubert index is a graphical method of determining the number of clusters.

In the plot of Hubert index, we seek a significant knee that corresponds to a significant increase of the value of the measure i.e the significant peak in Hubert index second differences plot.

*** : The D index is a graphical method of determining the number of clusters.

In the plot of D index, we seek a significant knee (the significant peak in Dindex second differences plot) that corresponds to a significant increase of the value of the measure.

*******************************************************************

* Among all indices measured:

* 8 proposed 2 as the best number of clusters

* 4 proposed 3 as the best number of clusters

* 4 proposed 4 as the best number of clusters

* 1 proposed 5 as the best number of clusters

* 4 proposed 6 as the best number of clusters

* 3 proposed 13 as the best number of clusters

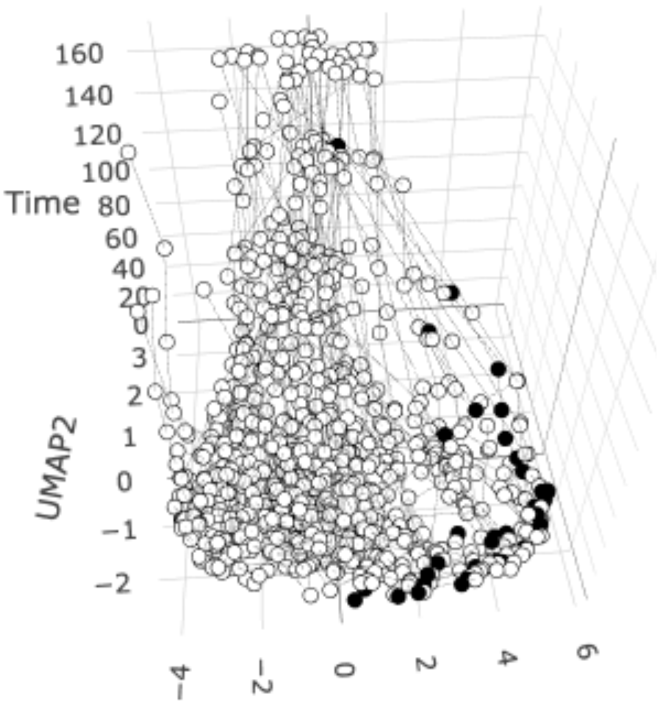

3D UMAP of proteomic patient-timepoints colored by association with death

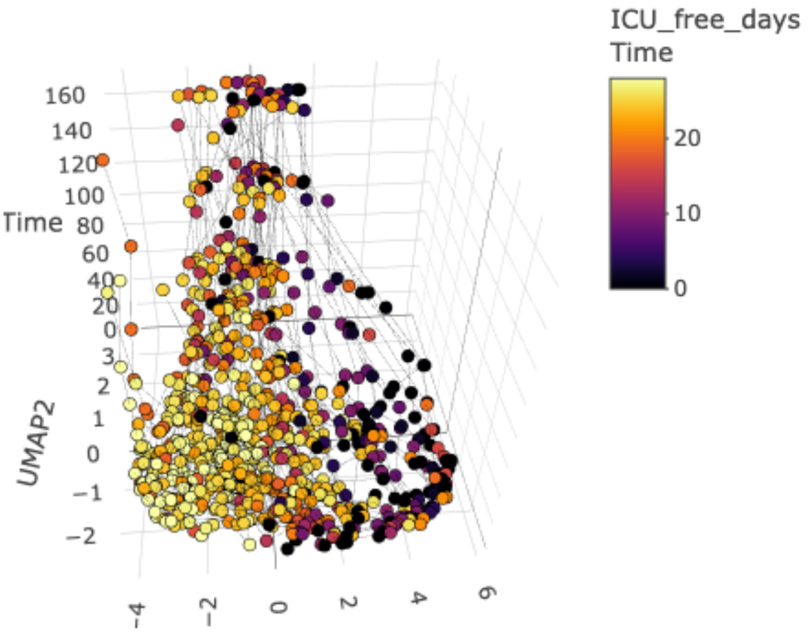

3D UMAP of proteomic patient-timepoints colored by association with ICU-free days (fewer = more severely injured)

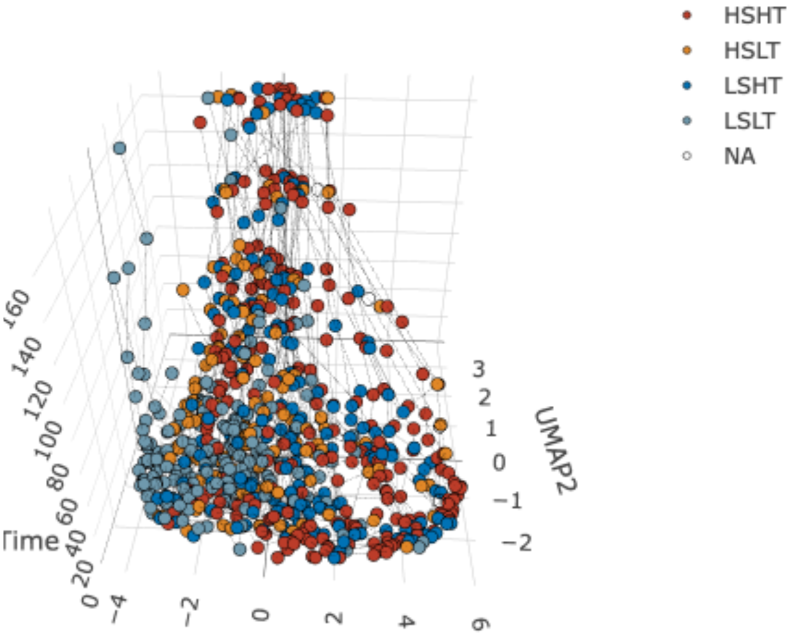

3D UMAP of proteomic patient-timepoints colored by association with shock and tissue injury

**Supplemental Figure 16 (Figure S16):** Proteomic patient state determination of cluster number, and basic characterization and distribution patient-timepoints associated with more severe- and less severe- injury.

#### *Supplementary FIGURE 17*

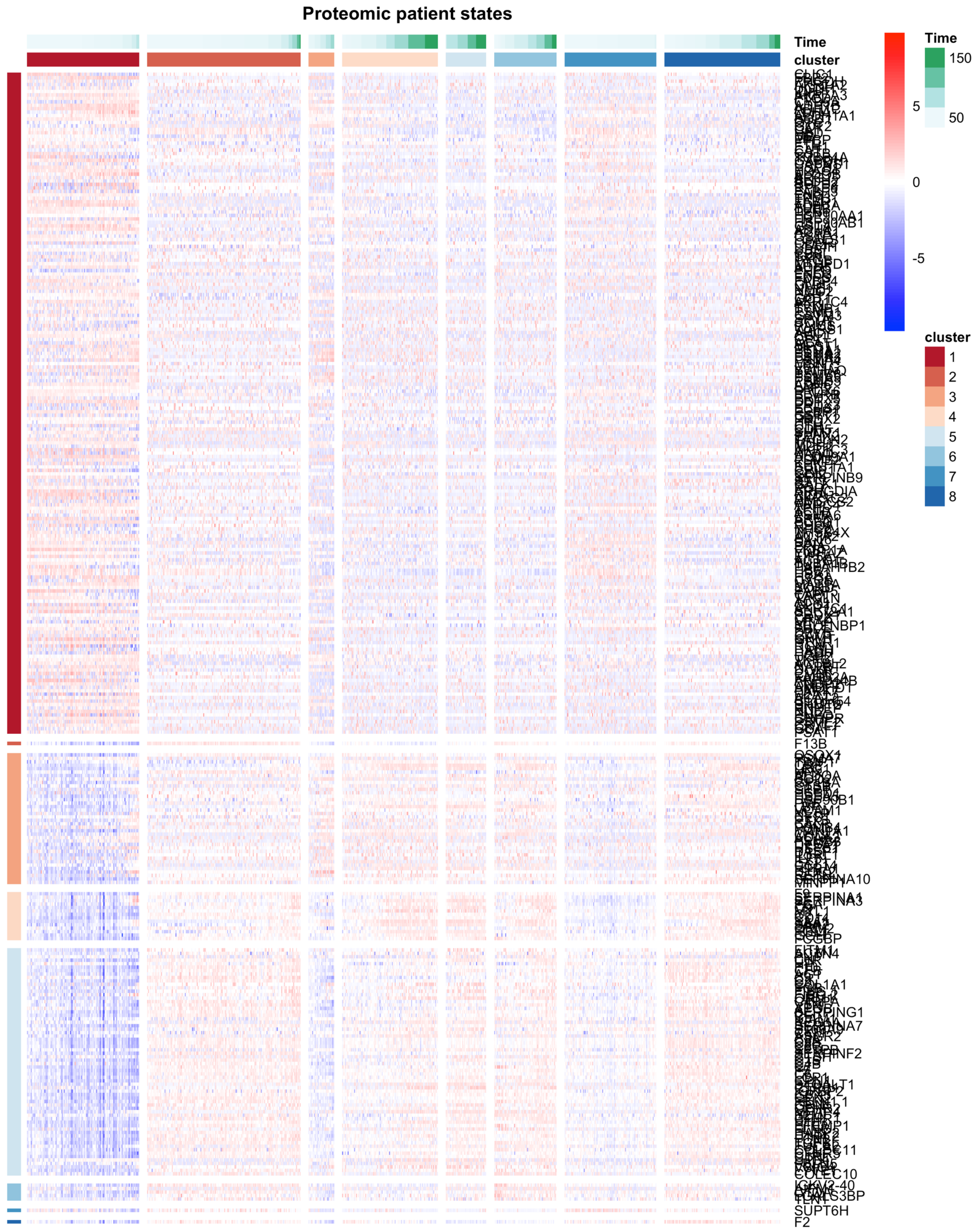

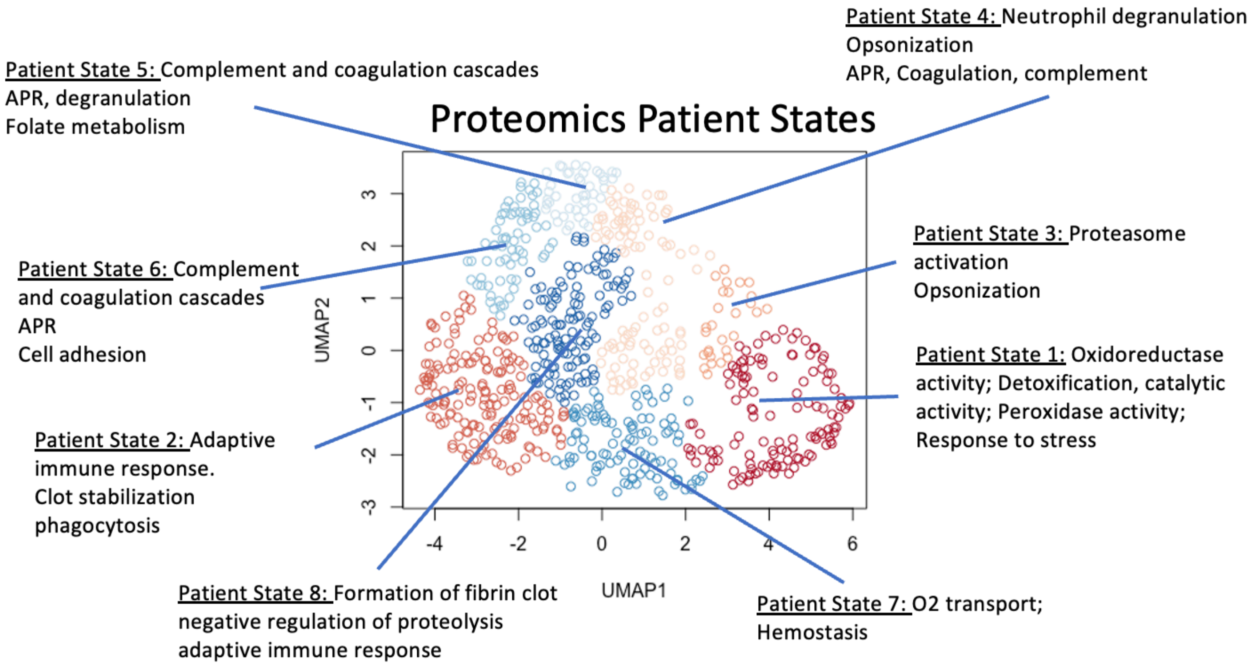

**Supplemental Figure 17 (Figure S17):** Heatmap of longitudinal proteomic patient state signature expression across all patient states. 2D UMAP of patient-timepoints colored by cluster, with overview of proteomic state as determined by pathway enrichment of analytes expressed at higher levels within each patient state.

#### *Supplementary FIGURE 18*

##### Vitals

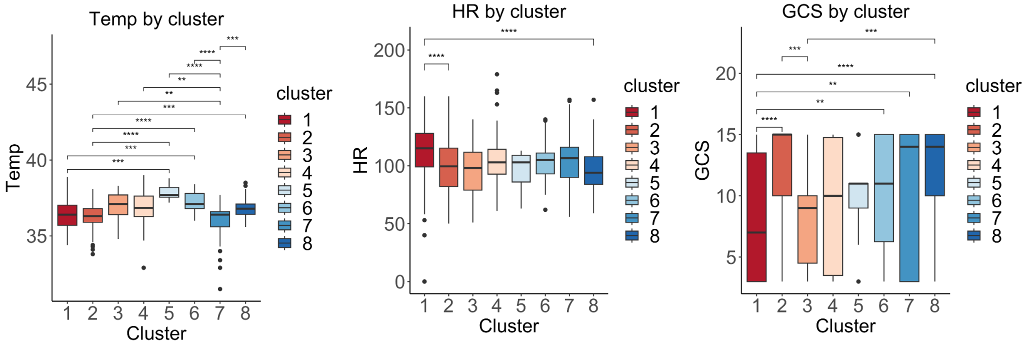

##### Blood chemistry

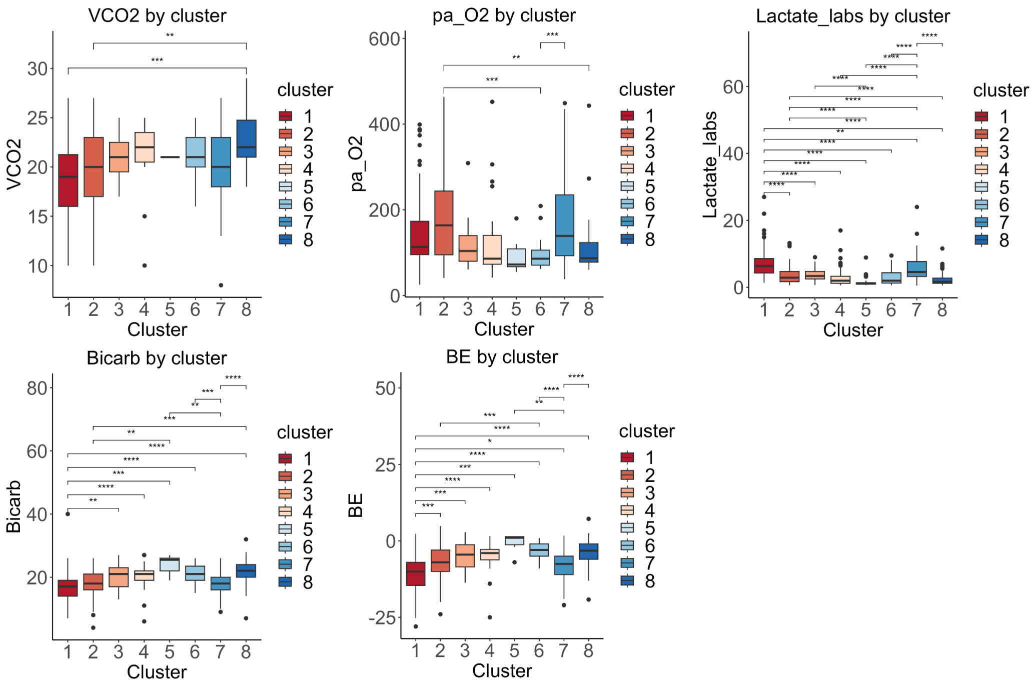

#### Blood products

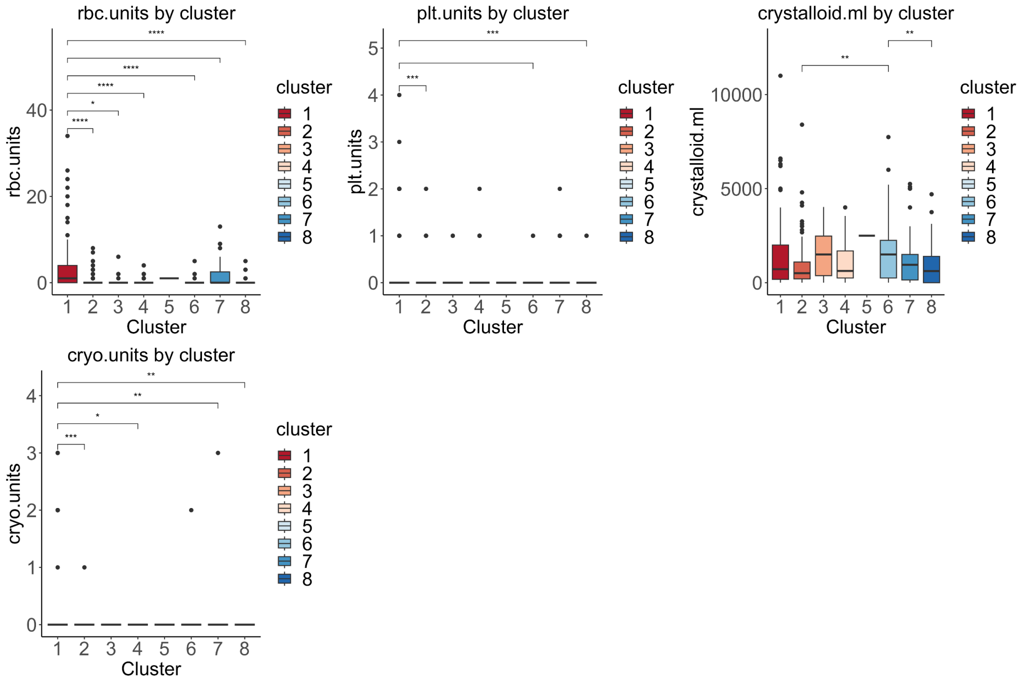

#### Blood count

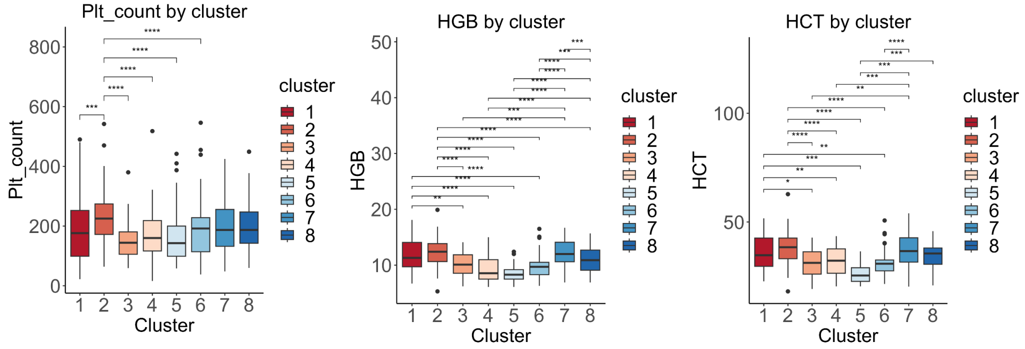

#### Clotting

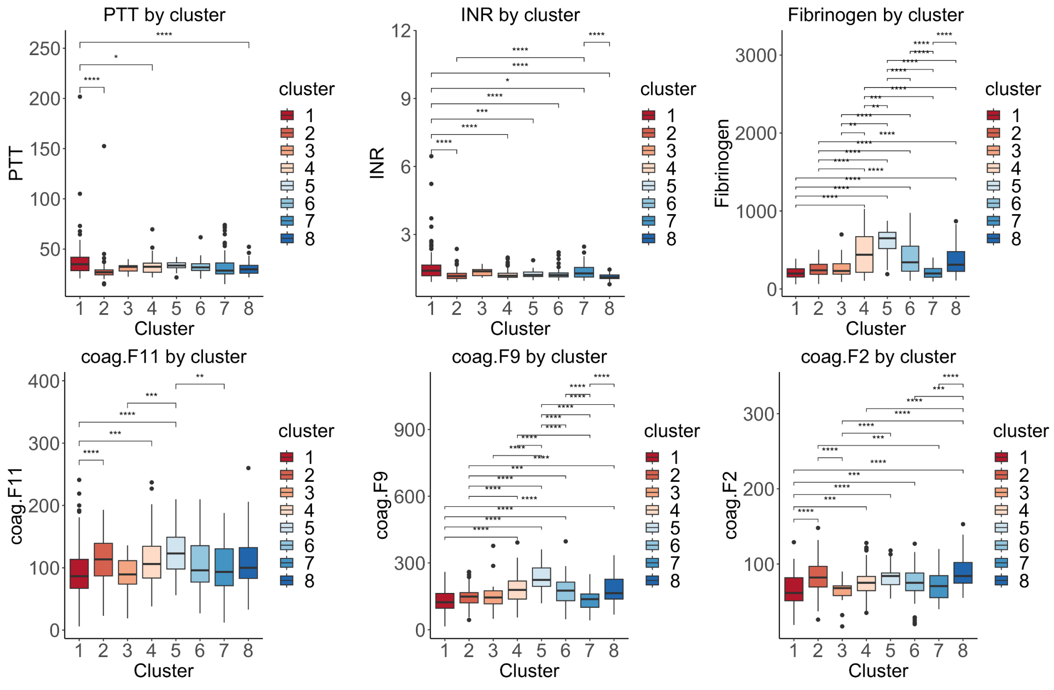

#### TEGs

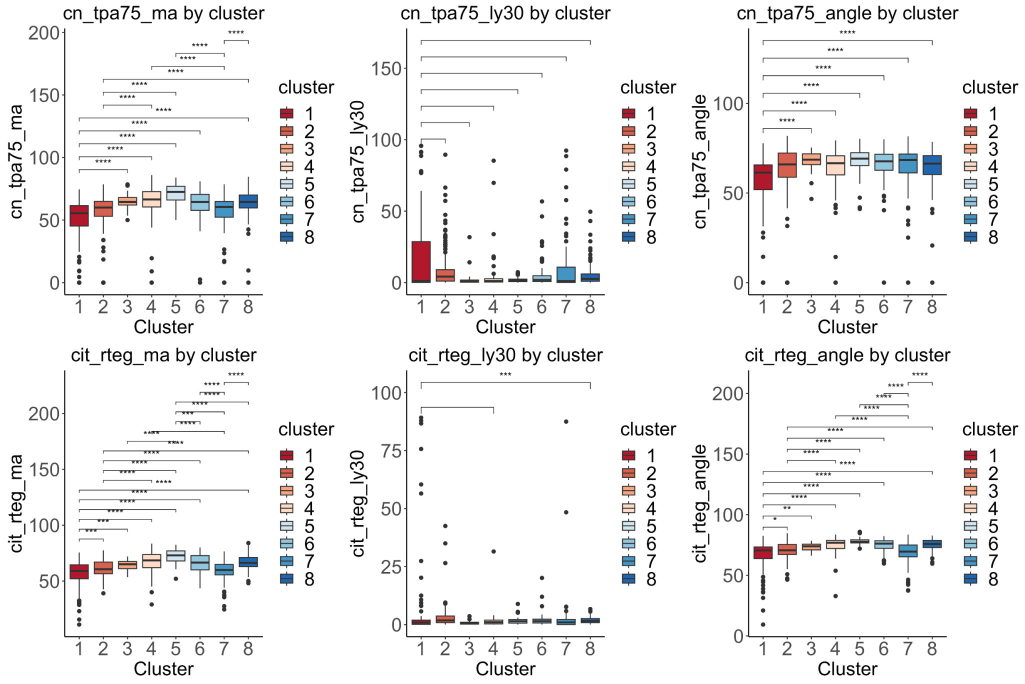

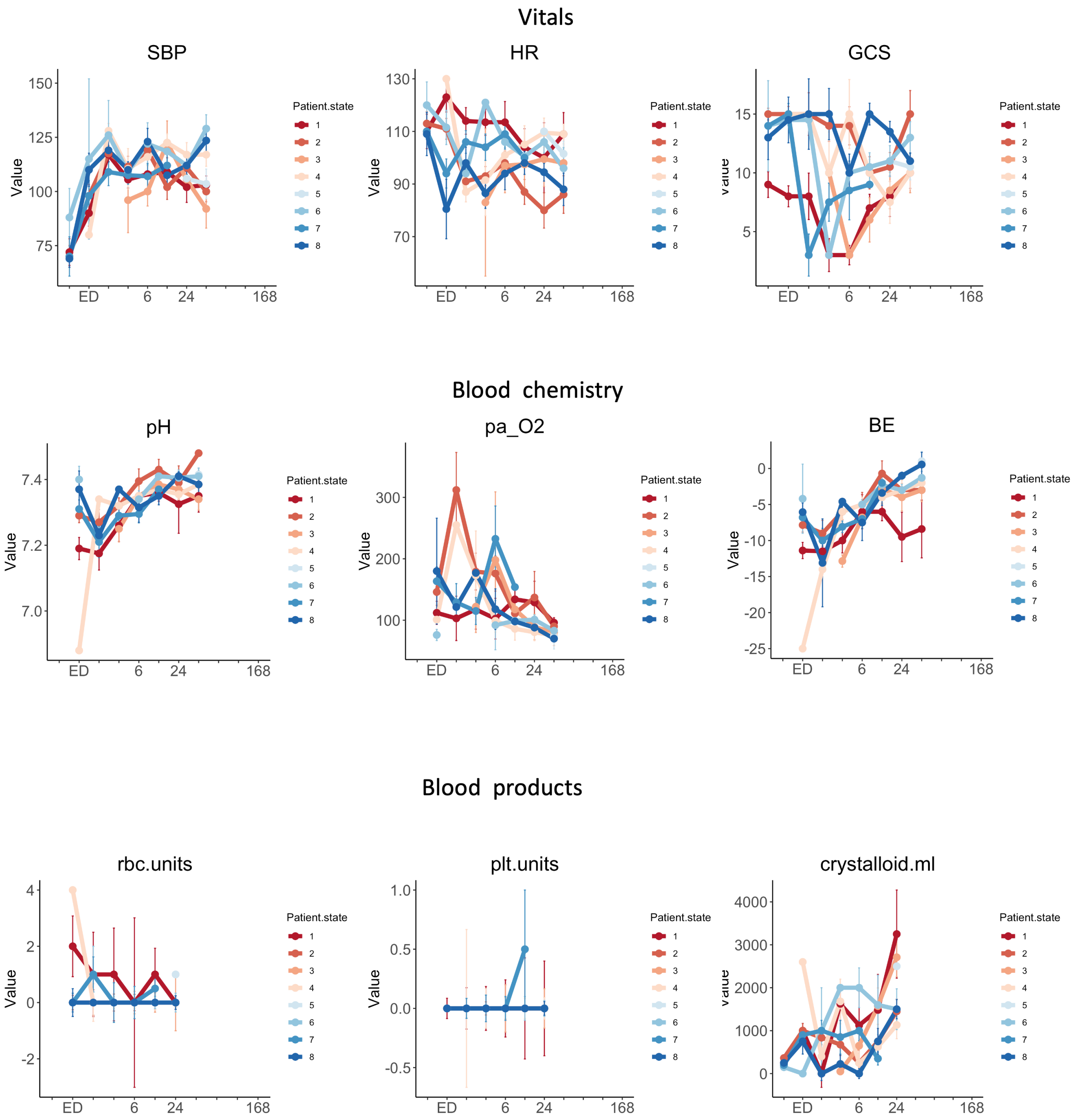

Vitals
SBP
HR
GCS
Blood chemistry
pH
pa_O2
BE
Blood products
rbc.units
plt.units
crystalloid.ml
Value
ED
6
24
168
Patient.state
1
2
3
4
5
6
7
8

#### Blood count

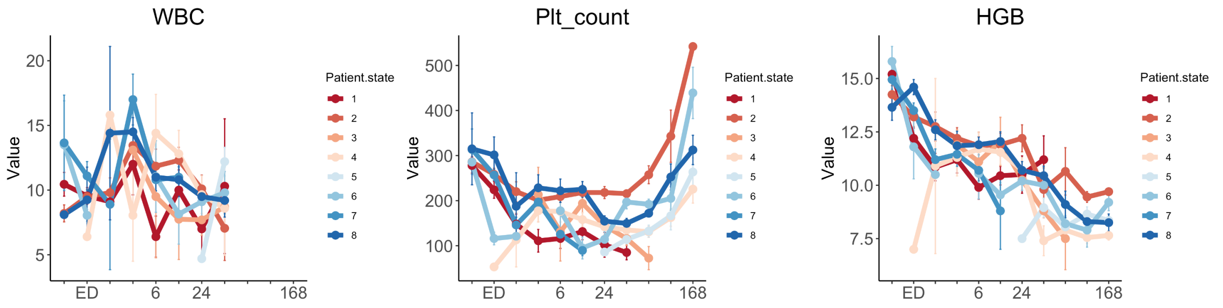

#### TEGs

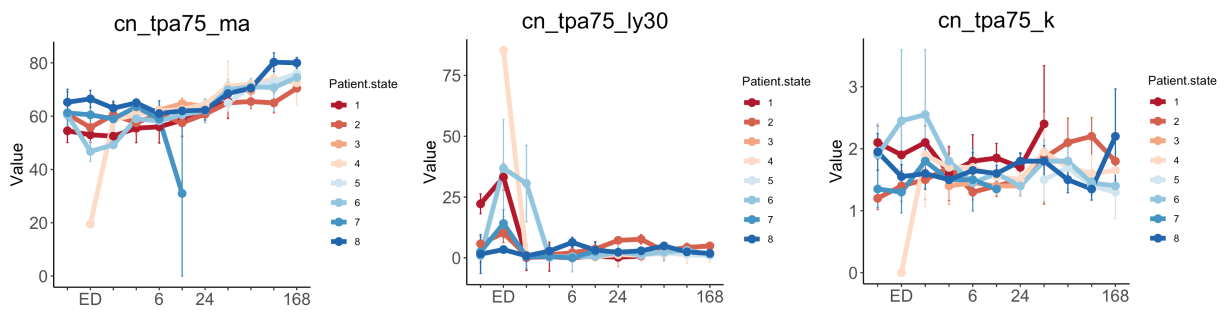

#### Clotting

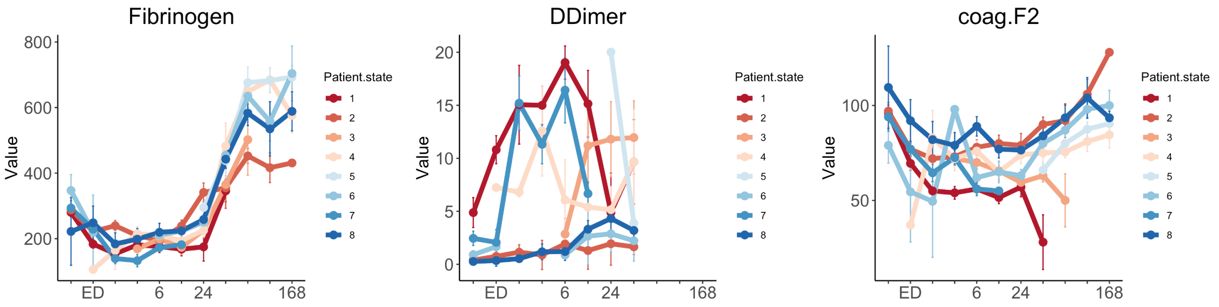

| Clinical assay | obs.tot | obs.groups | df.between | df.within | statistic | pvalue |
|---|---|---|---|---|---|---|
| Fibrinogen | 623 | 8 | 7 | 176.464722 | 48.3618672 | 7.47E-38 |
| HGB | 672 | 8 | 7 | 206.618983 | 39.1305483 | 1.15E-34 |
| coag.F5 | 637 | 8 | 7 | 193.665518 | 36.7795188 | 2.00E-32 |
| BD | 367 | 8 | 7 | 124.757513 | 36.8868773 | 1.56E-27 |
| cit_rteg_angle | 732 | 8 | 7 | 219.479054 | 27.1095219 | 1.20E-26 |
| cit_rteg_k | 730 | 8 | 7 | 218.932689 | 25.8041801 | 1.39E-25 |
| cit_rteg_g | 732 | 8 | 7 | 203.311733 | 26.0190158 | 2.97E-25 |
| cit_rteg_ma | 732 | 8 | 7 | 206.752613 | 25.3800773 | 7.29E-25 |
| Lactate_labs | 599 | 8 | 7 | 189.099647 | 26.0827402 | 8.50E-25 |
| coag.F9 | 637 | 8 | 7 | 184.891532 | 22.3477409 | 9.25E-22 |
| cn_ma | 730 | 8 | 7 | 216.923234 | 20.3496642 | 6.54E-21 |
| cn_tpa75_ma | 730 | 8 | 7 | 224.237535 | 19.4602578 | 2.92E-20 |
| INR | 650 | 8 | 7 | 192.749933 | 15.3652052 | 6.37E-16 |
| coag.F2 | 637 | 8 | 7 | 195.881733 | 15.3027193 | 6.48E-16 |
| cn_tpa75_ly30 | 703 | 8 | 7 | 225.42174 | 14.2849193 | 2.48E-15 |
| cn_tpa150_ly30 | 705 | 8 | 7 | 218.484444 | 13.9034634 | 7.19E-15 |
| DDimer | 417 | 8 | 7 | 73.9730277 | 18.9277968 | 3.13E-14 |
| cn_r | 730 | 8 | 7 | 195.480359 | 12.9566293 | 1.17E-13 |
| cn_tpa75_r | 730 | 8 | 7 | 200.414558 | 12.0642499 | 7.99E-13 |
| coag.plasm | 637 | 8 | 7 | 192.514661 | 11.8738866 | 1.51E-12 |
| coag.F8 | 637 | 8 | 7 | 189.848883 | 10.6588977 | 2.70E-11 |
| BE | 320 | 8 | 7 | 61.1902791 | 15.0782679 | 2.86E-11 |
| cn_tpa75_ly60 | 730 | 8 | 7 | 217.081184 | 10.0423739 | 7.30E-11 |
| SBP | 596 | 8 | 7 | 121.09716 | 10.8622179 | 1.40E-10 |
| dly30 | 730 | 8 | 7 | 221.192246 | 9.72592509 | 1.50E-10 |
| HCT | 495 | 8 | 7 | 98.8399022 | 11.3928289 | 1.60E-10 |
| Temp | 299 | 8 | 7 | 65.0395684 | 12.9533156 | 2.64E-10 |
| dly60 | 730 | 8 | 7 | 215.297414 | 9.47550575 | 3.04E-10 |
| pH | 303 | 8 | 7 | 63.5647922 | 11.3154227 | 3.23E-09 |
| GCS | 462 | 8 | 7 | 96.0856918 | 8.7550525 | 2.74E-08 |
| rbc.units | 491 | 7 | 6 | 132.494983 | 8.79551256 | 4.54E-08 |
| Bicarb | 303 | 8 | 7 | 59.6257869 | 9.02253775 | 1.53E-07 |
| cn_tpa75_angle | 730 | 8 | 7 | 224.385271 | 6.95127674 | 1.69E-07 |
| Plt_count | 662 | 8 | 7 | 199.337731 | 6.86137322 | 2.60E-07 |
| ffp.units | 560 | 7 | 6 | 136.454275 | 7.56597856 | 5.11E-07 |
| cit_rteg_ly30 | 732 | 8 | 7 | 222.536953 | 6.46593281 | 6.06E-07 |
| pa_O2 | 303 | 8 | 7 | 61.8571072 | 7.26986122 | 2.39E-06 |
| PTT | 645 | 8 | 7 | 203.198585 | 5.77102915 | 4.13E-06 |
| coag.F11 | 636 | 8 | 7 | 193.489214 | 5.60821685 | 6.66E-06 |

| Clinical assay | obs.tot | obs.groups | df.between | df.within | statistic | pvalue |
|---|---|---|---|---|---|---|
| cit_rteg_ly60 | 730 | 8 | 7 | 218.80178 | 5.48634764 | 7.97E-06 |
| cit_rteg_cl60 | 730 | 8 | 7 | 213.163612 | 5.47681283 | 8.40E-06 |
| cit_rteg_cl30 | 732 | 8 | 7 | 220.99956 | 5.41163499 | 9.60E-06 |
| coag.F7 | 637 | 8 | 7 | 191.530722 | 5.19348141 | 1.95E-05 |
| cn_angle | 730 | 8 | 7 | 212.365718 | 4.66962897 | 6.90E-05 |
| ion_Calcium | 295 | 8 | 7 | 93.5368952 | 4.4366986 | 0.000269535 |
| rteg_k | 118 | 6 | 5 | 17.6110668 | 8.14462018 | 0.000392442 |
| plt.units | 491 | 7 | 6 | 127.976065 | 4.00970994 | 0.001031629 |
| HR | 601 | 8 | 7 | 120.025345 | 3.75103916 | 0.001038429 |
| VCO2 | 170 | 7 | 6 | 41.1569012 | 4.30274321 | 0.001866152 |
| rteg_ma | 118 | 6 | 5 | 17.3864984 | 6.04168244 | 0.002051975 |
| rteg_angle | 118 | 6 | 5 | 17.2106468 | 5.87991787 | 0.002409321 |
| DBP | 563 | 8 | 7 | 121.719111 | 3.20114051 | 0.003796905 |
| rteg_g | 118 | 6 | 5 | 14.543458 | 4.67896979 | 0.009474372 |
| crystalloid.ml | 560 | 7 | 6 | 134.595529 | 2.78127694 | 0.013960616 |
| cn_k | 730 | 8 | 7 | 195.203937 | 2.49508237 | 0.017788481 |
| Potassium | 212 | 7 | 6 | 52.4939001 | 2.80201465 | 0.019338989 |
| Creatinine_labs | 161 | 7 | 6 | 40.3403285 | 2.8448694 | 0.020981819 |
| rteg_ly30 | 116 | 6 | 5 | 25.0532325 | 3.13597487 | 0.024722296 |
| cryo.units | 491 | 7 | 6 | 266.208178 | 2.3925504 | 0.028690718 |
| txa.mg | 491 | 7 | 6 | 191.354924 | 2.32184714 | 0.034611403 |
| cit_rteg_r | 730 | 8 | 7 | 197.039353 | 2.19958968 | 0.035853976 |
| cit_rteg_act | 732 | 8 | 7 | 209.724069 | 2.01825822 | 0.054217207 |
| pa_CO2 | 303 | 8 | 7 | 62.578566 | 1.90877264 | 0.083009169 |
| Glucose | 136 | 7 | 6 | 37.8925877 | 1.97168719 | 0.094177055 |
| cit_rteg_sp | 732 | 8 | 7 | 210.575177 | 1.6717794 | 0.117378175 |
| cn_ly30 | 730 | 8 | 7 | 207.357458 | 1.61748756 | 0.131959689 |
| cn_tpa75_k | 730 | 8 | 7 | 211.890914 | 1.28150071 | 0.260742145 |
| WBC | 444 | 8 | 7 | 92.2145595 | 1.11884381 | 0.358132556 |
| Calcium | 169 | 7 | 6 | 42.3053724 | 0.95613192 | 0.466174502 |
| rteg_sp | 118 | 6 | 5 | 13.0472176 | 0.63321841 | 0.678182088 |
| rteg_act | 118 | 6 | 5 | 12.9285081 | 0.2640918 | 0.924742936 |
| rteg_r | 118 | 6 | 5 | 13.2280173 | 0.18471416 | 0.963416344 |

**Supplemental Figure 18 (Figure S18):** Boxplots and line charts showing trends and statistical comparisons (ANOVA TukeyHSD posthoc) of clinical measurements per proteomics patient state.

#### *Supplementary FIGURE 19*

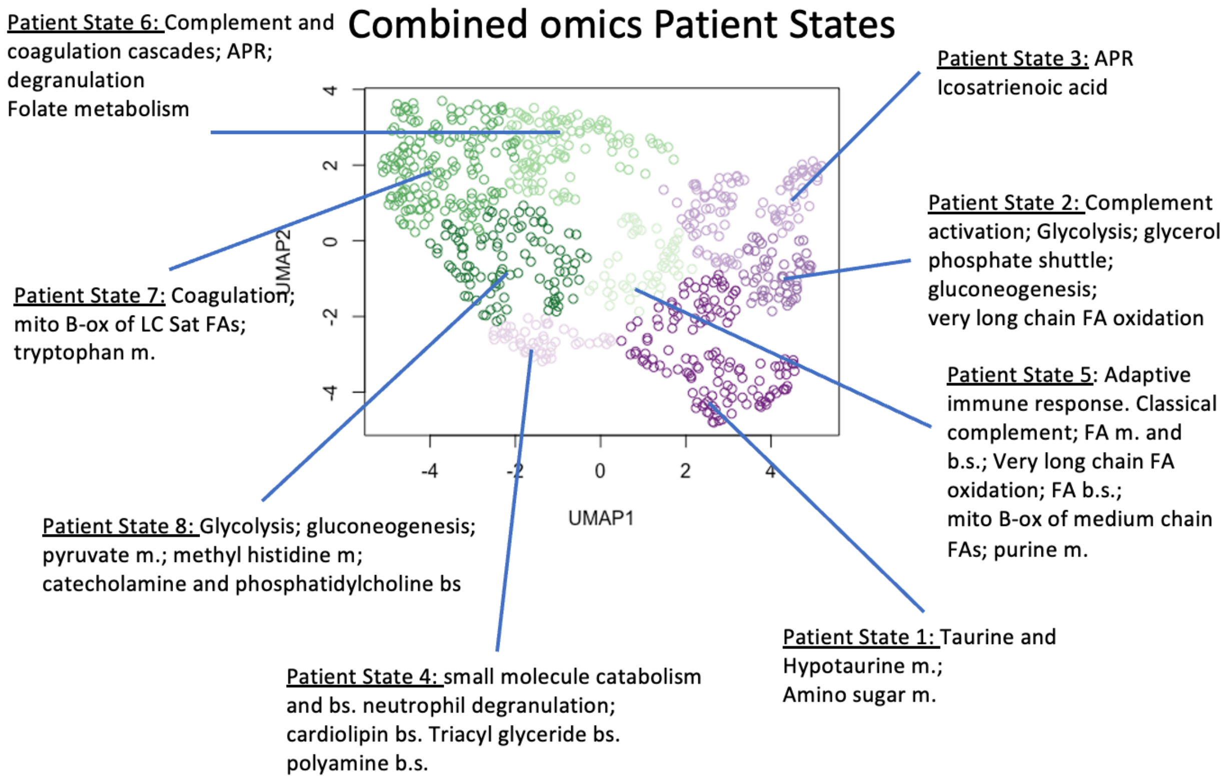

| PS | HGB | HCT | Fibrinogen | DDimer | PTT | INR | cn_ly30 | BE |
|---|---|---|---|---|---|---|---|---|
| 1 | 11.4 | 37.65 | 235 | 6.79 | 28.9 | 1.185 | 0.7 | -7 |
| 2 | 12.8 | 40 | 298 | 0.62 | 27.15 | 1.08 | 0.7 | -4 |
| 3 | 9.9 | 31.2 | 367 | 2.78 | 29.9 | 1.16 | 1.1 | -3 |
| 4 | 11.15 | 34.8 | 186 | 9.75 | 36.9 | 1.405 | 0.25 | -11.4 |
| 5 | 11.8 | 33.9 | 193 | 3.98 | 29.8 | 1.26 | 0.9 | -5 |
| 6 | 8.4 | 30.4 | 535 | 3.7 | 34.1 | 1.2 | 0.9 | -2.5 |
| 7 | 11.75 | 37.2 | 246.5 | 1.405 | 26.85 | 1.15 | 0.7 | -6.98 |
| 8 | 11.15 | 33.15 | 209.5 | 8.42 | 31.85 | 1.39 | 0.4 | -7.8 |

| PS | Lactate_labs | SBP | Temp | GCS | pH | Bicarb | pa_O2 | pa_CO2 | Plt_count | WBC |
|---|---|---|---|---|---|---|---|---|---|---|
| 1 | 3.25 | 103 | 36.5 | 11 | 7.33 | 19 | 97 | 38 | 173 | 9.85 |
| 2 | 1.9 | 101.5 | 36.5 | 15 | 7.35 | 20 | 127 | 37 | 258 | 8.9 |
| 3 | 1.4 | 112.5 | 37 | 11 | 7.39 | 22 | 90 | 36 | 208.5 | 10.7 |
| 4 | 7.3 | 88 | 36.25 | 4 | 7.23 | 15 | 126 | 40 | 176.5 | 9.15 |
| 5 | 3.3 | 111 | 36.8 | 11 | 7.34 | 21 | 88.5 | 37.5 | 186 | 11.95 |
| 6 | 1.5 | 111.5 | 37.3 | 10 | 7.39 | 23 | 92 | 37.5 | 141 | 9.35 |
| 7 | 3.2 | 108 | 36.5 | 14 | 7.315 | 19 | 134 | 36.5 | 207 | 10.85 |
| 8 | 5.3 | 103 | 36.5 | 8 | 7.32 | 17 | 113 | 33 | 152.5 | 9.5 |

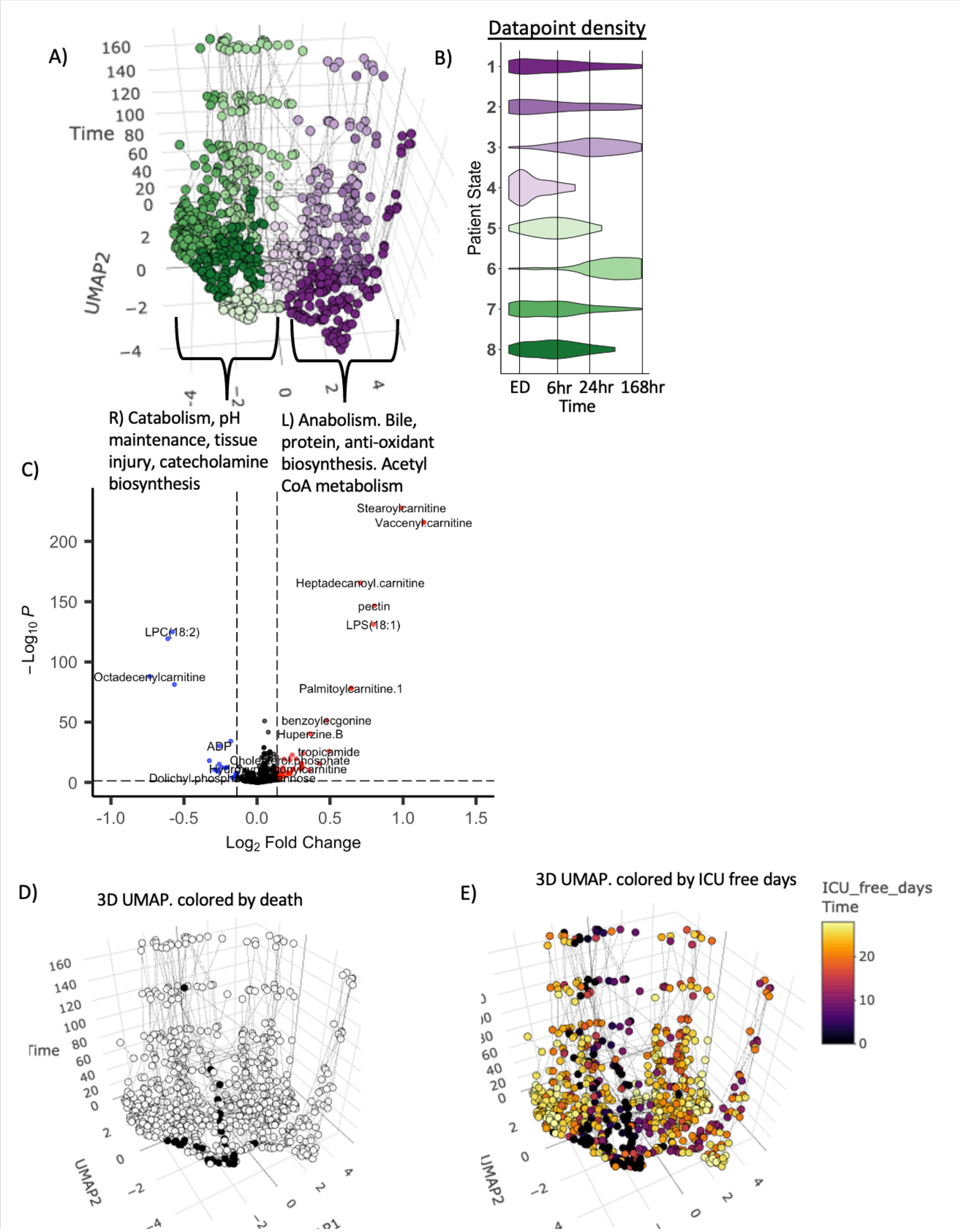
A)
Time
UMAP2
B)
Datapoint density
Patient State
ED
6hr
24hr
168hr
Time
R) Catabolism, pH maintenance, tissue injury, catecholamine biosynthesis
L) Anabolism. Bile, protein, anti-oxidant biosynthesis. Acetyl CoA metabolism
C)
Stearoylcarnitine
Vaccenylcarnitine
Heptadecanoyl.carnitine
pectin
LPS(18:1)
LPC(18:2)
Octadecenylcarnitine
Palmitoylcarnitine.1
benzoylecgonine
Huperzine.B
ADP
tropicamide
$-Log_{10}$ $P$
$Log_2$ Fold Change
D)
3D UMAP. colored by death
E)
3D UMAP. colored by ICU free days
ICU_free_days
Time

**Supplemental Figure 19 (Figure S19):** Combined proteomics and metabolomics patient states model, showing patient states, enrichment terms of proteins and metabolites elevated in each state, elevated metabolites in each meta state, median values of clinical measurements, clusters, and distribution of datapoints associated with death, ICU-free days, and S/T group.

#### *Supplementary FIGURE 20*

**Metabolomic trajectory signatures**

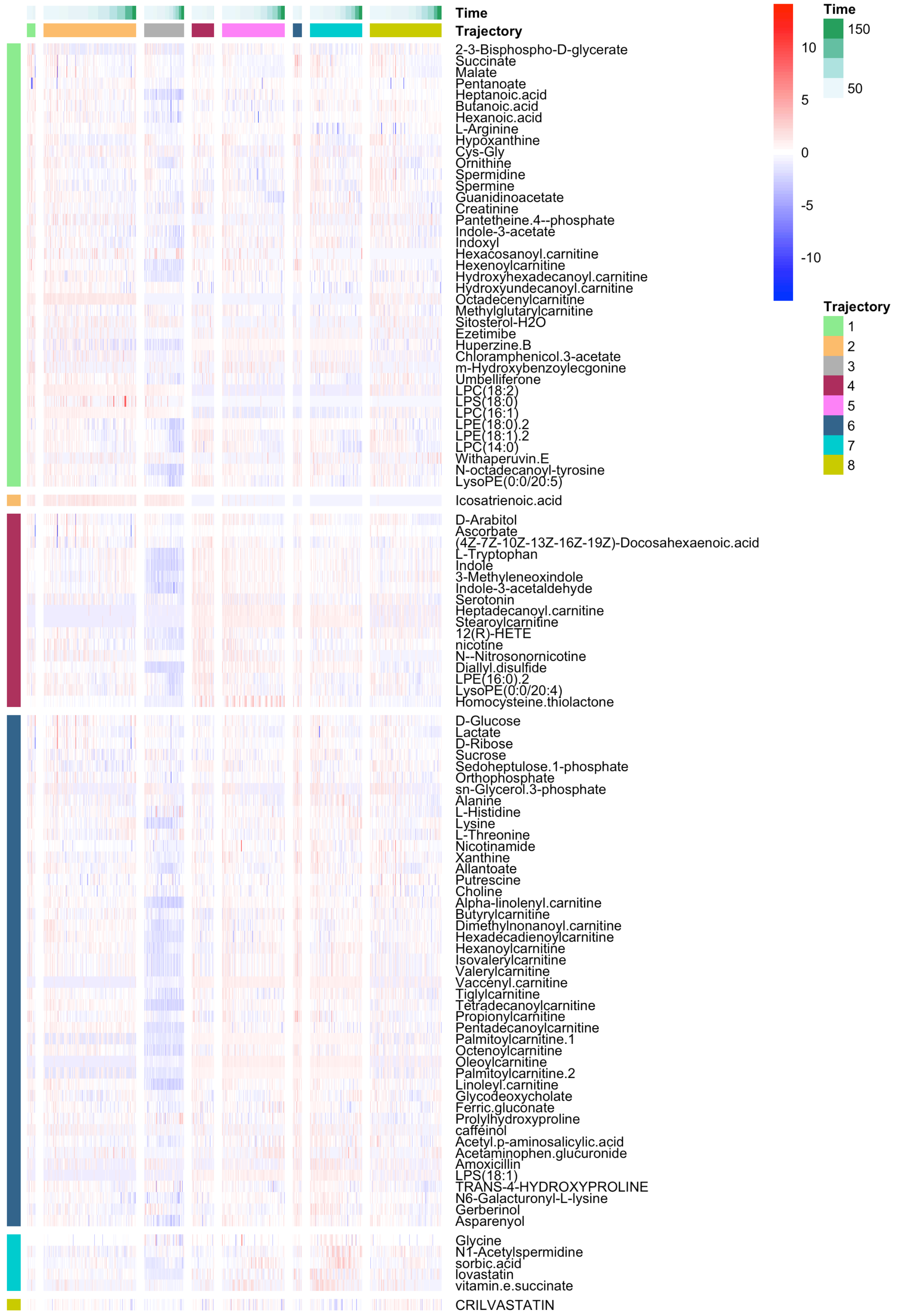

**Supplemental Figure 20 (Figure S20):** Metabolomics trajectories clustering results, and heatmap of metabolites expressed at high levels within each trajectory.

#### Supplementary FIGURE 21

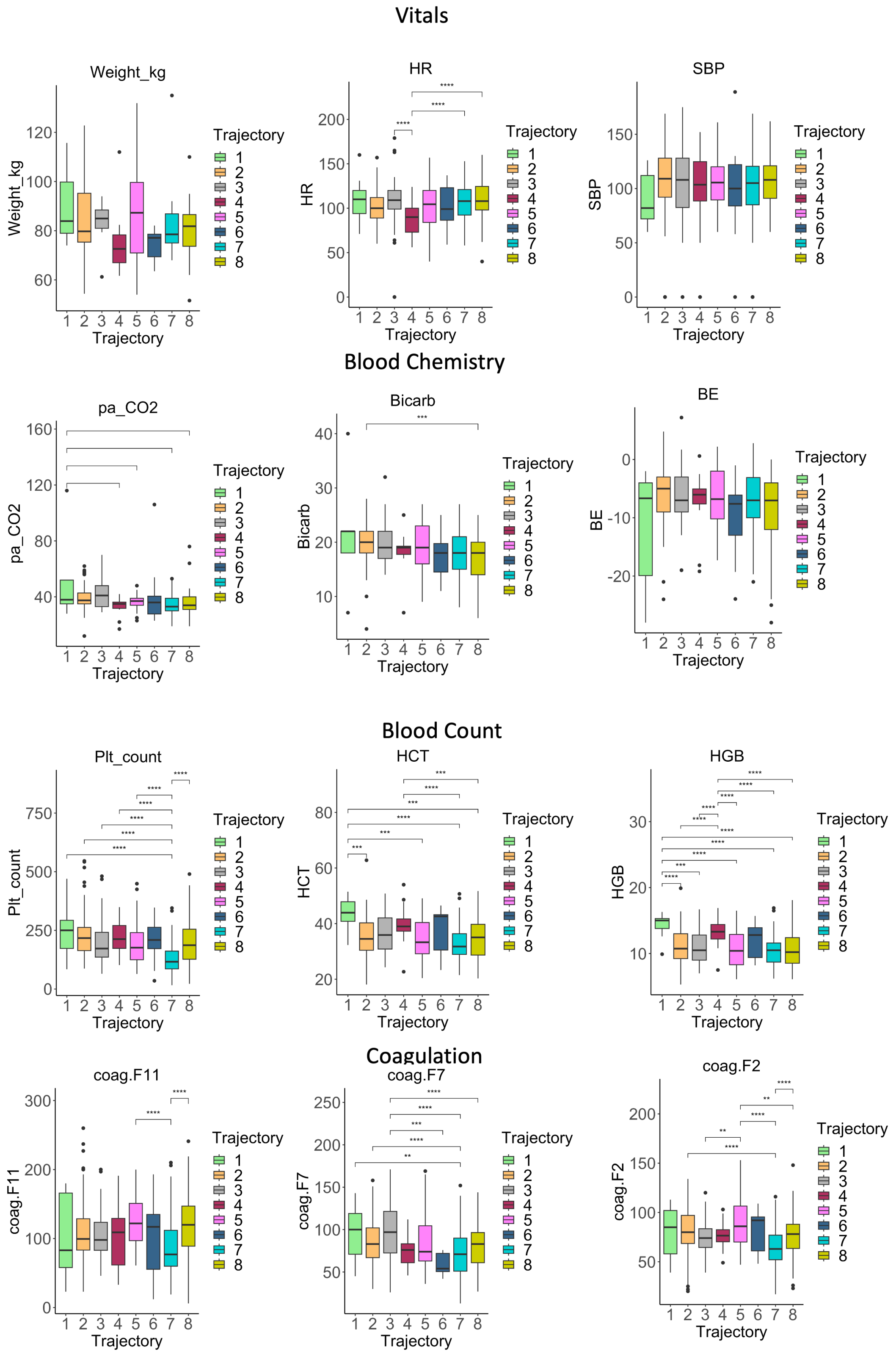

### Timeseries

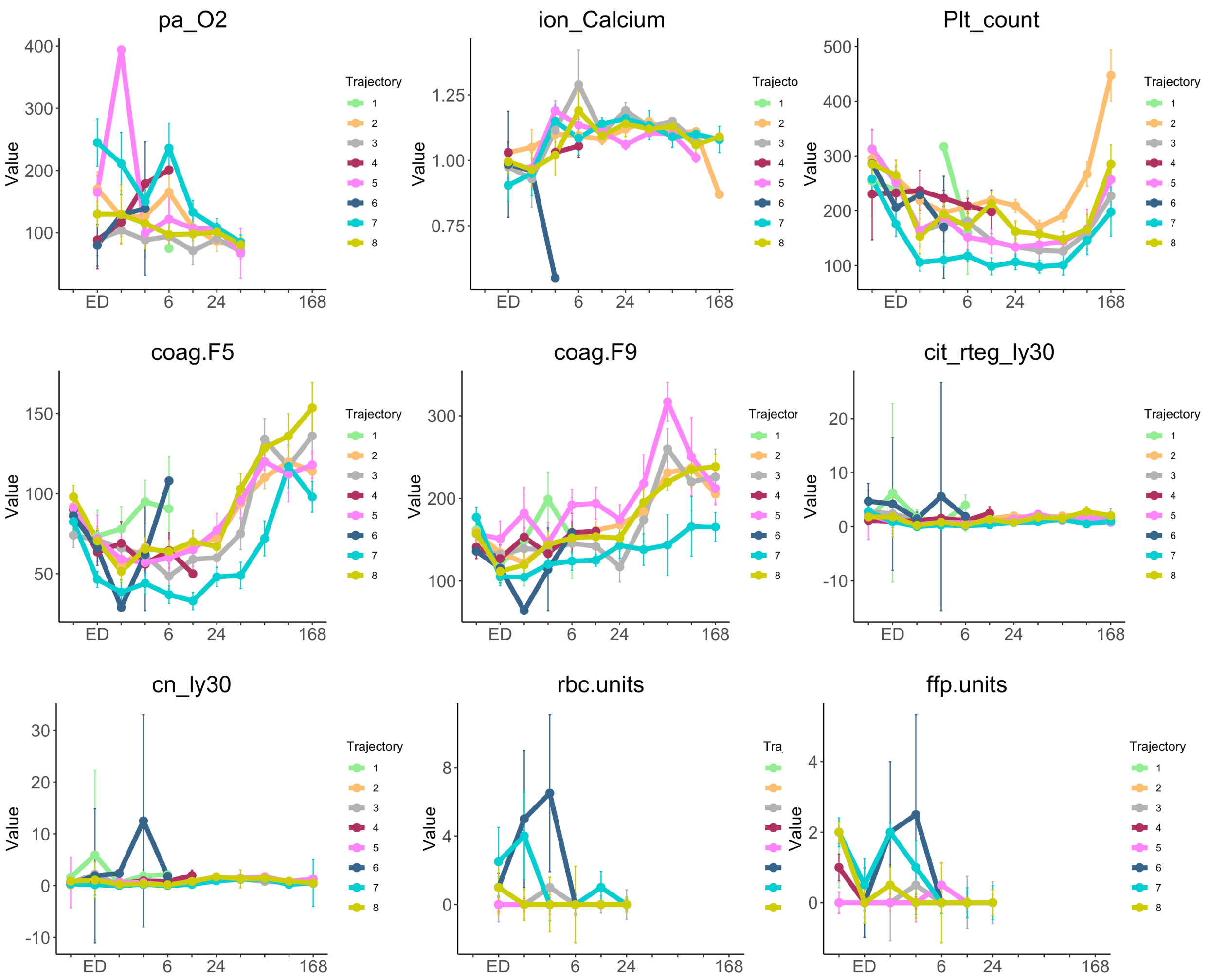
pa_O2
ion_Calcium
Plt_count
coag.F5
coag.F9
cit_rteg_ly30
cn_ly30
rbc.units
ffp.units
Value
ED
6
24
168
Trajectory
1
2
3
4
5
6
7
8

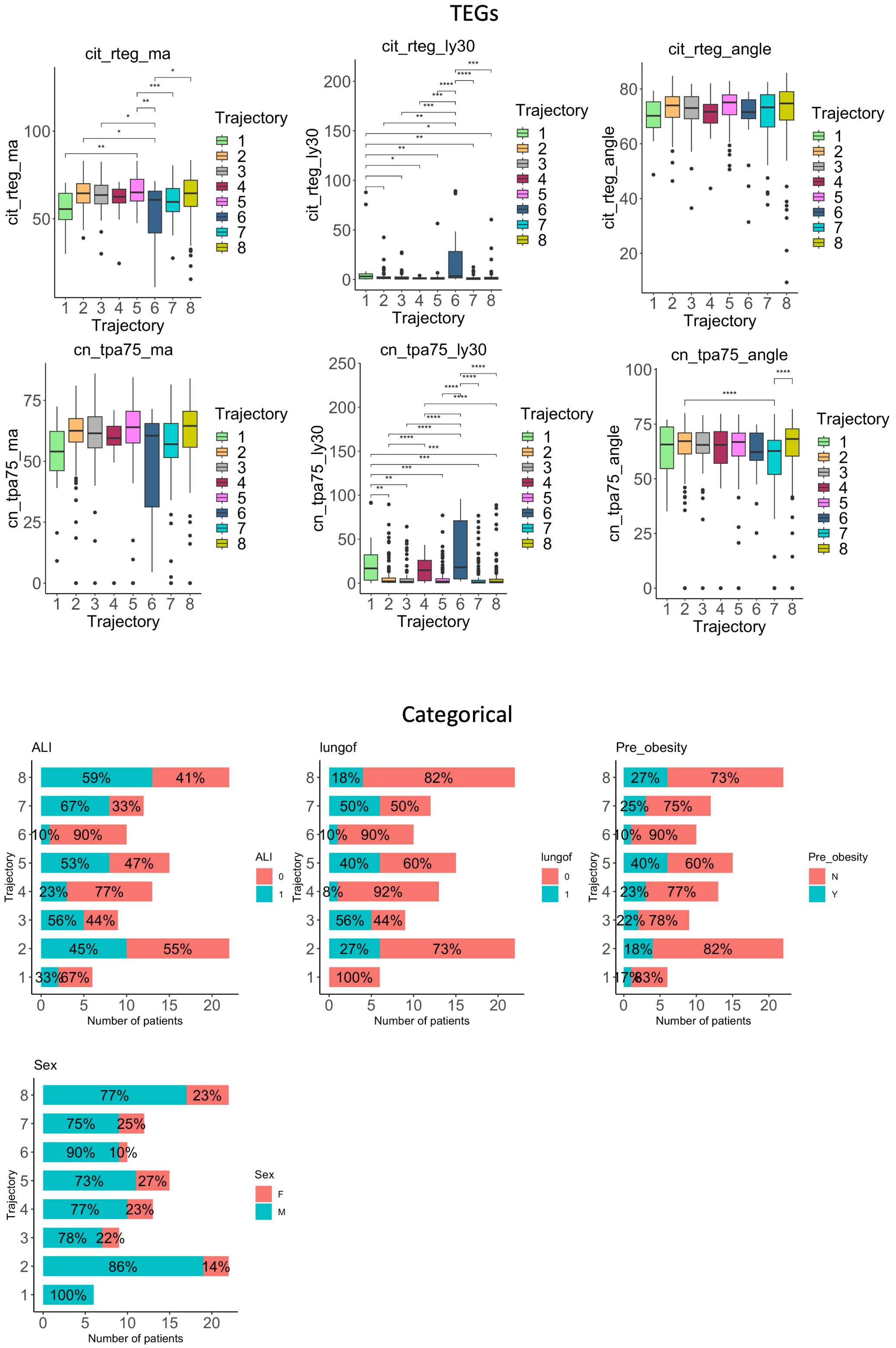

**Supplemental Figure 21 (Figure S21):** Boxplots, proportions, and line charts showing trends and statistical comparisons (ANOVA) of clinical measurements per metabolomics trajectories.

#### Supplementary FIGURE 22

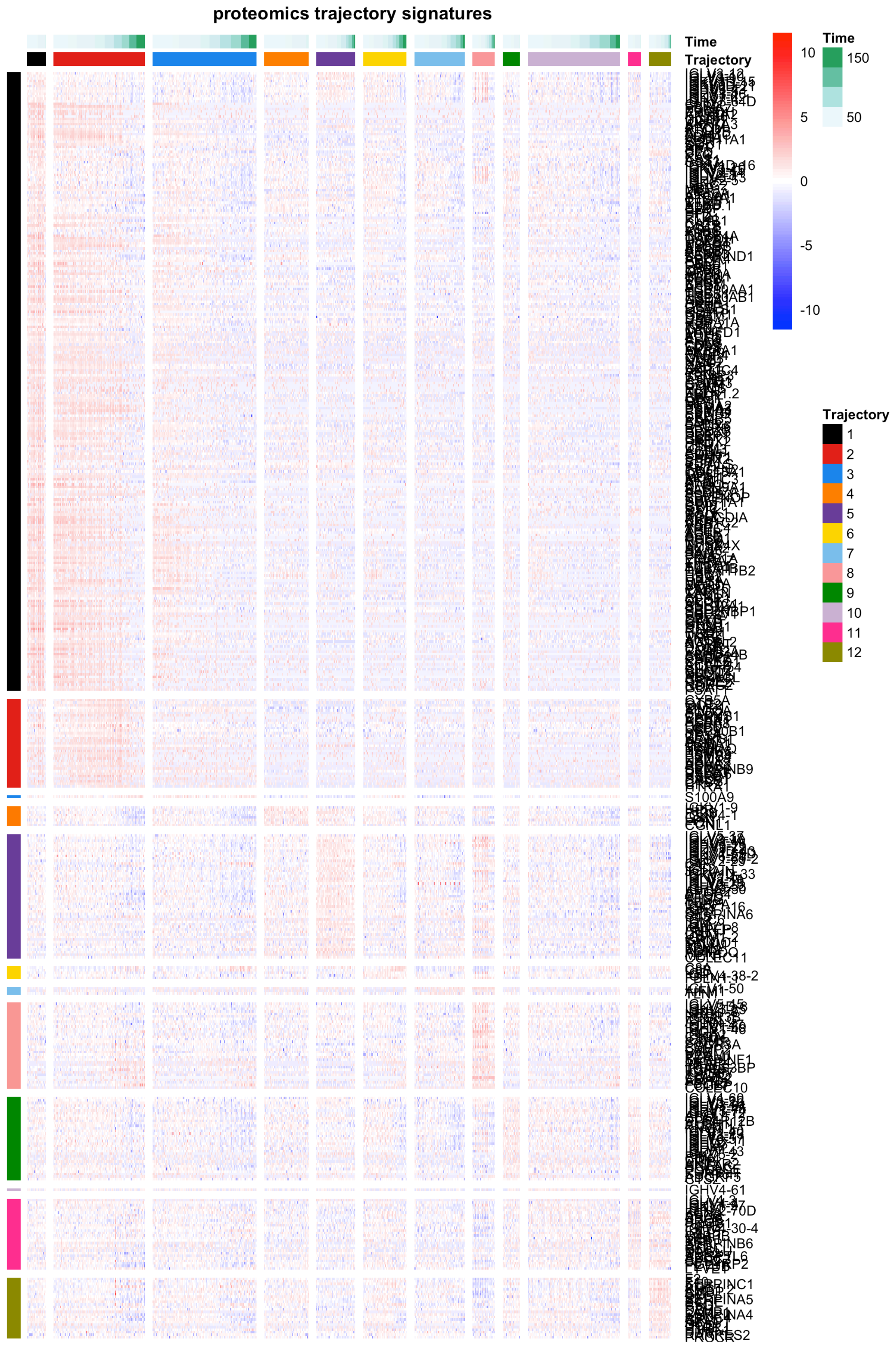

**Supplemental Figure 22 (Figure S22):** Proteomics trajectories clustering results, and heatmap of proteins expressed at high levels within each trajectory.

#### *Supplementary FIGURE 23*

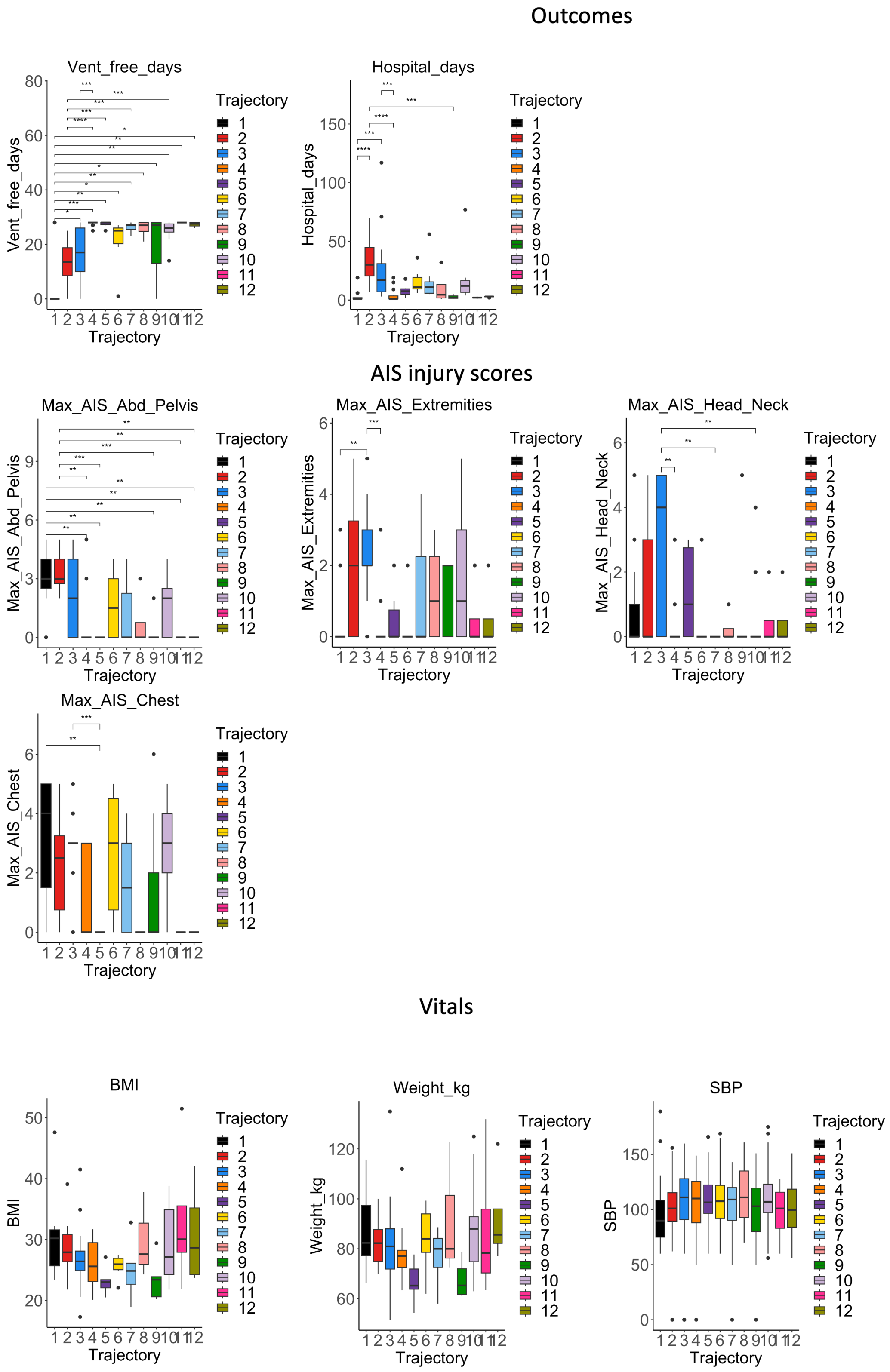

#### Blood Chemistry

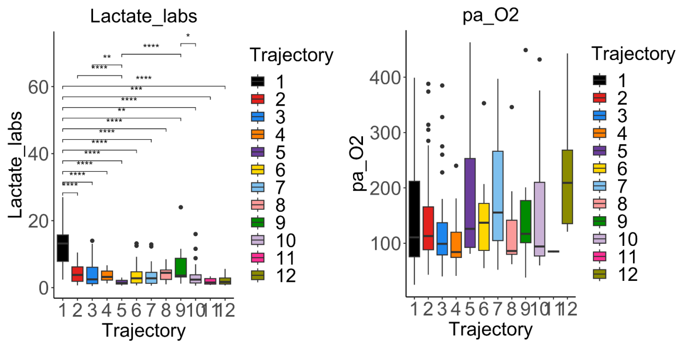

#### Blood Count

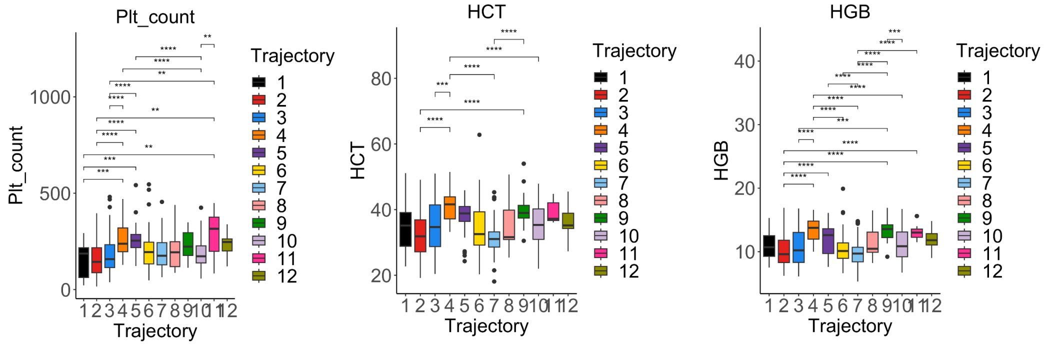

#### TEGs

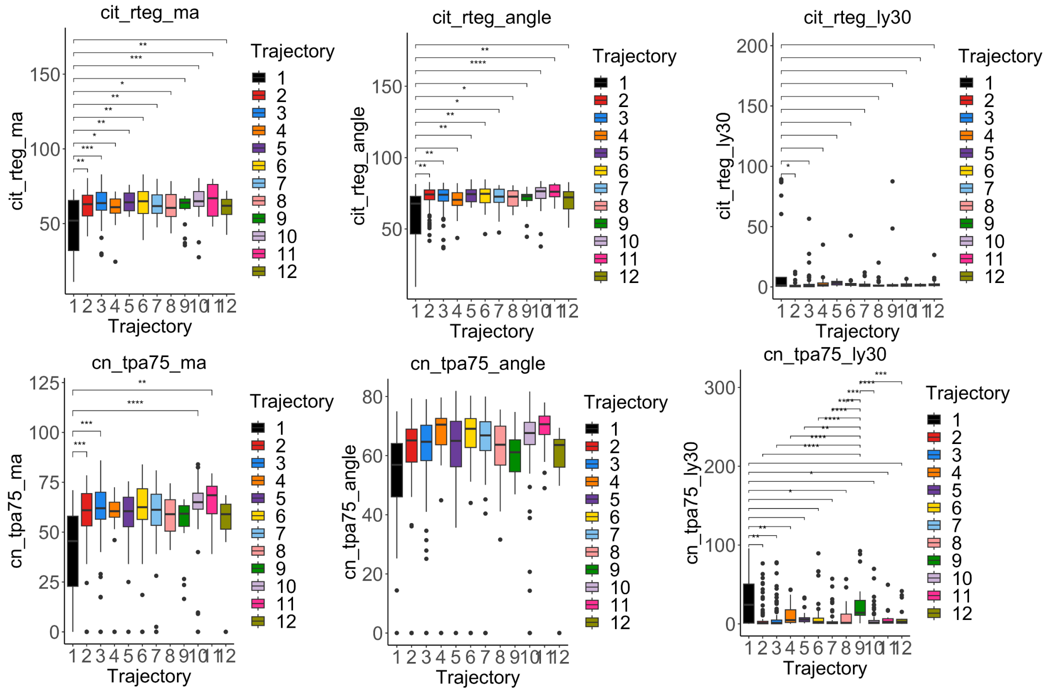

### Categorical

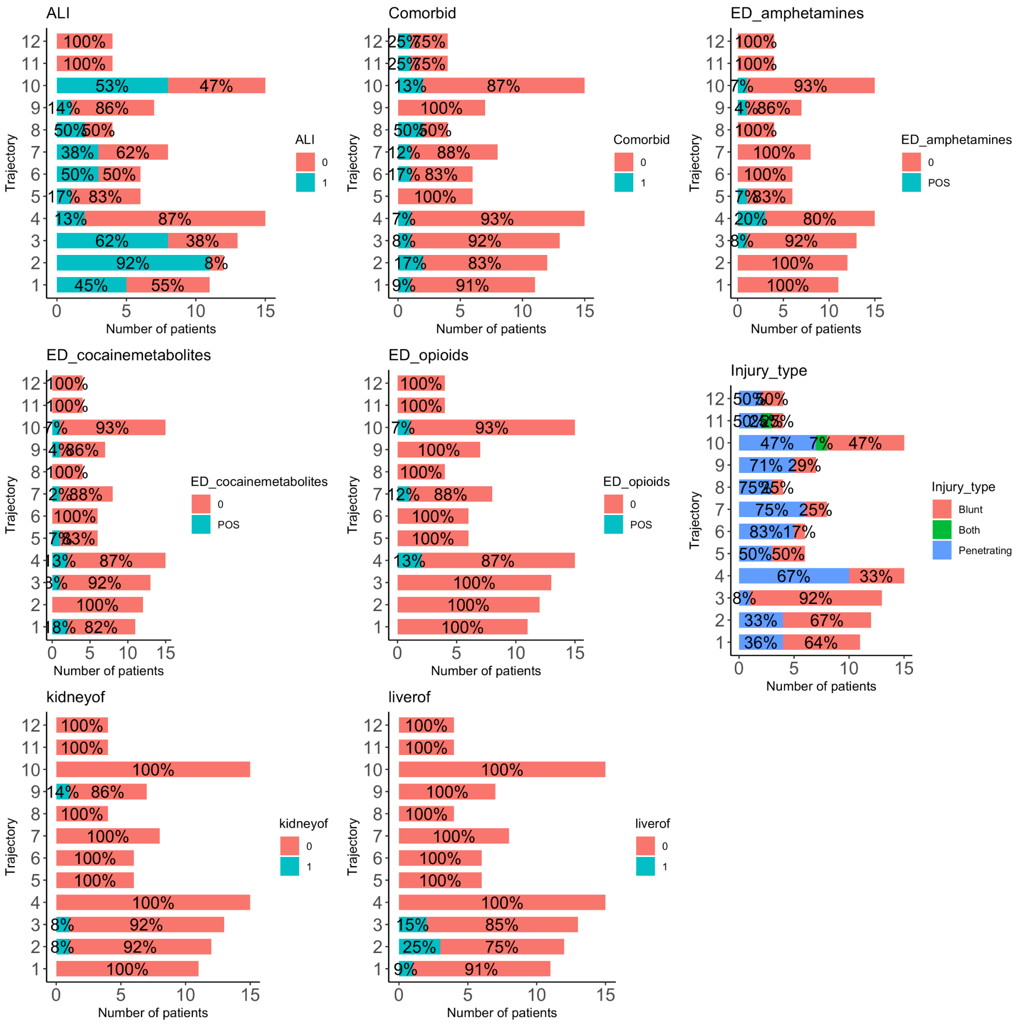

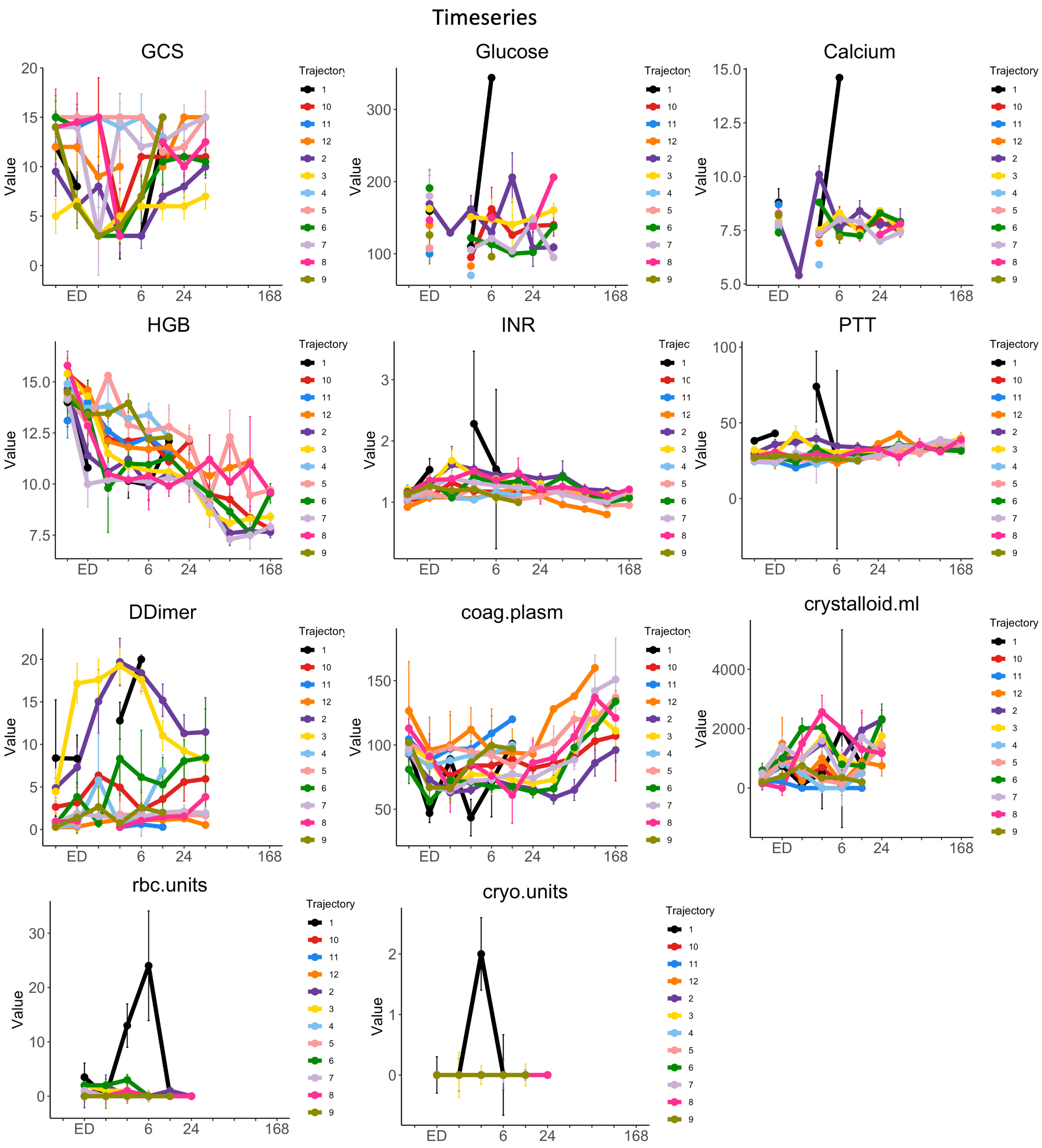
Timeseries
GCS
Glucose
Calcium
HGB
INR
PTT
DDimer
coag.plasm
crystalloid.ml
rbc.units
cryo.units
Value
ED
6
24
168
Trajectory
1
10
11
12
2
3
4
5
6
7
8
9

**Supplemental Figure 23 (Figure S23):** Boxplots, proportions, and line charts showing trends and statistical comparisons (ANOVA) of clinical measurements per proteomics trajectories.

#### Supplementary FIGURE 24

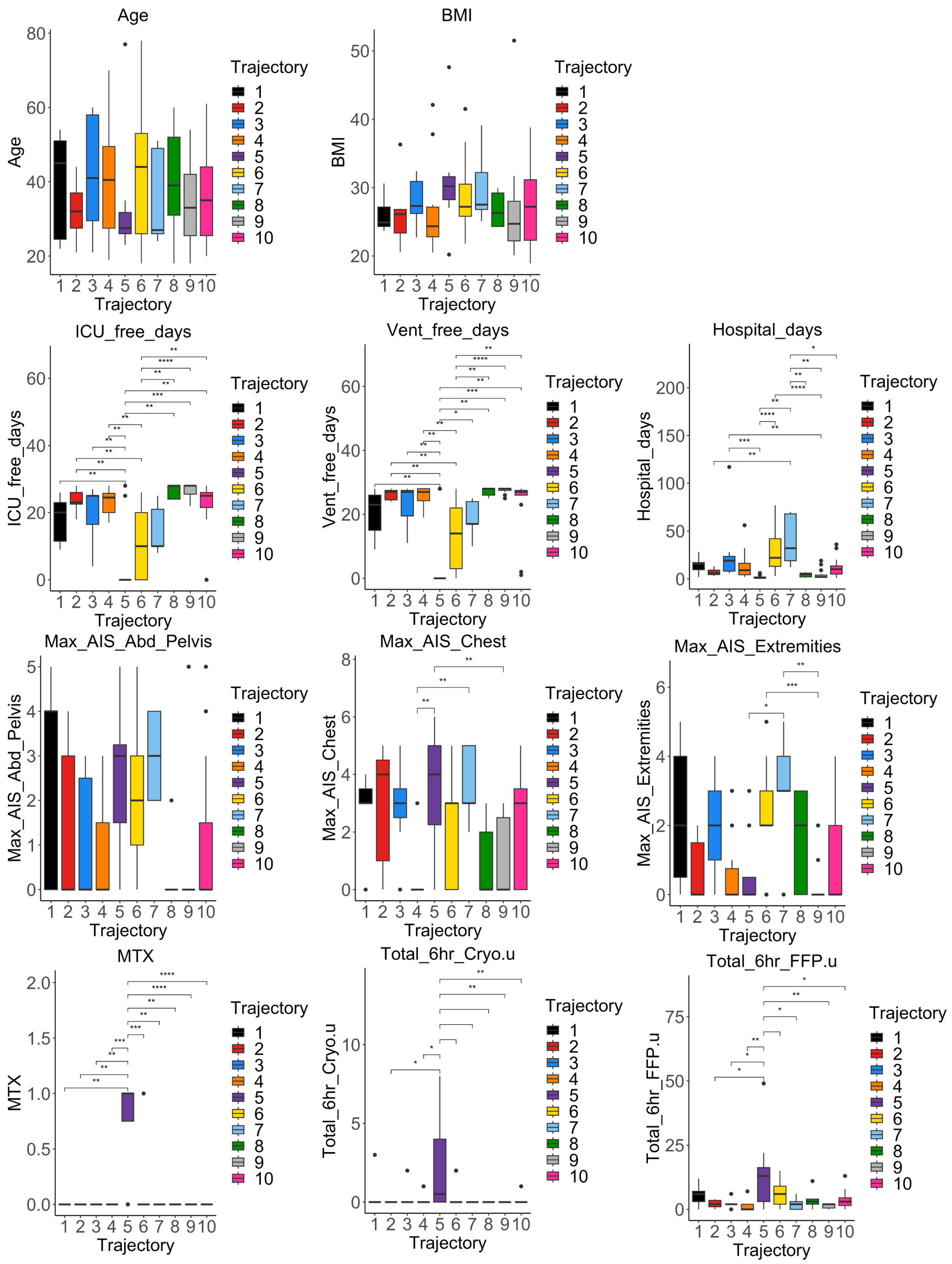

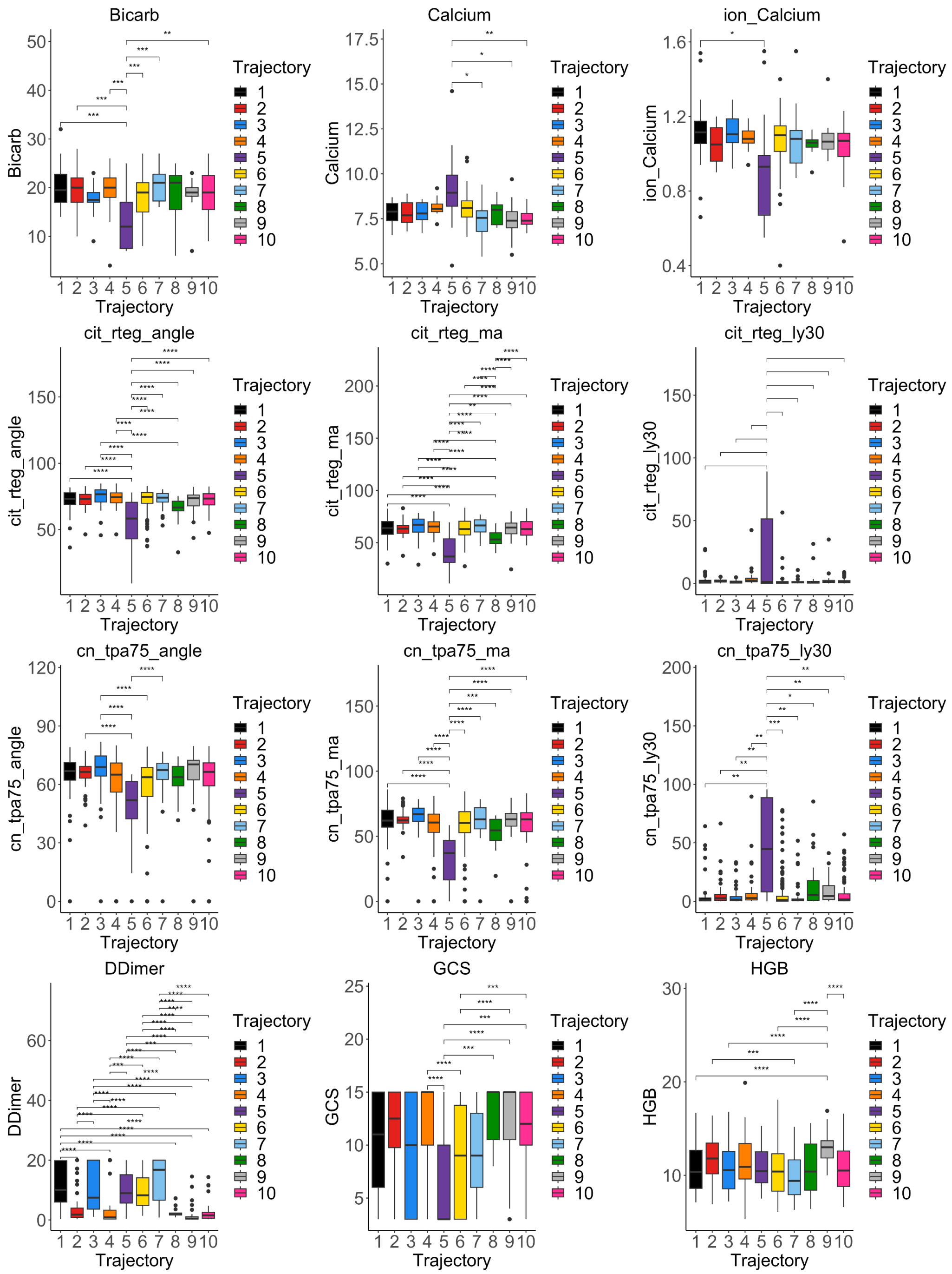
Bicarb
Calcium
ion_Calcium
cit_rteg_angle
cit_rteg_ma
cit_rteg_ly30
cn_tpa75_angle
cn_tpa75_ma
cn_tpa75_ly30
DDimer
GCS
HGB
Trajectory
1
2
3
4
5
6
7
8
9
10

### Categorical

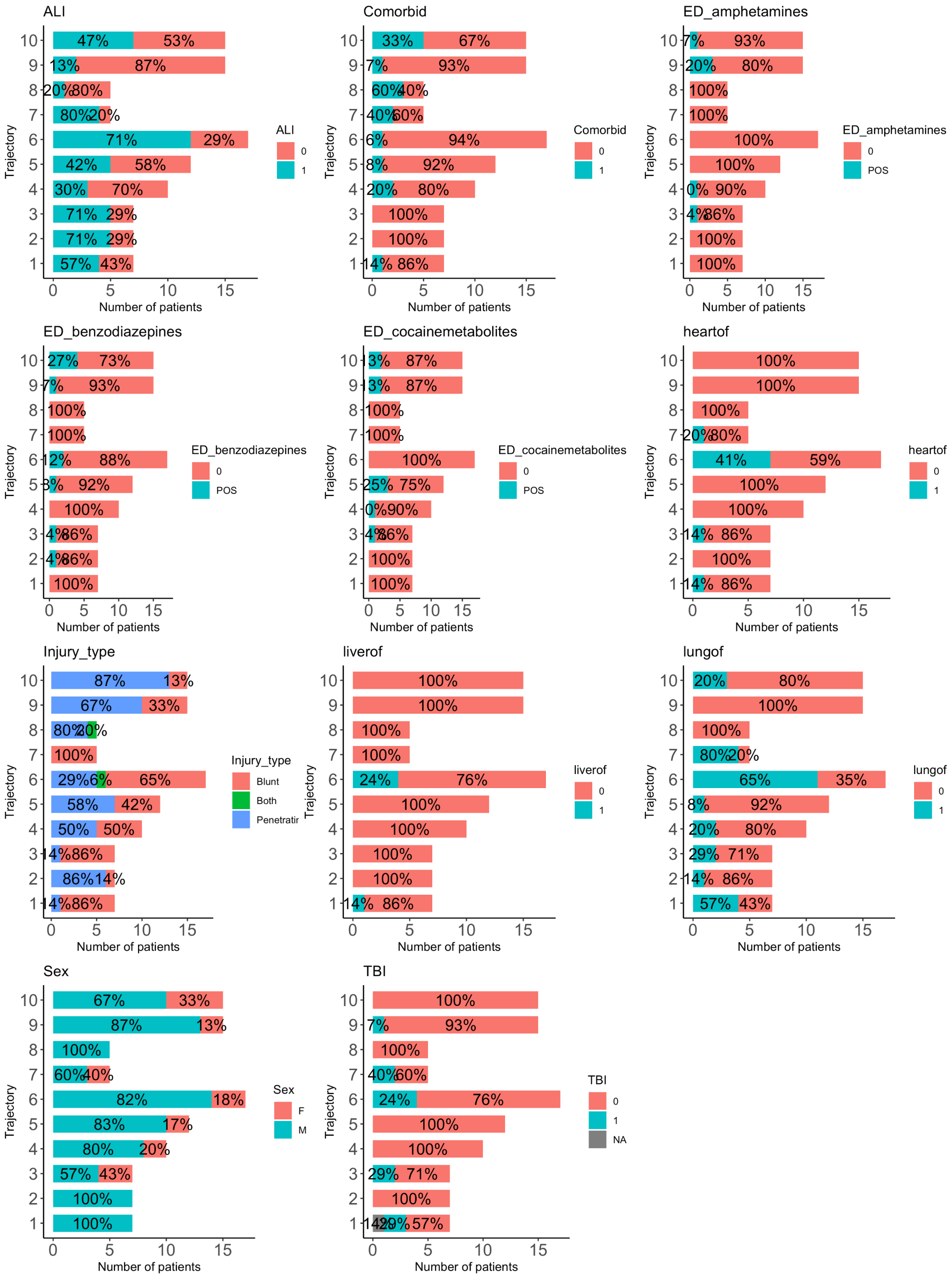

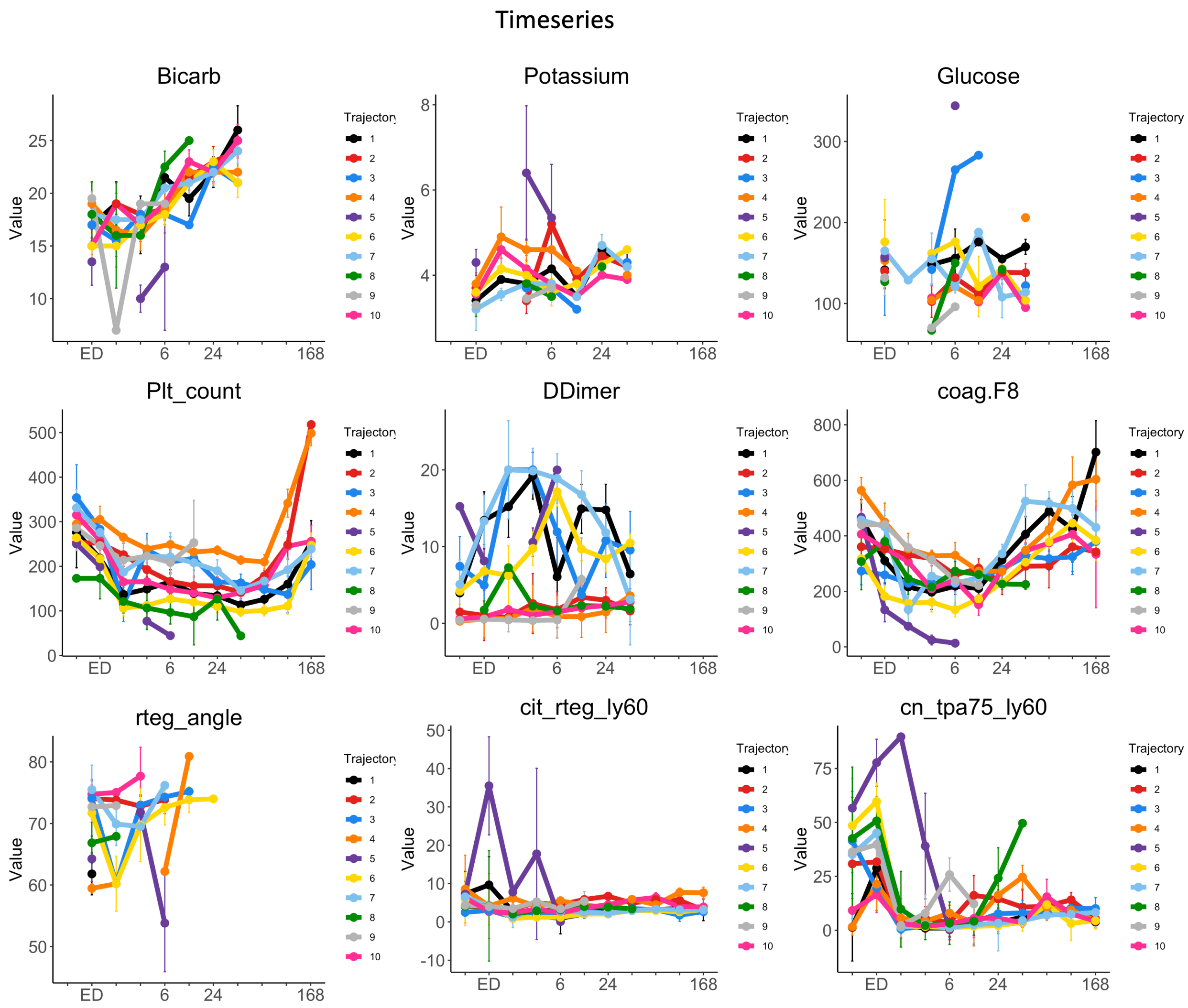

Timeseries
Bicarb
Potassium
Glucose
Plt_count
DDimer
coag.F8
rteg_angle
cit_rteg_ly60
cn_tpa75_ly60
Value
ED
6
24
168
Trajectory
1
2
3
4
5
6
7
8
9
10

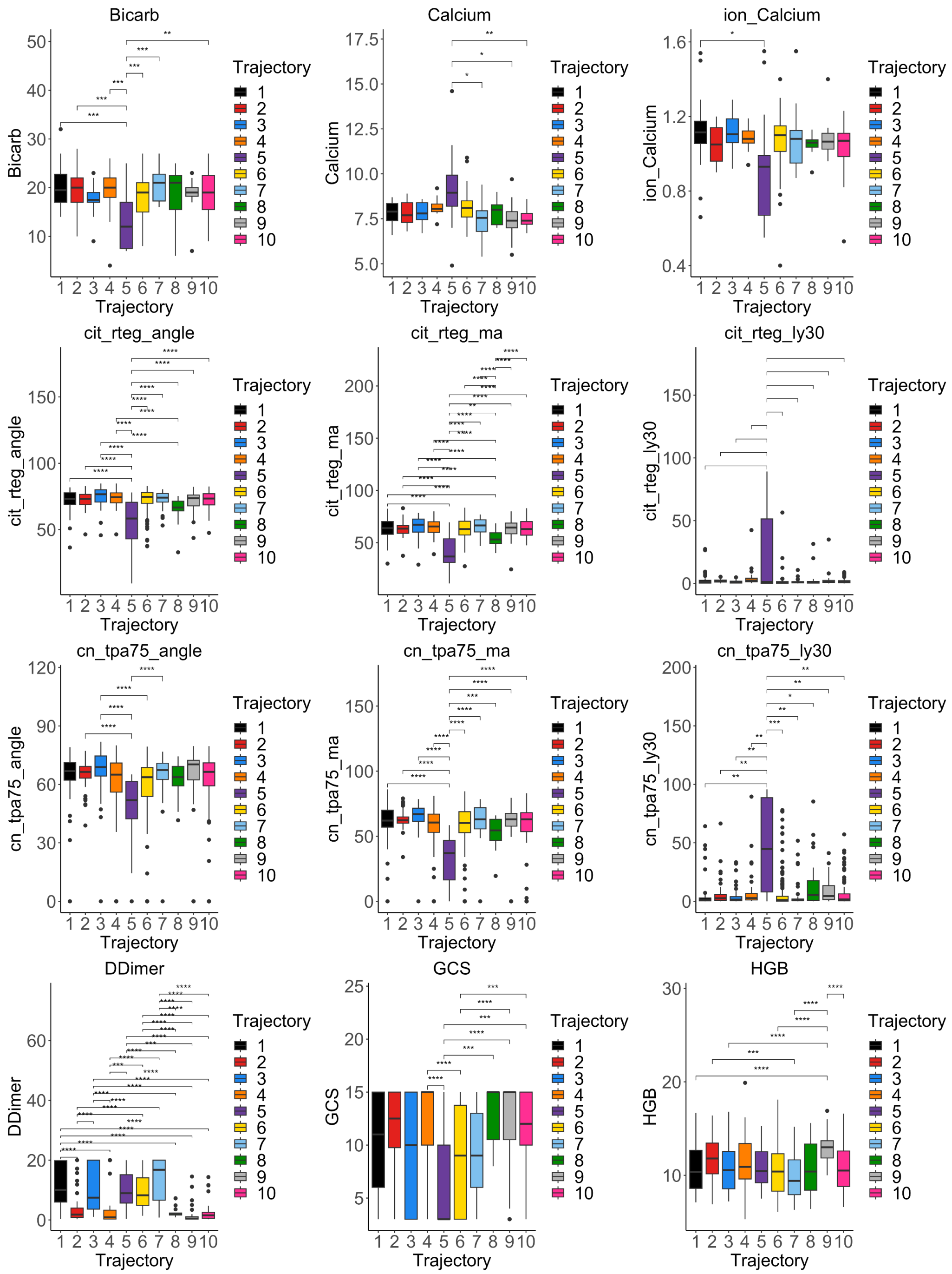

**Supplemental Figure 24 (Figure S24):** Boxplots, proportions, and line charts showing trends and statistical comparisons (ANOVA) of clinical measurements per omics trajectories.

#### Supplementary FIGURE 25

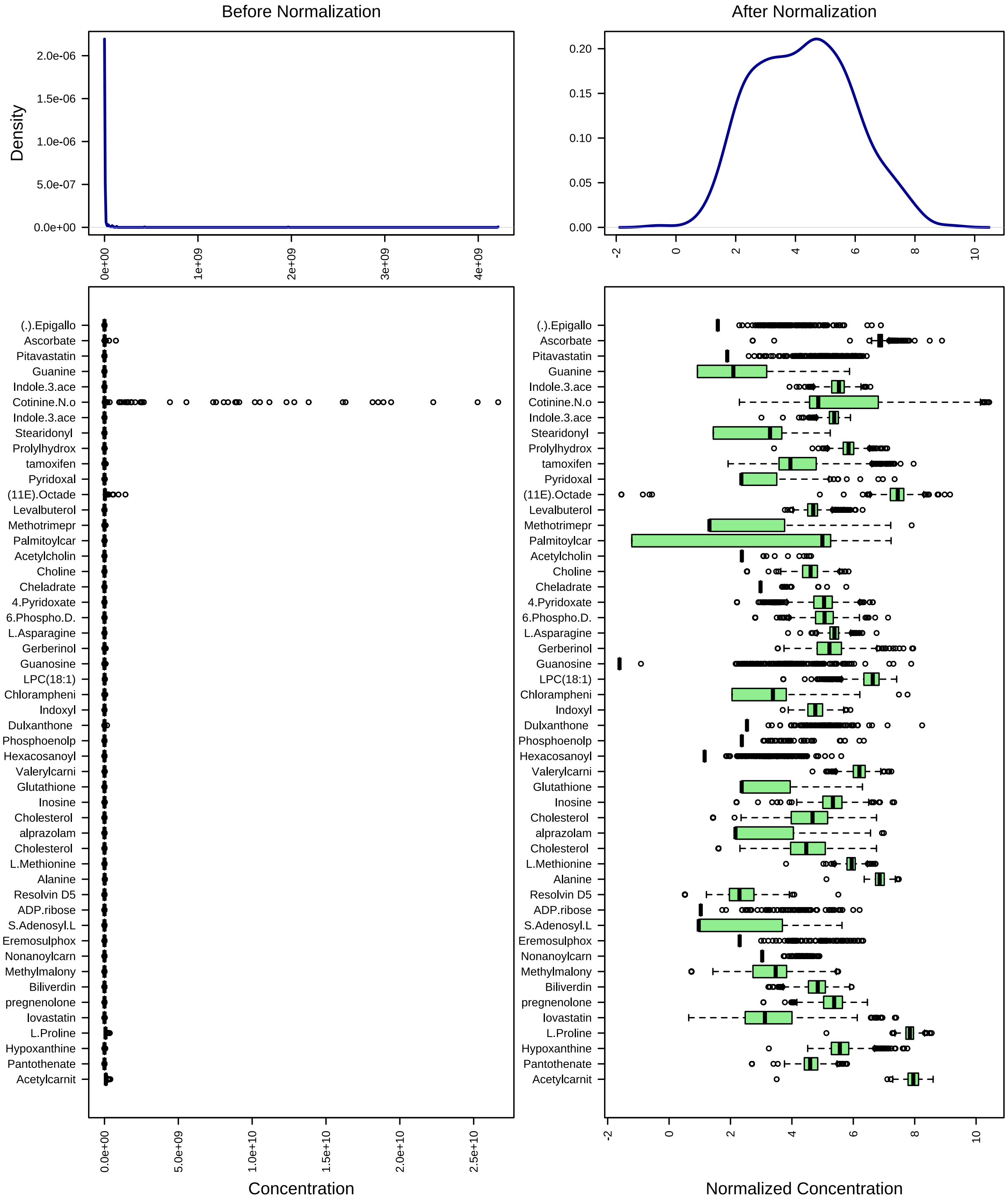

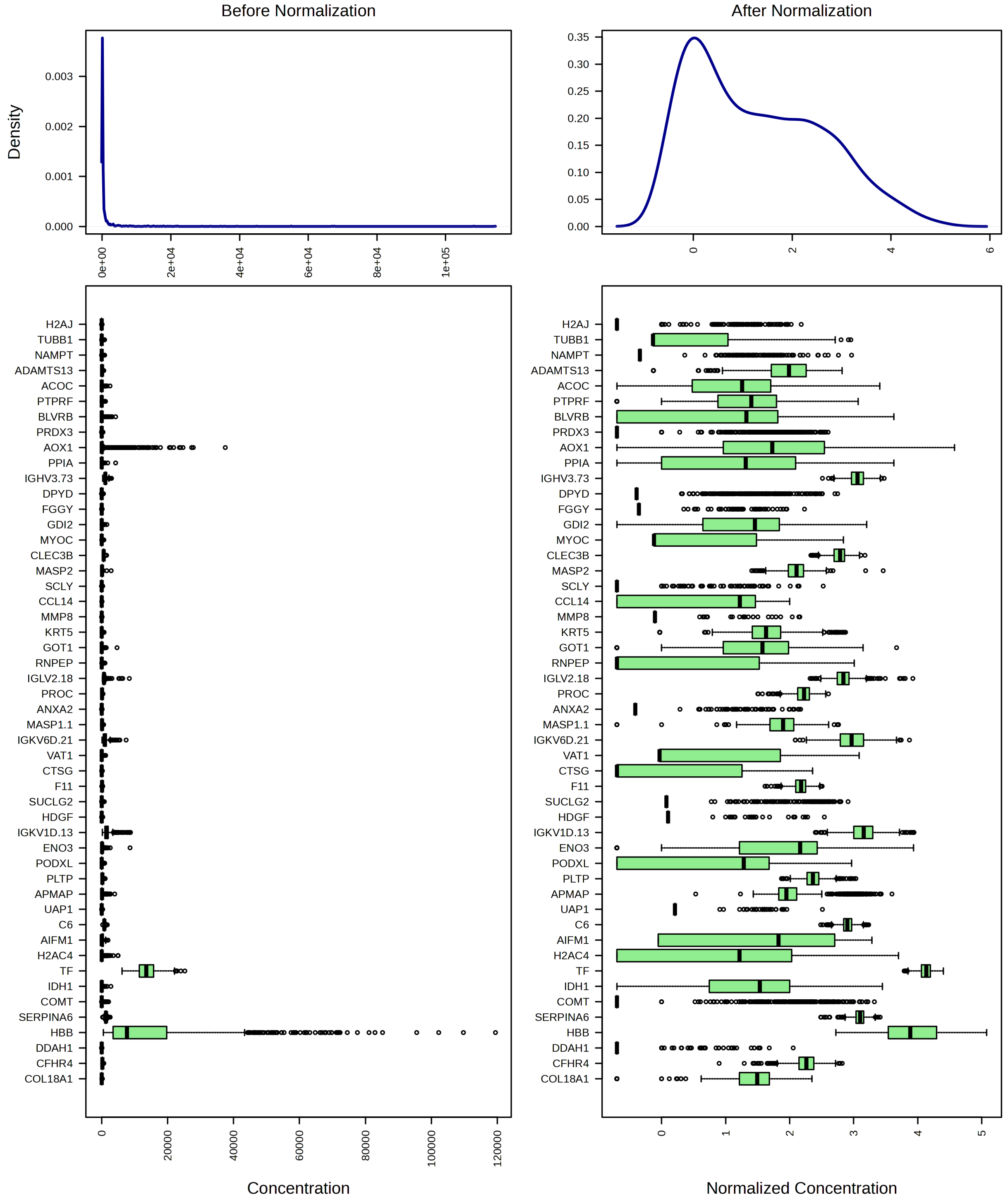

**Supplemental Figure 25 (Figure S25):** Histogram of metabolomic (upper plot) and proteomic (lower plot) data distribution before and after log2 normalization (upper
